# Prioritizing Neuroactive Ligands Using Motif-Guided Virtual Discovery and Zebrafish Profiling

**DOI:** 10.64898/2026.01.15.699747

**Authors:** Ari B. Ginsparg, Jaqueline A. Martinez, Ishaan Patel, Angélique Buton, Laura T. Lee, Raheel Sarwar, Rocco Moretti, Stephanie Puig, Summer B. Thyme

**Affiliations:** Department of Neurobiology, The University of Alabama at Birmingham Heersink School of Medicine, Birmingham, Alabama 35294, USA; Department of Biochemistry and Molecular Biotechnology, UMass Chan Medical School, Worcester, Massachusetts 01605, USA; Department of Computational Biology, University of Pennsylvania, Philadelphia, Pennsylvania, 19104; Department of Psychiatry and Behavioral Sciences, UMass Chan Medical School, Worcester, Massachusetts 01605, USA; Department of Chemistry, Vanderbilt University, Nashville, Tennessee 37240, USA

## Abstract

Virtual screening of ultra-large chemical libraries is a highly effective strategy for early-stage drug discovery. However, these pipelines often yield thousands of molecules that pass computational filters, and in silico-derived interaction energies do not consistently predict experimental efficacy. Furthermore, many high-affinity hits do not necessarily function effectively in an organism with tissues, barriers, and extensive off-target possibilities. A major hurdle in drug discovery is the prioritization of top candidates for rodent testing. Here, we introduce Rosetta Engine for Anchoring Ligands with a Motif (“REAL-M”), a novel computational screening algorithm that uses structural interaction data from the Protein Data Bank (PDB) to guide ligand placement and selection. Using the hypocretin receptor as a test case for this computational pipeline, 28 of 30 predicted antagonists significantly blocked binding of the cognate peptide agonist in a PRESTO-Tango cell-based reporter assay, including six chemically diverse molecules with comparable efficacy to preexisting antagonists. Three of the six molecules significantly mitigated hypocretin-induced larval zebrafish hyperactivity. Secondary testing with a zebrafish *hcrtr2* null mutant ensured that behavioral phenotypes were not due to off-target interactions, which we did observe with preexisting antagonists. This pipeline is readily adaptable to the thousands of zebrafish proteins with highly conserved binding pockets.

## INTRODUCTION

Advances in computational power and chemical synthesis have revolutionized in silico drug discovery. Billions of theoretically synthesizable molecules can be screened before purchasing through services like Enamine^1^ or the Pan-Canadian Chemical Library^2^. Docking using Enamine libraries has yielded high-affinity inhibitors for numerous targets (e.g., cannabinoid receptors^3^ and melatonin receptors^4^). Recently, the Enamine REadily AccessibLe (REAL) Space library has dramatically expanded to reach 76.9B in 2025, and new ways of effectively traversing this space are needed. With notable exceptions^3^, most successful discovery campaigns used conventional docking approaches.

An inherent challenge of docking is exclusive reliance on the energy function to sample ligand placements and evaluate binding. Energy functions must balance speed and accuracy, resulting in only a loose correlation between predicted and experimental affinity^5^. Predicted complexes often contain unrealistic interactions, and visual inspection of molecules is an integral part of most computational discovery pipelines^1,4,6^. Even after candidates pass in silico binding energy cutoffs and filtering, it is common to inspect as many as 5,000 molecules^7^. Better computational prioritization of molecules for experimental testing would improve objectivity and scalability by reducing the need for expert involvement.

Once high-affinity ligands for a target are identified, the next bottleneck for prospective drugs is efficacy and safety in an animal model. This step is critical to exclude off-target effects in unexpected cell types as well as assess molecules’ ability to pass tight-junction barriers. Although new alternatives such as tissue chips are emerging for preclinical drug testing^8^, rodent studies remain the field standard^9^. Testing a sufficient number of animals, however, comes with high costs and effort for characterizing a single drug^10^. Zebrafish are an emerging alternative vertebrate model for drug testing. More than 80% of proteins associated with human disease are conserved in zebrafish^11^, genetic models of disease are easily produced and often accurately recapitulate human pathophysiology, and multiple molecules discovered using zebrafish are in clinical trials^12^. Since the first zebrafish small molecule screen in 2000^13^, there have been over one hundred such studies with readouts of embryonic morphology, transgenic reporters for signaling pathways, behavior, and more^14,15^. Their low cost and scalability, combined with features such as an intact blood-brain barrier at larval stages^16^, make zebrafish a promising platform for pre-screening candidates from computational discovery campaigns.

Here, we present a new computational drug discovery approach and demonstrate the value of zebrafish for defining on- and off-target interactions of hits. Built within the Rosetta modeling software suite^17^, this method leverages Rosetta’s established energy function to optimize and evaluate final ligand placements^18^. Key advances include an ultrafast grid-based clash check for speed and the use of structural information from the PDB to guide the placement and evaluation of molecules, minimizing reliance on the energy function. Using resources from the Open Science Grid (OSG) consortium^19–22^ to screen sections of Enamine chemical space, this new approach was applied successfully to the conserved hypocretin receptor, which is involved in sleep regulation. Effective antagonists were identified using a cell-based PRESTO-Tango GPCR activity assay^23^, and larval zebrafish behavioral testing confirmed their function. This work establishes a blueprint for discovering ligands across the many diverse proteins shared by zebrafish and humans, uniting high-throughput computation with rapid in vivo validation.

## RESULTS

### REAL-M: a motif-guided screening approach

At the core of the new computational approach (**Fig. 1A, Supplementary Methods**) is a virtual library of unique protein-ligand interaction geometries, referred to as “motifs”^24^ (**Supplementary Table 1**). Motifs consist of six non-hydrogen atoms: three from an amino acid and three from a ligand. Each motif retains the bonding of motif atoms and their spatial relationships. The REAL-M algorithm uses motifs to place ligands in a target binding pocket with respect to a pre-selected anchor residue of the matching amino acid type. Following the motif-based docking search, predicted complexes that pass energy score cutoffs are filtered for the presence of additional motif interactions. This approach biases interfaces toward native-like contacts, minimizing reliance on the energy function.

**Figure 1.**
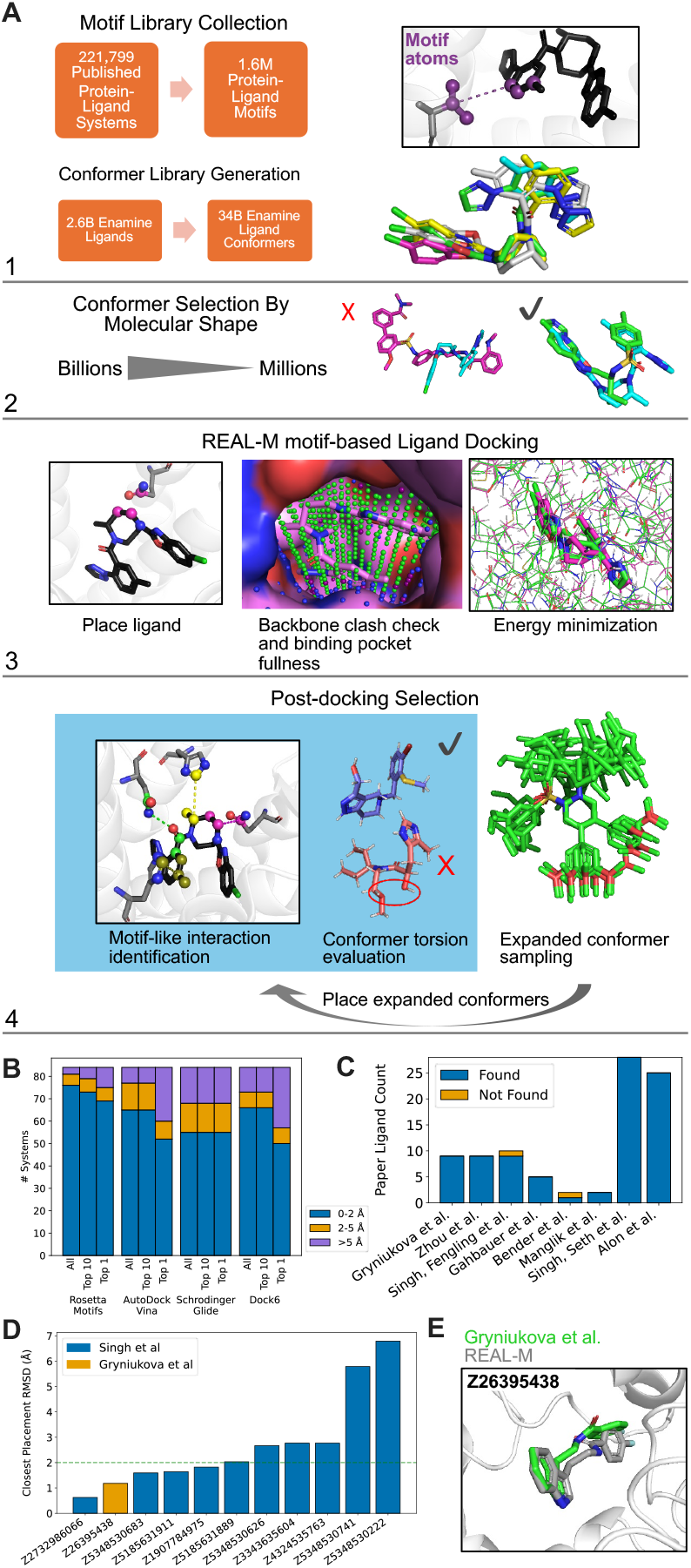
REAL-M approach to screening Enamine chemical space. (**A**) Steps of the computational pipeline. 1: Preparation of motif library and conformer library. 2: Alignment and comparison of shapes against a target ligand shape using VAMS ShapeDB. 3: REAL-M search. 4: Refinement of filter-passing molecules from step 3 by screening for motif-like interactions and strain and expanding conformers from approximately 15 to up to 250, followed by re-placing them using the step 3 process and re-evaluation. (**B**) Cross-software comparison of the recovery of 84 protein-ligand systems from the DUD-E library. Stacked bar plots representing the closest placement from all passing attempted placements, the top 10 placements by free energy, and the single top free energy placement. (**C**) Recovery of hits from previous computational docking campaigns. Ligands from the respective publication were combined with up to 10,000 ligands of similar molecular weight. Ligand placements were evaluated for free energy, total interaction count, and ratio of interactions that resembled motifs. Ligands were considered found if they had at least six motif-like interactions with receptor residues and were either present within the top 10% of free energy scores with at least 25% of binding pocket interactions resembled motifs or if at least 50% of binding pocket interactions resembled motifs. (**D**) Closest RMSD between published and REAL-M predictions for 11 ligands from two publications. The green horizontal line indicates placement similarity of < 2 Å. (**E**) REAL-M-predicted placement of compound Z26395438 (grey) against the published placement by Gryniukova et al., which was docked into in PDB 4ZZI (green). REAL-M used a motif derived from PDB 3M9F.

An effective drug discovery strategy minimizes the use of computational resources while screening large compound libraries. Considering different ligand conformations further compounds the challenge. Here, we used Conformator software^25^ to expand a set of 2.6 billion virtual molecules from the Enamine REAL collection to 34.6 billion conformers. Even if a docking algorithm assessed each molecule in a few seconds, it would take hundreds of years on a single computer to traverse these billions of molecules. To prioritize this conformer library for our screens, we filtered by shape, taking advantage of the previously published open-source software VAMS^26^ – Volumetric Aligned Molecular Shapes. Importantly, any shape, such as a known ligand or the inverse volume of the target binding pocket, can be an input to VAMS, which compares millions of shapes in a fraction of a second and scores their similarity. Within the motif-based protocol, the final stage of high-resolution docking and minimization of free energy is the most resource-intensive step (**Supplementary Fig. 1A**). To improve overall performance, we implemented multiple stepwise filters. First, and most effective, an ultra-fast, grid-based check for clashing of placements with backbone atoms was designed. For this method, the protein and ligand are represented as 3-dimensional grids of zeros and ones with a resolution of 1 Å, and shared occupancy in the same grid space triggers abandonment of the placement. In addition to removing clashes, the process is used to assess how completely the ligand can fill the binding pocket, eliminating those that are small and unlikely to make sufficient productive contacts. Then, attractive and repulsive energies are sequentially collected to form filters preventing molecules from entering high-resolution docking and free energy evaluation. Molecules are ultimately selected by assessing the presence of motifs, ligand conformational strain, and binding energy, and re-docking those that pass filters with an expanded conformer set of up to 250 per molecule.

To evaluate the performance of REAL-M, we compared its ability to recover known complexes to other commonly used programs for virtual drug discovery. For a set of 84 protein-ligand systems from the Directory of Useful Decoys Enhanced (DUD-E)^27^, low-energy placements were consistently comparable to or more accurate than AutoDock Vina^28^, Schrödinger-Maestro Glide^29^, DOCK 6^30^, and AlphaFold 3^31^ (**Fig. 1B, Supplementary Fig. 1B, Supplementary Fig. 1C**). Although AlphaFold3 can accurately model individual protein-ligand complexes, its computational cost currently precludes application at the scale required for large-scale screening, and it does not outperform docking methods on larger datasets^32,33^. Motif-based docking was also able to rediscover Enamine ligands selected by other software^6,34–40^. Successful molecules were combined with a random set of 10,000 similarly sized Enamine ligands for the motif-based search, and 34 out of 36 ligands were among the top hits when applying our standard selection criteria for prioritizing REAL-M outcomes (**Fig. 1C, Fig. 1D, Fig. 1E, Supplementary Fig. 2**).

**Figure 2.**
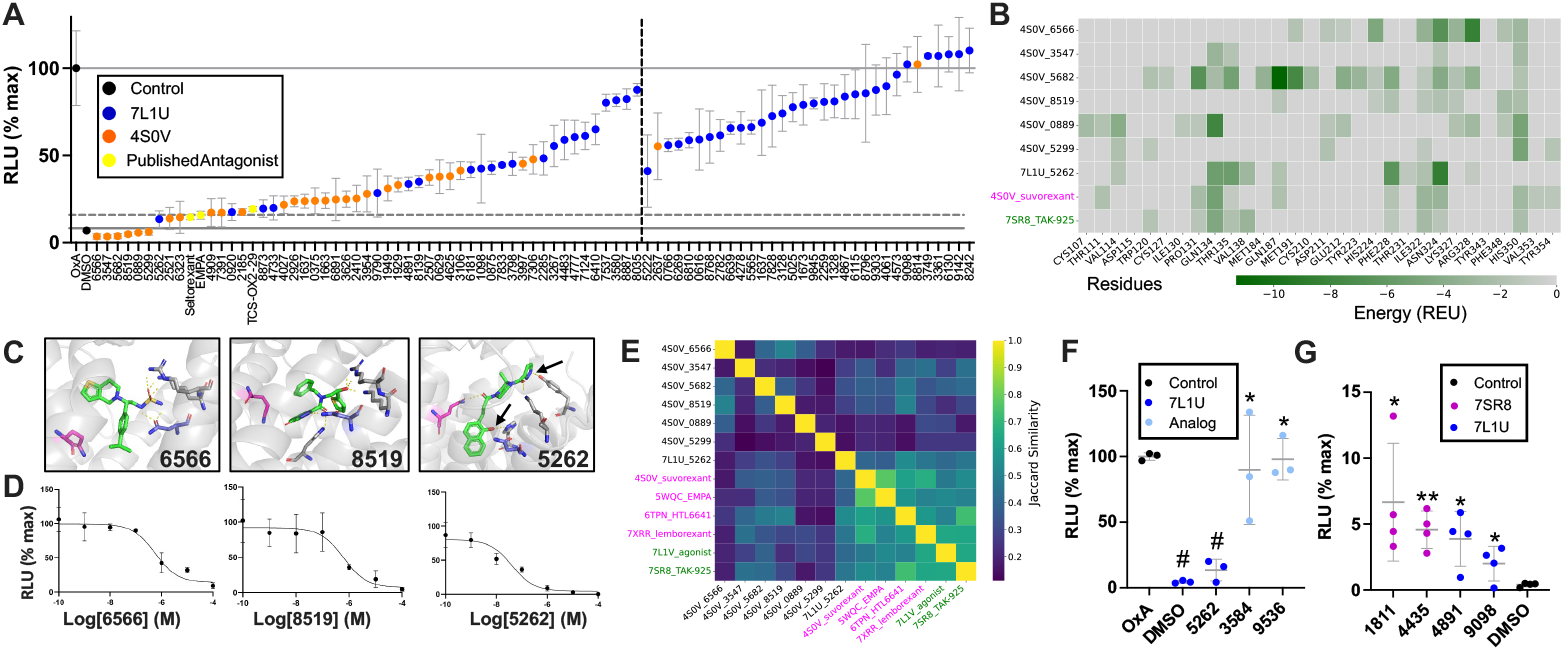
Modulators of HCRTR2 Identified by REAL-M. (**A**) PRESTO-Tango GPCR assay results of HCRTR2 antagonists. Relative luminescence values were normalized against a positive control of OxA. Drugs are separated by a vertical dashed line, with samples with significant differences from the OxA control (p < 0.05) on the left and non-significant samples on the right. Significance determined using a one-tailed Welch’s t-test (**B**) Rosetta-predicted interaction energies (Rosetta Energy Units, REU) for contacts in the binding pocket of the seven strongest antagonists (black), suvorexant (magenta), and TAK-925 (green). (**C**) Binding mode of three antagonists with GLN134 in magenta and ASN324 blue. Arrows on 5262 indicate interactions that were eliminated with analogs. Pymol sessions are available on the linked Zenodo repository. (**D**) PRESTO-Tango dose-response curves for corresponding antagonists 6566 (*IC*_50_ = 5.78e-7 M), 8519 (*IC*_50_ = 6.13e-7 M), and 5262 (*IC*_50_ = 4.811e-8 M). (**E**) Jaccard similarity of residue interaction profiles between 7 strongest experimental antagonists (black), published antagonists (magenta), and published agonists (green). (**F**) PRESTO-Tango GPCR assay results for experimental antagonist 5262 and two analog drugs with modified functional groups to inhibit predicted residue interactions. ^*^ = p < 0.05 for analog vs 5262, # = p < 0.01 for 5262 and DMSO vs OxA (one-tailed Welch’s t-test). (**G**) Presto Tango GPCR assay results for four weak HCRTR2 agonists at 1e-4 M with percentage RLU compared to TAK-861 at 1e-5 M. Groups significantly different to DMSO using a one-tailed Welch’s t-test are labeled with ^*^ (p < 0.05) or ^**^ (p < 0.01).

### Screening for modulators of the hypocretin receptor

The hypocretin/orexin receptor HCRTR2 was selected as the test case for REAL-M. This receptor is structurally and functionally conserved between zebrafish and human, and inactive and active structures have been characterized^41,42^ (**Supplementary Fig. 3**). There are both antagonists and agonists in clinical trials or approved for sleep disorders^43,44^. Active states of the receptor revealed that GLN134 is flipped upwards (**Supplementary Fig. 3**), a shift thought to be important for receptor activation^42,45^. In these activated structures, this glutamine residue interacts with the hypocretin peptide (PDB 7L1U^42^) and a chemical agonist (PDB 7SR8^45^). We ran REAL-M with the inactive (4S0V) and active states (7L1U), using ASN324 as the inactive and GLN134 as the active anchor. The inputs for VAMS shape selection were truncated hypocretin peptides for 7L1U and three structurally characterized antagonists for 4S0V (**Supplementary Table 2, Supplementary Table 3**). For each input PDB, 12 million conformers were prioritized from the 34.6 billion for the motif-based search. The VAMS selection with the hypocretin peptides yielded substantial convergence in hits, whereas each antagonist yielded an almost exclusively unique set of approximately 4 million conformers.

**Figure 3.**
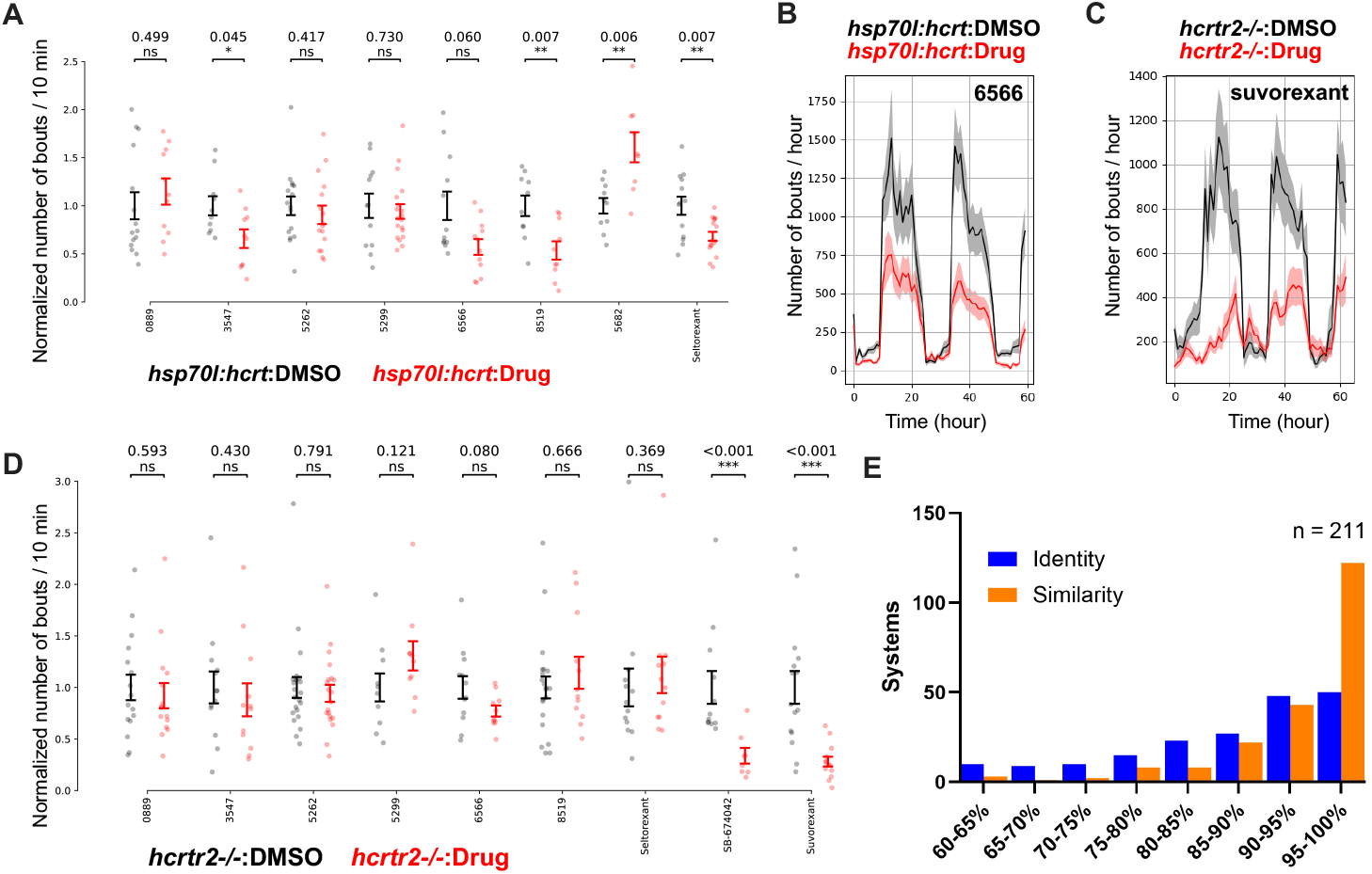
Zebrafish-based assessment of HCRTR2 antagonists. (**A**) Summary of zebrafish activity on the first experiment day (9:00-23:00, 5 dpf) following heat shock, comparing drug-treated (20 μM) to DMSO-treated larvae. The number of bouts was normalized against the average bout count from the control group per experiment to between experiments. Although non-significant in this summary plot, the 6566 compound displays significant antagonism when assessed with a linear mixed model (**Supplementary Fig 8**). (**B**) Example multi-day activity profile of larval zebrafish following heat shock for a functional antagonist. The zebrafish experience heat shock on the afternoon of 4 dpf and the activity tracking begins that night. This timecourse includes the same day 1 data as is summarized per fish in panel A. (**C**) Example multi-day activity profile of *hcrtr2* mutant larval zebrafish, comparing drug-treated (5 μM, as suvorexant is lethal at higher doses) to DMSO-treated. (**D**) Summary of *hcrtr2* mutant larval zebrafish activity on the first experiment day (9:00-23:00, 5 dpf), comparing drug (20 μM, except suvorexant, which was 5 μM) to DMSO. For improved visualization, data for 5682 is available only in **Supplementary Fig. 8**. (**E**) Percent binding pocket residue identity (blue) and similarity (orange) determined for 210 GPCRs. Human and zebrafish systems were aligned against each other, and the closest residue, if any, within 2 angstroms was considered for comparison. If no residue was found within two angstroms, the residue comparison was not accounted for in the identity or similarity determination.

Placements from the motif-based ligand search were filtered and then refined to achieve the final set of molecules for visual inspection and synthesis. Filters for predicted binding energy and motif-like interactions prioritized approximately 15k ligands for downstream refinement (**Supplementary Fig. 4, Supplementary Table 4**). As subtle differences in the conformers and their placement can have a large effect on binding energy, an expanded set of conformers for each molecule was re-run through placement and filtering to better distinguish between top candidates. After final filtering, 327 placements representing 162 unique potential binders of the active state and 461 placements representing 137 unique potential binders of the inactive state were selected for manual review for favorable binding modes (**Supplementary Fig. 4, Supplementary Table 4**). Post-review, 95 molecules were selected from this already highly enriched set for synthesis by Enamine, mainly curating for redundant pharmacophores, and 82 were received and tested (**Supplementary Table 5, Supplementary Table 6, Supplementary Table 7**).

Over half of the tested molecules significantly inhibited binding of the orexin A peptide to the hypocretin receptor using the PRESTO-Tango GPCR activity assay (**Fig. 2A**). Of those modeled by docking against the 4S0V inactive state, which has GLN134 in the downward orientation, 28 of 30 were significant antagonists. Of those docked against the active state, many (21 of 52) were also effective antagonists despite being explicitly anchored to the upward-facing GLN134 in the active state (7L1U). The seven strongest inhibitors, which have an *IC*_50_ comparable to commercially available molecules (**Fig. 2B, Supplementary Fig. 5**), included one from the 7L1U screen. All seven interact with unique binding pocket residues (**Fig. 2C, Fig. 2D**) and form diverse protein-ligand complexes compared to both structurally characterized antagonists and agonists (**Fig. 2E, Supplementary Fig. 6**). We were surprised that a high-affinity antagonist (5262) was identified by docking against the active state of the receptor. To substantiate its predicted binding mode, we searched the Enamine library for ligand analogs missing atoms involved in a key interaction (**Supplementary Table 6**). Both selected analogs eliminated the ability of 5262 to block peptide binding (**Fig. 2F**). From the active state 7L1U screen, only very minimal receptor activation was observed when the molecules were separately tested for agonism (**Fig. 2G, Supplementary Fig 7**), indicating that stabilizing the upward flipped GLN134 is insufficient to induce the active conformation.

We extended agonist discovery by conducting a secondary screen with a single motif and over 2.9B molecules. This screen did not rely on VAMS to filter the library but instead achieved the necessary speed by limiting the motif library to a single interaction geometry. One chemical agonist with an available activated structure (7SR8) maintains the receptor in an activated conformation by not only binding GLN134 but also through a sulfonamide interaction that intercalates between two helices (**Supplementary Fig 7**). Screening sulfonamide-containing molecules from both the Enamine 2.6B and 64.9B libraries identified two additional weak agonists (**Fig. 2G, Supplementary Fig. 7, Supplementary Table 8**), further supporting the conclusion that the glutamine flip is necessary but insufficient for full receptor activation. Over half of the twenty molecules tested from the 64.9B library were also highly effective antagonists (**Supplementary Fig. 7**), blocking the hypocretin peptide with similar efficacy to preexisting antagonists but via a very different binding mode.

### Assessment of efficacy and toxicity in animal models

To determine whether antagonists identified by the PRESTO-Tango assay could influence animal behavior, we screened the top seven in zebrafish. Transgenic overexpression of hypocretin peptide (Hcrt) following promoter activation with heat shock leads to a significant increase in movement for multiple days^46^ (**Supplementary Fig. 8**). Three of seven (3547, 6566, 8519) mitigated the hypocretin-induced hyperactivity (**Fig. 3A, Fig. 3B**). To determine whether behavioral effects were mediated through the zebrafish hypocretin receptor *hcrtr2*, we generated a loss-of-function mutant and compared the effects of these molecules compared to DMSO in the homozygous mutant background. The homozygous mutant has no significant impacts on larval behavior (**Supplementary Fig. 9A**), as was previously observed for another mutant line^47^. We discovered, however, that the FDA-approved suvorexant and the commercially available antagonist SB-674042 caused dramatic reductions in larval movement even in *hcrtr2* mutants (**Fig. 3C, Fig. 3D, Supplementary Fig. 9B**). In contrast, most newly identified molecules produced only mild, non-significant effects, indicating that the behavioral changes in the heat shock line were specific to hypocretin signaling. The marked effects of suvorexant and SB-674042 in the mutant background suggest these compounds can have significant and behaviorally relevant off-target interactions.

## DISCUSSION

Here, we introduce a new knowledge-guided approach for virtual ligand screening and leverage the unique strengths of the zebrafish model to prioritize candidates. The REAL-M computational algorithm, built in the Rosetta software suite, combines spatial interaction constraints from published structures with a physics-based energy function. Refinement with the energy function is preceded by a rapid clash check (**Fig. 1A, Supplementary Fig. 1A**), an advance critical for achieving scalability while maintaining prediction accuracy, consequently allowing assessment of over 2.9 billion molecules in one virtual screen (**Supplementary Fig. 7**). Post-screening of predicted complexes for the presence of multiple motif interactions further nominates high-quality binding poses, mitigating the common docking challenge of thousands of hits being retained after energy filtering^5,7^. REAL-M was highly successful in computational benchmarks compared to existing software (**Fig. 1**) and in experimental antagonist identification (**Fig. 2A**). Behavioral screening in zebrafish revealed in vivo efficacy for predicted ligands (**Fig. 3**).

We successfully identified dozens of antagonists for the hypocretin receptor, using both inactive and active structures as starting points for discovery (**Fig. 2A**). The upward-facing orientation of GLN134 has been highlighted as critical for receptor activation^42,45^. Thus, we targeted this orientation by using the glutamine as a motif anchor, starting from two separate activated receptor structures (7L1U and 7SR8), but identified effective antagonists and only weak agonists (**Fig. 2G**). We expect that the binding modes of these modeled compounds are accurate and stabilize the upright GLN134, as many act as antagonists and modification of key predicted interactions eliminated binding of the 7L1U-derived molecule 5262 (**Fig. 2F**). Even though over 2.9B sulfonamide-containing molecules from the 64.9B Enamine library were screened with REAL-M, they still represent a small fraction of chemical space, reflecting the continued need for custom molecule design and synthesis. These agonists can serve as diverse starting points for further development: a structural comparison of their binding mode with that of TAK-861^43^, a potent agonist (**Fig. 2G**), would provide insight into the subtle differences beyond GLN134 that are necessary to drive activation.

Although the current implementation of REAL-M proved highly effective in identifying potent ligand binders with diverse chemotypes and binding modes (**Fig. 2, Supplementary Fig. 6**), there are several directions for continued improvement. The challenge of strong agonist identification highlighted the reliance of REAL-M on functionally relevant structures and its algorithmic limitations. The method assumes a relatively rigid receptor backbone during motif placement, and each placement occurs sequentially. Subtle structural changes can have large effects on binding energy and ligand ranking, as demonstrated by the importance of expanding the ligand conformers from 15 to 250 in the refinement round of REAL-M. This frontier could be explored by modifying the rapid clash check method and anchored placement to accommodate structural ensembles, such as those generated by molecular dynamics, instead of being limited to single protein backbones, conformers, and motifs. The current motif library is also sidechain-centric, and motif-guided contacts with backbone atoms in the receptor pocket are possible in the future. If integrated with REAL-M, using AlphaFold2 models as starting points for discovery^48^ would make over one-third of the human proteome accessible to virtual drug screening^49^.

Our work demonstrates the opportunity of incorporating zebrafish testing in drug discovery pipelines. Over 70% of genes are conserved between humans and zebrafish^11^, and the binding pockets of 165 GPCRs, which are major drug targets, are >90% similar (**Fig. 3E**). Screening molecules in zebrafish serves to nominate top candidates and uncover off-target actions and toxicity through mutant-based comparisons (**Supplementary Fig. 9**). The zebrafish platform is ideal for screening molecules from small-scale synthesis, as each behavior experiment typically requires a maximum of 300 micrograms. Mutants can be readily generated, and the highly sensitive phenotyping approaches accessible with this model, such as brain activity mapping^50^, could illuminate even subtle unintended effects. Regardless of phenotype face validity, the zebrafish serves as an effective screening platform when the receptors are conserved or if a humanized model can be generated. Although pharmacokinetic and pharmacodynamic properties differ between species, a molecule that produces robust, on-target effects in zebrafish represents an ideal candidate for further development.

## MATERIALS AND METHODS

### Zebrafish husbandry

Zebrafish experiments were approved by the UAB and UMass Chan Institutional Animal Care and Use Committees (IACUC protocols 22155, 21744, and 202300000053). The *hcrtr2*^uab480^ mutant was generated in an Ekkwill-based strain using CRISPR/Cas9 as previously described^50^ and is available from ZIRC. The *Tg (hsp70l:hcrt)* line was previously described^46^. Line generation and genotyping details are available in **Supplementary Table 9**. Both adults and larvae were maintained on a 14 h/10 h light/dark cycle at 28°C, and experimental larvae were grown at a density of less than 160 per 150 mm petri dish. Larvae without inflated swim bladders were excluded from experiments. Each experiment used larvae from a single parental pair, control animals were always siblings, and all larvae were genotyped after the behavioral run was completed.

### Motif library and conformer library preparation

Motifs were extracted from 221,799 protein-ligand structures. Duplicate motifs were eliminated based on atom types and spatial placement, and the composition of the final library is depicted in Supplementary Table 1. A set of 2.6 billion synthesizable ligands was downloaded from REAL Enamine catalog in .sdf file format, and Conformator^25^ was used to generate up to 15 energetically favorable conformers of each ligand. Rosetta’s molfile_to_params.py script was used to convert each conformer into a Rosetta-readable .params format. VAMS ShapeDB^26^ was then used to generate a database of molecular shapes to compare for all conformers within the conformer library against a target molecular shape. The entire conformer library was aligned and stored in a database, which was queried with target molecules to find similar shapes. All conformers were ranked via a similarity score compared to the input molecule. For 4S0V-based discovery, the conformer library was compared against published antagonists EMPA, HTL6641, and Lemborexant (**Supplementary Table 2**). For 7L1U-based discovery, the conformer library was compared against truncations of orexin A (**Supplementary Table 3**). Although molecules were used as input to VAMS, a grid that represents the complement of the binding pocket can also be used. For 7SR8-based discovery of sulfonamide, shape selection was not used, and the conformer library was instead limited to molecules that contain the sulfonamide group and the motif library to only a single motif matching the TAK-925 interaction. Open Babel^51^ was used for molecule file format conversions and splitting of multiple molecule files into single molecule files. RDKit^52^ was used for molecule file format conversions, calculation of molecular properties (SMILES strings, molecular weight, chemical formula), detection of molecular fragments, and calculation of intermolecular Tanimoto similarity.

### REAL-M discovery search

Conformers were placed in the binding pocket of HCRTR2. For receptor PDB files, the receptor chain was extracted from the full file, and all ligands and waters were removed. The 4S0V screen used ASN324 as the anchor residue for motif placement, as it is a central residue with hydrogen bonding potential. The 7L1U and 7SR8 screens used the highly shifted GLN134 residue (**Supplementary Fig. 3**) as the anchor. Conformers passing score cutoffs were further screened for the presence of motifs, prioritizing those with a high proportion of these native-like interactions compared to total interactions. Those passing metrics of free energy, proportion of real motif-like interactions, and interactions with residues of interest, were further refined by rerunning the search process with an expanded set of 250 conformers. Placed ligands were screened again with the same criteria, along with a strain score predicted with TLDR Strain^53^ and recovery of the placement with AutoDock Vina^28^. Passing ligand placements were visually screened for a final evaluation and selection of ligands to be synthesized by Enamine. All Pymol sessions for ordered molecules are available at the following Zenodo repository: https://zenodo.org/records/17730882?token=eyJhbGciOiJIUzUxMiJ9.eyJpZCI6ImI1ZWY4ZTc0LTUxOGYtNGUxYS04MzU2LTJjZWRmOTEzOTA4YSIsImRhdGEiOnt9LCJyYW5kb20iOiI2NjA1N2M3M2QxMjhjZmJkZTBkODVmZDY0ODRlNTNkMSJ9.6HZWKvnmzOZ9WkTcGcmujO55iFpd8Jg3fzUSougc3Ew5sXClVvwmH5DlaxAwyz0hu50hMVY4p-gH1EMTMXnAxw

### Recovery benchmarks

To benchmark REAL-M against existing drug discovery software, the recovery of native placements of known ligands from a diverse set of 84 DUD-E^27^ protein-ligand systems was compared. A protein homology query was performed on each DUD-E system, and motifs from any homologous PDB were not included in the motif library. RMSD was calculated for each placement. The smallest distance was collected for each system for all predicted placements, the top 10 systems by free energy, and the single lowest free energy placement. To determine whether REAL-M can rediscover molecules from published docking campaigns, the search process was run by combining the hit molecules with a random set of 10,000 others of similar molecular weight. We determined if the publication ligand would be identified by our pipeline if a ligand placement had at least six motif-like interactions and either was present in the top 10% of placements by free energy score with at least 25% of binding pocket interactions resembling real interactions from the PDB or if at least 50% of binding pocket interactions resembling real interactions from the PDB.

### PRESTO-Tango assay

Synthesized compounds were screened using the PRESTO-Tango^23^ GPCR activity assay for agonistic or antagonistic effects on HCRTR2. Confluent HTLA cells were passaged into white, clear-bottom 384-well plates at 5,000 cells in 50 μL DMEM media (Gibco) with 10% FBS per well. Cells were incubated for 24 hours at 37°C, followed by transfection with a cocktail of the PRESTO-Tango HCRTR2 construct (Addgene plasmid # 66399), serum-free Opti-MEM (Gibco), and FuGENE (Promega). Cells were allowed to incubate for 24 hours at 37°C. Wells were treated with experimental drug or orexin A (OxA) peptide (Phoenix Pharmaseuticals - cat# 003-30) that was initially diluted in DMSO and diluted to a working concentration with PRESTO-Tango assay buffer of 2% 1M HEPES in HBSS (Gibco). For agonist experiments, cells were treated with 20 μL of drug solution diluted to yield the desired final concentration. For antagonist experiments, cells were treated with 10 μL of drug solution, incubated for an hour at 37°C, and then treated with 10 μL OxA for a final concentration of 1e^-7^ M. Negative control wells had an equivalent DMSO concentration to experimental conditions. After incubation for 24 hours at 37°C, wells were treated with 5 μL of BrightGlo reagent (Promega - cat# E2610) and incubated in darkness at room temperature for 15 minutes. A back seal sticker (Perkin Elmer) was applied to the bottom of the plate to make the well bottoms opaque. All experiment wells were measured for relative luminescence with an Envision 2105 (Perkin Elmer) plate reader. The signals were normalized against the average positive control luminescence. Data was visualized and plotted, along with *EC*_50_ or *IC*_50_ for dose-response curves, using GraphPad Prism 10. The following commercially available molecules were tested in addition to those from Enamine: MK-1064 (MedChemExpress - cat # HY-19914), SB-674042 (MedChemExpress - cat # HY-10898), Seltorexant (JNJ-42847922) (MedChemExpress - cat # HY-109012), TCS-1102 (MedChemExpress - cat # HY-10900), TCS-OX2-29 (MedChemExpress - cat # HY-100452), and TAK-861 (Overporexton) (Selleckchem - cat# E4740). Suvorexant (MK-4305, Belsomra) (Cayman Chemical - cat# 22911) was toxic to the cells.

### Zebrafish behavior screen

All behavior experiments were conducted from 4-7 days post-fertilization (dpf), with single larvae in individual wells of a 96-well square-well plate. For heatshock experiments, before plating, larvae were transferred into a 50 mL conical tube with 20 mL fish water with methylene blue and submerged in a 38°C water bath for 90 minutes. The 96-well plate contained alternating columns of drug and control conditions, with 20 μM of drug (unless otherwise specified) or an equivalent concentration of DMSO in a 500 μL volume of fish water. As this is a multi-day experiment, the plates were sealed with oxygen-permeable film. For sealed experiments, we typically overfill the wells and seal the plate without air bubbles^54^; this procedure is not possible when comparing drugs to controls in the same plate. The remaining air bubble sometimes resulted in condensation, obscuring tracking, so all videos were checked manually, and larvae with poor tracking were eliminated blind to genotype. The larvae were tracked in custom-built behavior box apparatuses, and the activity was analyzed as previously described^54^. Raw data is available at the following Zenodo repository: https://zenodo.org/records/17730882?token=eyJhbGciOiJIUzUxMiJ9.eyJpZCI6ImI1ZWY4ZTc0LTUxOGYtNGUxYS04MzU2LTJjZWRmOTEzOTA4YSIsImRhdGEiOnt9LCJyYW5kb20iOiI2NjA1N2M3M2QxMjhjZmJkZTBkODVmZDY0ODRlNTNkMSJ9.6HZWKvnmzOZ9WkTcGcmujO55iFpd8Jg3fzUSougc3Ew5sXClVvwmH5DlaxAwyz0hu50hMVY4p-gH1EMTMXnAxw

## Acknowledgments

This work was supported by NIH grant DP2 NS132107, a Pew Biomedical Scholars award, Klingenstein-Simons Fellowship award in Neuroscience, and Alkermes Pathways award to SBT. The research was completed using services provided by the OSG Consortium^19–22^, which is supported by the National Science Foundation awards #2030508 and #2323298. We thank the UAB and UMass Chan fish facility staff and the Research Computing teams at UAB and UMass Chan, particularly Prema Soundararajan, for supporting this study. The HTLA cells were a gift from Bryan Roth. The HCTR2-Tango construct was a gift from Bryan Roth (Addgene plasmid # 66399 ; http://n2t.net/addgene:66399 ; RRID:Addgene_66399). The *Tg (hsp70l:hcrt)* line was a gift from David Prober. We thank Matthew D. Smith for early contributions to the ligand motif search algorithm, Enamine representatives Roman Hromov and Viktoriia Kharchenko for assistance, and Faith Williams and Lori Ginsparg for labeling support.

## Author contributions

SBT and AG conceived of the study and with contributions from all authors. SBT, AG, AB, and SP designed experiments and analyzed data. JAM, AG, and AB performed cell and animal experiments. AG developed the computational algorithms. IP, LTL, RH, SBT, and RM contributed to developing computational tools and completing computational experiments.

