## Supplementary Information for "Prioritizing Neuroactive Ligands Using Motif-Guided Virtual Discovery and Zebrafish Profiling"

For Ari B. Ginsparg *et al.*

**This PDF file includes:**

Supplementary Methods
Supplementary Figures 1-9
Supplementary Tables 1-9

### Supplementary Methods

All paths are from GitHub repositories. Unless otherwise specified, the full path is https://github.com/toadlover/ginsparg_thymelab_thesis with relevant subdirectories listed.

The following workflow describes REAL-M. The main steps (grey) of the pipeline are listed in descending order. Actions included in each main step are listed to the right in the order that they occur, with blue designating processing steps and orange filtering. If a placed ligand does not pass a filtering step, it is immediately discarded, and the next placement is made.

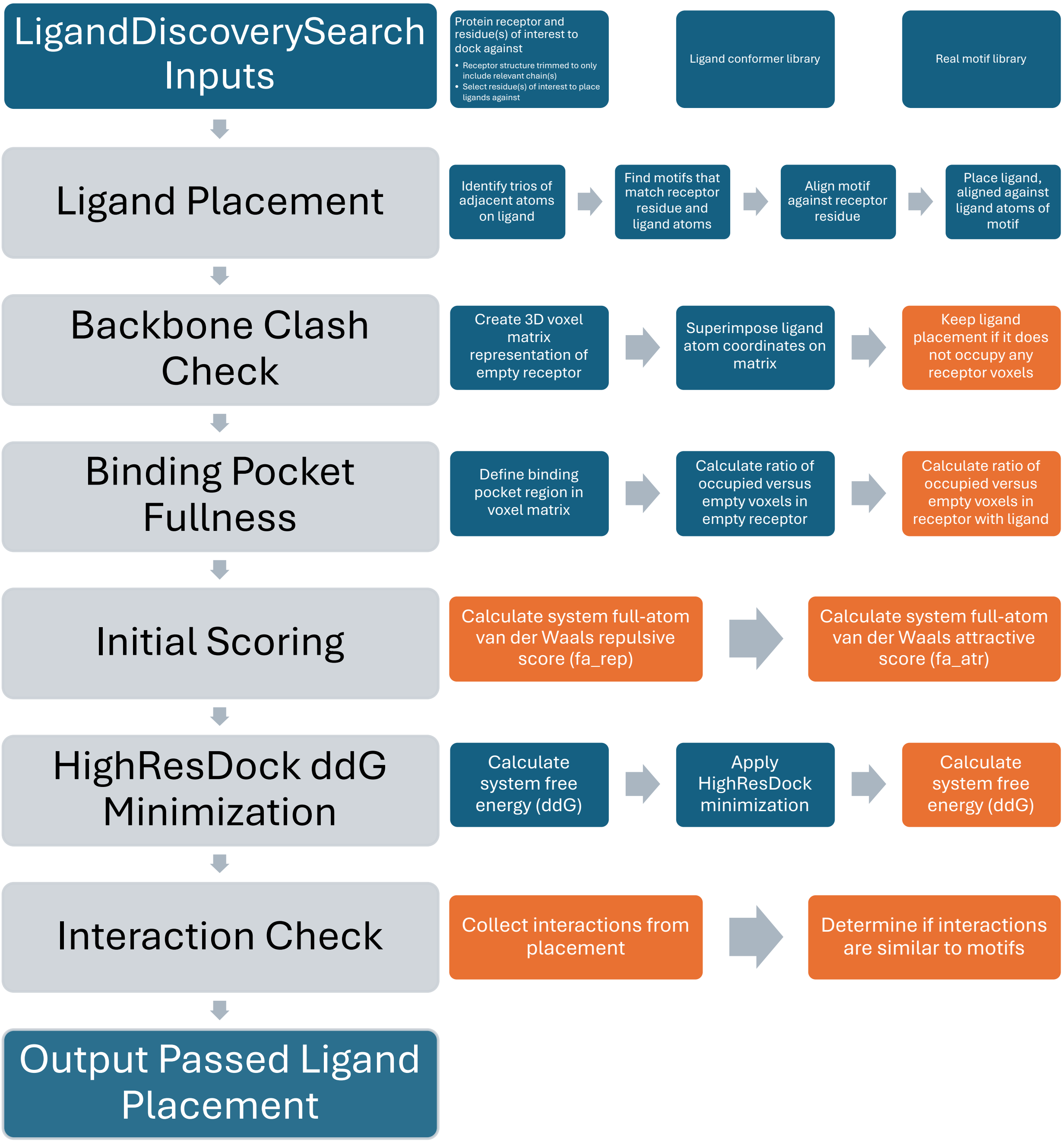

Docker Container and Singularity Image Creation

To promote reproducibility of software environments, Singularity images were created for the Conformator, VAMS ShapeDB, and Rosetta programs. Instructions to recreate these environments are located within ginsparg_thymelab_thesis/dockerfiles/ in respective folders for Conformator, Rosetta, and ShapeDB. Within each folder is a file named "Dockerfile", which gives Docker the instructions to make a container. The container is converted into an image by Singularity. The script build_commands.sh was used to automate the generation of images from respective Dockerfiles. This pipeline used Docker version 24.0.5 and Singularity version 4.1.0.

Version of Rosetta repository used for container generation: https://github.com/RosettaCommons/rosetta/commit/a5347d89a92c63988ece5a79da78e6fba666785b

Example command for Singularity image generation:

```
#Note: This script should be run at the location of the Dockerfile
#This example is to build the Rosetta container
#This results in a Singularity image named rosetta.sif
./build_commands.sh rosetta
```

Steps in build_commands.sh to create the Docker container and convert it to a Singularity image:

```
#Using docker version 24.0.5 and singularity version 4.1.0
#$1 is the name of the container that you want, which will pass down all steps

#build container:
sudo docker build - < Dockerfile -t $1 --progress=plain

#get image ID in Docker:
image_id=$(sudo docker images | grep $1 | awk '{print $3}')

#write image to a tar file that singularity can use:
sudo docker save $image_id -o $1.tar

#build Singularity image from Docker archive
sudo singularity build $1.sif docker-archive://$1.tar
```

A singularity image contains a replicable instance of software that can be repeatedly accessed using Singularity's shell command:

```
#Example to activate the Rosetta Singularity image
#The image can be exited using ctrl+D
singularity shell rosetta.sif
```

Motif Library Generation

***Protein Data Bank File Collection***

A total of 221,799 structures containing small molecules were obtained from the RCSB PDB in early 2023. Files were initially collected as a list of comma-separated PDB codes (/ligand_motifs_collection/pdb_list_all_newline_separated.txt). For easier processing, file lists were separated by PDB codes starting with 1, 2, 3, 4, 5, 6, and 7-9. RCSB's batch_download.sh script was used to pull PDB files from the file list:

```
./batch_download.sh -f pdb_list_789.txt -p
```

#### *Protein Data Bank File Preparation and Motif Collection*

The following workflow describes the motif library collection. The main steps (grey) of the pipeline are listed in descending order. Actions included in each main step are listed to the right in the order that they occur, with blue designating processing steps and orange filtering. Steps with a purple background are processes of the Rosetta remove_duplicate_motifs app.

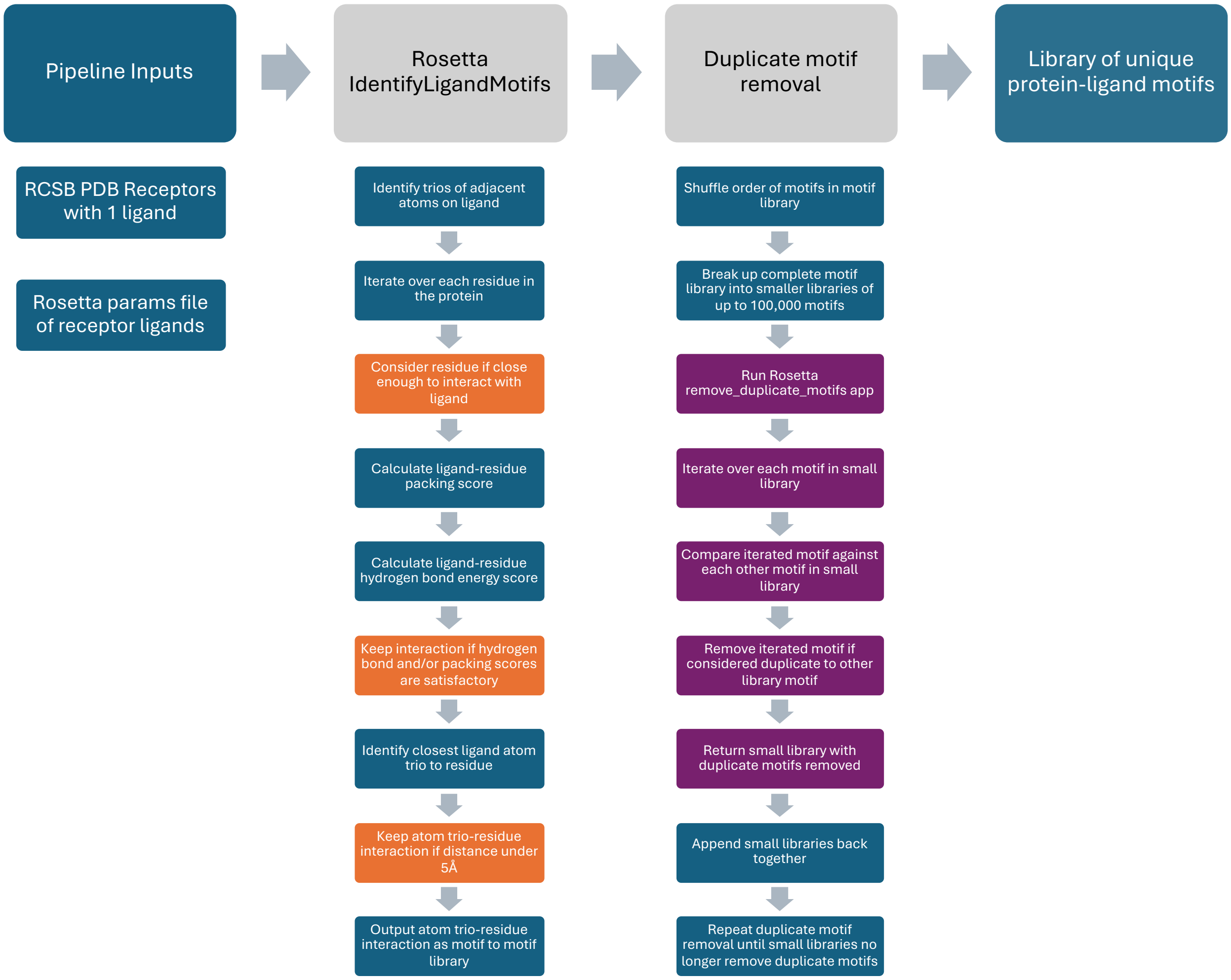


The script motif_database_generator_updated_1_27_23.py was run on a directory of protein-ligand PDB files to prepare the files and collect motifs. It requires no input arguments and processes all PDB files in the running directory. The script first creates copies of each PDB that contain a single ligand (e.g., 7BYU_1PG.pdb and 7BYU_EDO.pdb). The input Rosetta argument files and ligand params file, required for all small molecules, are also generated by the script prior to using Rosetta for motif collection.

Motifs are collected through the Rosetta identify_ligand_motifs app, which is also instantiated by the Python script. First, amino acid residues that are within a distance of 1.5 times the sum of the ligand and residue neighbor radii (NBR radius) are selected for potential motifs. Then, the ligand-residue side chain packing and hydrogen bond scores are derived, and the interaction was considered if the combined score was smaller than -1. The three closest adjacent atoms in the ligand were selected to represent the motif and was recorded if the closest atom was within 5 Å of the residue. Only one ligand-residue motif can be derived per residue.

The identify_ligand_motifs app can also be run directly without starting from the Python script:

```
#allow ligands in system to exist and not have to worry about having params files
of them included
-ignore_unrecognized_res

#Inputted PDB file
-s 7BYU_1PG.pdb

#minimum cutoff for hydrogen bond score
-hb_score_cutoff -0.3

#minimum cutoff for packing score
-pack_score_cutoff -0.5

#optional flag to not print out generated motifs as pdb files
#Default value is true, and visualization can be good for debugging
#For large scale runs, it is worth including this flag and keeping it set as false
-output_motifs_as_pdb false
```

identify_ligand_motifs example command:

```
/rosetta/source/bin/identify_ligand_motifs.linuxgccrelease @7BYU_1PG_flags
```

***Protein-Ligand Duplicate Motif Removal***

Motifs from the PDB systems were compiled into a single motifs file. A pipeline was run on this file to remove motifs that were duplicates, based on the following criteria: a match of all atom types for the amino acid and ligand, a RMSD less than 0.8 Å, and a jump angle difference less than 0.3 radians.

To discover duplicate motifs, each motif in the list was compared against all others. A Slurm script, remove_duplicate_motifs_helper.job, called the Python scripts shuffle_motif_lines.py and remove_duplicate_motifs_helper.py to iteratively remove duplicate motifs from the complete list. The starting motifs list was broken into smaller segments of up to 100,000 motifs that were filtered for duplicates and then spliced back together. The order of motifs in the spliced file was randomized. This process is repeated at least five times until duplicate motifs were no longer being removed from the randomized smaller motifs lists. Variables at the start of remove_duplicate_motifs_helper.job were defined to target the starting motifs file, path to Rosetta executable for duplicate motif removal, and the number of motifs per smaller motifs file.

```
#starting motifs file
file_name=200000_sample_motifs_file.motifs
#path to the executable for remove_duplicate_motifs
path_to_executable=/rosetta/source/bin/remove_duplicate_motifs.linuxgccrelease
#number of motifs per sub-file (ideally 100k-1M)
#there is a balance between too few motifs per sub to find duplicates and the jobs
running too long (too many motifs per sub)
```

```
motifs_per_sub=100000
```

Below are example argument flags for the Rosetta remove_duplicate_motifs app.

```
#option to allow overwriting files when one currently exists
-out::overwrite true

#name of initial input motifs file that may contain duplicates
-motif_filename 5000_motifs.motifs

#Name of output motifs file with duplicates removed
-output_file 5000_motifs_duplicates_removed.motifs

#distance threshold for RMSD of atoms in angstroms
-duplicate_dist_cutoff 0.8

#angle threshold for atoms in radians
-duplicate_angle_cutoff 0.3
```

remove_duplicate_motifs example command:

```
/rosetta/source/bin/remove_duplicate_motifs.linuxgccrelease  @flags
```

Conformer Library Preparation

***Enamine Library Preparation***

A set of 2.6 billion synthesizable ligands was downloaded from the REadily AccessibLe (REAL) Enamine catalog in .sdf file format as a single file (download date ca. July 2021). To improve library handling, ligands were separated into groups of up to 50,000, referred to as “chunks”. Chunks were numerically labeled from 00000 to 53084. Each chunk was further subdivided into up to ten “sub-chunks” of 5,000 ligands for a total of 530,842 sub-chunks, each one a single file in .sdf format.

***Conformer Library Preparation***

Using Conformator, up to 15 diverse and energetically favorable conformers were generated for every ligand. Conformator needs to be activated with an active license key before being able to generate conformers. A license can be requested from the Universitat Hamburg website (https://www.zbh.uni-hamburg.de/forschung/amd/software/conformator.html). Each conformer in .sdf format was separated into an individual .sdf file using Open Babel, renamed by adding a unique number from 0-14. For each conformer, the required Rosetta ligand params file was then generated. Both steps were completed by running the script make_conformers_and_params.py on each sub-chunk.

make_conformers_and_params.py example command:

```
#split_new_named_0.sdf is a file containing 5,000 unique ligands
#activate this script from within the Conformator container
python ginsparg_thymelab_thesis/conformator/make_conformers_and_params.py
split_new_named_0.sdf (conformator_license_key)
```

Example call to Conformator within make_conformator_and_params.py:

```
#split_new_named_0.sdf is a file containing 5,000 unique ligands
#Flags used:
#-i - name of input ligand file (with path if necessary)
#-o - name of output file with all conformers of initial ligand (with path if
necessary)
#--keep3d - keep 3d representation of conformers
#--hydrogens - add/keep hydrogens for generated conformers
#-n - maximum number of conformers to keep
#-v - verbosity, set to zero to minimize output
conformator_1.2.1/conformator -i split_new_named_0.sdf -o confs.sdf --keep3d --
hydrogens -n 15 -v 0
```

Example call to Open Babel within make_conformator_and_params.py:

```
#confs.sdf is a resulting file with all conformers from starting sub-chunk
#individual_confs.sdf is the base name of all individual conformer files after
being split by babel
#-m – denotes babel to split a multiple molecule file into multiple individual
molecule files
babel confs.sdf individual_conf.sdf -m
```

Example call to molfile_to_params.py within make_conformator_and_params.py:

```
#Z3822861734_0.sdf is the first conformer of an example ligand from the REAL
library
#-n – name to use to keep with the corresponding params file
#--keep-names – retain the name of the ligand instead of default names chosen by
script
#--long-names – allow full ligand name to be kept instead of 1 or 3 letter short-
hand
#--clobber – overwrite any existing file of same name
#--no-pdb – do not make a pdb representation of ligand when making .params file
python rosetta/source/scripts/python/public/molfile_to_params.py Z3822861734_0.sdf
-n Z3822861734_0 --keep-names --long-names --clobber --no-pdb
```

After running make_conformator_and_params.py on a file of ligands, a tar compressed directory is made that contains the conformers and Rosetta params. The folder is made in the directory that the script is called, and is named after the source sdf files. For example, a file named “ligand_files.sdf” would result in a folder named “ligand_files”.

The example ligand_files directory contains the following:

```
confs_named.sdf – This file contains all ligand conformers that were generated by
by Conformator, grouped in a single file. The conformers are named by their source
ligand name followed by and underscore and a unique number to designate the
conformer.
ligand_files_name_list.txt – This file contains a list of all conformers with
unique numerical identifiers, separated with one conformer per line. This file can
be useful to easily accessing all file names. The file name before “_name_list.txt”
is derived from the name of the input file.
ligand_files.sdf – This is a copy of the original ligands file for reference.
single_conf_params – This is a folder filled with Rosetta params files with one
file per conformer.
single_conf_sdfs – This is a folder with each conformer in sdf format for shape
alignment and comparison with ShapeDB.
```

#### *Conformer Library Shape Database Generation*

The program VAMS ShapeDB was used to generate a database of molecular shapes to compare for all conformers within the conformer library against a target molecular shape. All ligands were aligned using the script align.py:

```
#argument 1 – name conformers file in sdf format
#argument 2 – name of output file of aligned conformers
python ginsparg_thymelab_thesis/shapedb/align.py confs_named.sdf aligned.sdf
```

The aligned conformers were run through ShapeDB's Create function to make the database. The shell script make_db_and_condense_params.sh functions with the ShapeDB Singularity image to automate the process of database generation along with the step in the next section "Conformer Params Files Compression":

```
#argument 1 – name of compressed directory
ginsparg_thymelab_thesis/shell_scripts/make_db_and_condense_params.sh
ligand_files.tar.gz
```

Example call to ShapeDB Create:

```
#-h flag allows hydrogens to be kept in creating database
#-Create flag indicates making a molecular shape index
#-in indicates the name of the input file (aligned conformers in aligned.sdf)
#-db indicates the name of the output database file that can be used for shape
comparison
/pharmit/src/build/shapedb -h -Create -in aligned.sdf -db db.db
```

#### *Conformer Params File Compression*

To reduce the memory overhead of the conformer library, a text file was generated to reduce the representation of the set of conformer params files for a ligand. Redundant data within each file was condensed into a shorthand to eliminate the presence of long repeated strings across multiple files. The script condense_conformer_params_data.py looks at all params files in the directory where it is called and makes the corresponding text files for each ligand. Generated text files are named (ligand_name)_shorthand_params.txt, such as Z3822861734_shorthand_params.txt.

Example call to condense_conformer_params_data.py:

```
#call this script within the directory of params files to condense
python
ginsparg_thymelab_thesis/params_file_compression/condense_conformer_params_data.py
```

From a condensed file, a single params file can be extracted back to its original state using the script extract_single_param_from_condensed_file.py:

```
#argument 1 – name of shorthand params file
#argument 2 – conformer number identified (0-15)
#argument 3 – name of ligand, used in writing output file
python
ginsparg_thymelab_thesis/params_file_compression/extract_single_param_from_condense
d_file.py Z3822861734_shorthand_params.txt 0 Z3822861734
```

make_db_and_condense_params.sh creates a new compressed directory with the same name as the previous file, effectively overwriting it, and containing the following:

```
db.db – This folder contains the ShapeDB database for all aligned conformers. This
will be used for the ShapeDB NNSearch operation step.
single_conf_params – This is a folder filled with text files that contain the
condensed data for all ligand conformers. Each file represents all conformers, and
params data can be extracted with extract_single_param_from_condensed_file.py
ligand_files_name_list.txt – This file contains a list of all conformers with
unique numerical identifiers, separated with one conformer per line. This file can
be useful to easily accessing all file names. The file name before "_name_list.txt"
is derived from the name of the input file.
ligand_files.sdf – This is a copy of the original ligands file for reference.
```

After extracting a params file with extract_single_param_from_condensed_file.py, a cleanup script, fix_condensed_param_file_spacing.py, is necessary to correct the formatting for Rosetta:

```
#argument 1 – input decompressed params file to be fixed
#this creates a corrected decompressed params file with the prefix "fixed_", which
can later be renamed
#In this example: fixed_Z3822861734_0.params
python ginsparg_thymelab_thesis/params_file_compression/
fix_condensed_param_file_spacing.py Z3822861734_0.params
```

#### *Conformer Shape Selection*

A query molecule for searching the conformer library for similar shapes was aligned against the other aligned molecules. The aligned ShapeDB database was compared against the query molecule using the nearest neighbor NNSearch function. This was repeated for each input shape, creating a similarity score file for all conformers compared to each input shape.

The shell script nnsearch.sh can be used with the ShapeDB container to automate the steps involved to run NNSearch and collect lists of similarity scores. The script is run on the compressed directory that was created using the make_db_and_condense_params.sh.

Example call to nnsearch.sh:

```
#argument 1 - the compressed folder that includes a shapeDB database from
make_db_and_condense_params.sh
#argument 2 – the target ligand to compare against the conformer database, aligned
to the database
./nnsearch.sh ligand_files.tar.gz suvo_shifted.sdf
```

Example call to ShapeDB NNSearch for getting a similarity score list against suvorexant:

```
# -NNSearch flag indicates to use shapedb's nnsearch
# -k flag indicates the number of nearest neighbors to return by shape similarity;
ideally have k = total number of conformers in aligned database so that we return
information on all conformers
# -ligand indicates target ligand file that is used to compare against all ligands
in the database; in this case, suvorexant aligned to the conformer library
# -db command indicates which shapedb database file to use against target ligand
# -print indicates to write out the nearest neighbors to output stream
# The written NNSearch results were piped into a text file for further processing
```

```
/pharmit/src/build/shapedb -NNSearch -k 10000 -ligand suvo_shifted.sdf -db db.db -
print >  scored_confs_against_suvo_shifted.txt
```

The resulting text file contains a list of conformers, sorted by shape similarity scores that range from 0 to 1. The lower the score, the more similar the conformer is to the input shape.

```
Z1366118392_7   0.616334795952
Z1366118392_1   0.62587761879
Z1366118392_2   0.634557366371
Z3847735140_2   0.63533437252
Z3670357625_6   0.638292491436
Z1366118392_4   0.638695240021
Z1366118392_10  0.653636813164
Z3847735140_8   0.654058337212
Z3670357625_11  0.655198395252
Z3847735140_7   0.655532360077
```

If the target structure does not contain a molecule to use as input for ShapeDB NNSearch, the binding pocket cavity can be used instead. One way the cavity shape can be generated is with the `output_space_fill_matrix_pdbs` flag in the REAL-M `ligand_discovery_search_protocol` app. This app will produce the output binding pocket in .pdb format, and Open Babel can be used to convert the pocket representation into .sdf format. The pocket needs to be aligned with the library, and then it can be used as an input for shape similarity in the same way as a molecular target.

Score lists for the entire conformer library were collected and sorted conformers with the lowest scores for the REAL-M discovery algorithm. Score lists for batches of 100 consecutive chunks were compiled into larger lists to reduce the number of lists being compared. The script get_top_ligands_top_chunk_top_lists_between_2_super_chunk_lists.py was used for multiple iterations to select the best-scoring conformers. Two files were merged and sorted by similarity score, and the top scoring conformers were retained. Merged lists were iteratively compared against each other in binary comparisons until all lists were effectively merged. We set 8,000,000 as the target maximum number of conformers.

Example call to get_top_ligands_top_chunk_top_lists_between_2_super_chunk_lists.py:

```
#argument 1 – prefix used for identifying target shape; in this example, suvorexant
aligned to conformer library
#argument 2 – maximum number of conformers to retain in merged list
#argument 3 – string to identify the range of chunks represented in first input
file
#argument 4 – string to identify the range of chunks represented in second input
file
#argument 5 – string to identify the range of chunks represented in merged output
file

python
ginsparg_thymelab_thesis/shapedb/ligand_score/get_top_ligands_top_chunk_top_lists_b
etween_2_super_chunk_lists.py suvo_shifted 8000000 00000_00099 00100_00199
00000_00199
```

Naming convention for shape similarity scores:

```
#(1) - identifying target prefix
#(2) - maximum number of conformers
#(3) - chunk range minimum
#(4) - chunk range maximum

(1)_best_(2)_chunks_(3)_(4).txt

#example file name for resulting suvorexant score from chunks 00000-00099
suvo_shifted_best_8000000_chunks_00000_00099.txt
```

The script binary_merge_of_top_shapedb_ligand_files_master.py was used to run instances of get_top_ligands_top_chunk_top_lists_between_2_super_chunk_lists.py to compare all list files for a given target shape against each other in 10 rounds of binary pairing, resulting in a single final list.

Example call to binary_merge_of_top_shapedb_ligand_files_master.py:

```
#argument 1 – prefix used for identifying target shape; in this example, suvorexant
aligned to conformer library
#argument 2 – maximum number of conformers to retain in merged list
#argument 3 – Path to the location of
get_top_ligands_top_chunk_top_lists_between_2_super_chunk_lists.py
#This example would result in making a file called:
suvo_shifted_best_8000000_chunks_00000_53099.txt
#Note, this script was originally written to utilize the Slurm Workload Manager to
allow binary comparisons to occur in parallel. The calls to Slurm for job
submission and queue monitoring can be adapted to other workload managers.

python
ginsparg_thymelab_thesis/shapedb/ligand_score/binary_merge_of_top_shapedb_ligand_fi
les_master.py suvo_shifted 8000000 ginsparg_thymelab_thesis/shapedb/ligand_score/
```

Example format of a top conformers file:

```
#similarity score (negative),ligand name and conformer number, chunk, subchunk
-0.575547099113,Z1560248380_15,35097,1
-0.597168087959,Z3726527945_14,50897,4
-0.598591566086,Z3725859640_6,50505,0
-0.602205276489,PV-004903415132_3,43637,5
-0.603496074677,PV-004531279940_9,42243,7
-0.604922473431,PV-006308993672_13,48903,3
```

REAL-M Discovery Search

***File Preparation***

Using a list of similar ligands and their accession locations, selected ligands are extracted from the conformer library into directories of user-designated size and compressed.  The script pull_conformers_from_conformer_library.py script calls pull_conformer_sub.py, extract_single_param_from_condensed_file.py, and fix_condensed_param_file_spacing.py:

```
#argument 1 – source file in csv format that has the ligand with conformer
(separated by underscore), chunk, and subchunk
#Example: PV-006308993672_13,48903,3
#argument 2 – Bucket output location
#argument 3 – Number of ligands to cluster together to be used in Rosetta discovery
#argument 4 – Location and script for pull_conformer_sub.py
#argument 5 – Location and script for extract_single_param_from_condensed_file.py
#argument 6 – Location and script for fix_condensed_param_file_spacing.py
#Note, this script was originally written to utilize the Slurm Workload Manager to
allow binary comparisons to occur in parallel. The calls to Slurm for job
submission and queue monitoring can be adapted to other workload managers.
#The script uses s3cmd to work with buckets.

python
ginsparg_thymelab_thesis/prepare_ligands_for_discovery/pull_conformers_from_conform
er_library.py top_conformers_for_discovery.csv s3://Ox_truncations_12M/ 1000
ginsparg_thymelab_thesis/params_file_compression/extract_single_param_from_condense
d_file.py
ginsparg_thymelab_thesis/params_file_compression/fix_condensed_param_file_spacing.p
y
```

Bundles of 1,000 conformers were stored in "test_params" directories – the name that Rosetta requires for large batches of ligand files – inside uniquely named outer directories. The test_params directories also contain a list file called residue_types.txt, which is a list of all conformer params files. Additional files called patches.txt and exclude_pdb_component_list.txt must be present in the test_params folder but are left empty for this usage of Rosetta.

Example contents of residue_types.txt:

```
## the atom_type_set and mm-atom_type_set to be used for the subsequent parameter
TYPE_SET_MODE full_atom
ATOM_TYPE_SET fa_standard
ELEMENT_SET default
MM_ATOM_TYPE_SET fa_standard
ORBITAL_TYPE_SET fa_standard

## Params files
Z1560248380_15.params
Z3726527945_14.params
Z3725859640_6.params
PV-004903415132_3.params
PV-004531279940_9.params
PV-006308993672_13.params
```

#### *Ligand Discovery Search*

This app calls a LigandDiscoverySearch object to place the conformers in the test_params directory against the selected input protein PDB file, using motifs. Below is an example of the Rosetta integration test for this protocol, which docks suvorexant in the pocket of the 4S0V peptide chain using motifs in motif_list.motifs:

```
#call to ligand_discovery_search_protocol.linuxgccrelease that takes command
arguments from “flags”
```

```
/rosetta/source/bin/ligand_discovery_search_protocol.linuxgccrelease @flags
```

Contents of flags:

```
#keep seed constant
-constant_seed

#input empty receptor protein
-s 4s0v_receptor_only.pdb

#directory of ligand(s) to attempt to dock
-params_directory_path test_params/

#ligand motifs library
-motif_filename motif_list.motifs

#index of residue(s) to dock ligands against
-protein_discovery_locus 423

#minimum cutoffs for fa_atr, fa_rep, and ddg to be under
-fa_atr_cutoff = -2
-fa_rep_cutoff = 150
-ddg_cutoff = -9

#constrain coordinates
-constrain_relax_to_start_coords

#optional flags for demonstrative purposes of extra features of app:

#space fill method
#define cube-shaped binding pocket about coordinate (coordinate corresponds to
within 4s0v.pdb, shifts with script)
-binding_pocket_center_sf 54,6,53
-binding_pocket_radius_sf 7
#define cutoff of how much of binding pocket volume must be filled compared to
empty pocket (>15% more filled than empty when ligand is placed)
-space_fill_cutoff_differential_score_sub 0.15

#optional export of space fill matrices to PDB (only recommended for debugging and
tuning cutoffs for binding pocket)
#Including in this test, since it can help show off the feature and troubleshoot if
this test breaks
-output_space_fill_matrix_pdbs true

#placement motifs collection
-collect_motifs_from_placed_ligand true

#motifs against residues of interest
-significant_residues_for_motifs 447,450,419,134,86,423,451,85,62
-minimum_motifs_formed_cutoff 6
-minimum_significant_motifs_formed_cutoff 1

#mandatory at least 1 real motif
-minimum_ratio_of_real_motifs_from_ligand 0.01
```

```
#must form a motif with ASN423
-mandatory_residues_for_motifs 423

#check real motifs
-check_if_ligand_motifs_match_real true
-duplicate_dist_cutoff 1.2
-duplicate_angle_cutoff 0.45

#do not output generated ligand motifs as pdbs
-output_motifs_as_pdb false

#keep x number of placements, if set to 0, keeps all
-best_pdbs_to_keep 0
```

Post-Discovery Ligand Selection

The following workflow was followed to select ligands for synthesis. The main steps (grey) of the pipeline are listed in descending order. Actions included in each main step are listed to the right in the order that they occur, with blue designating processing steps and orange filtering.

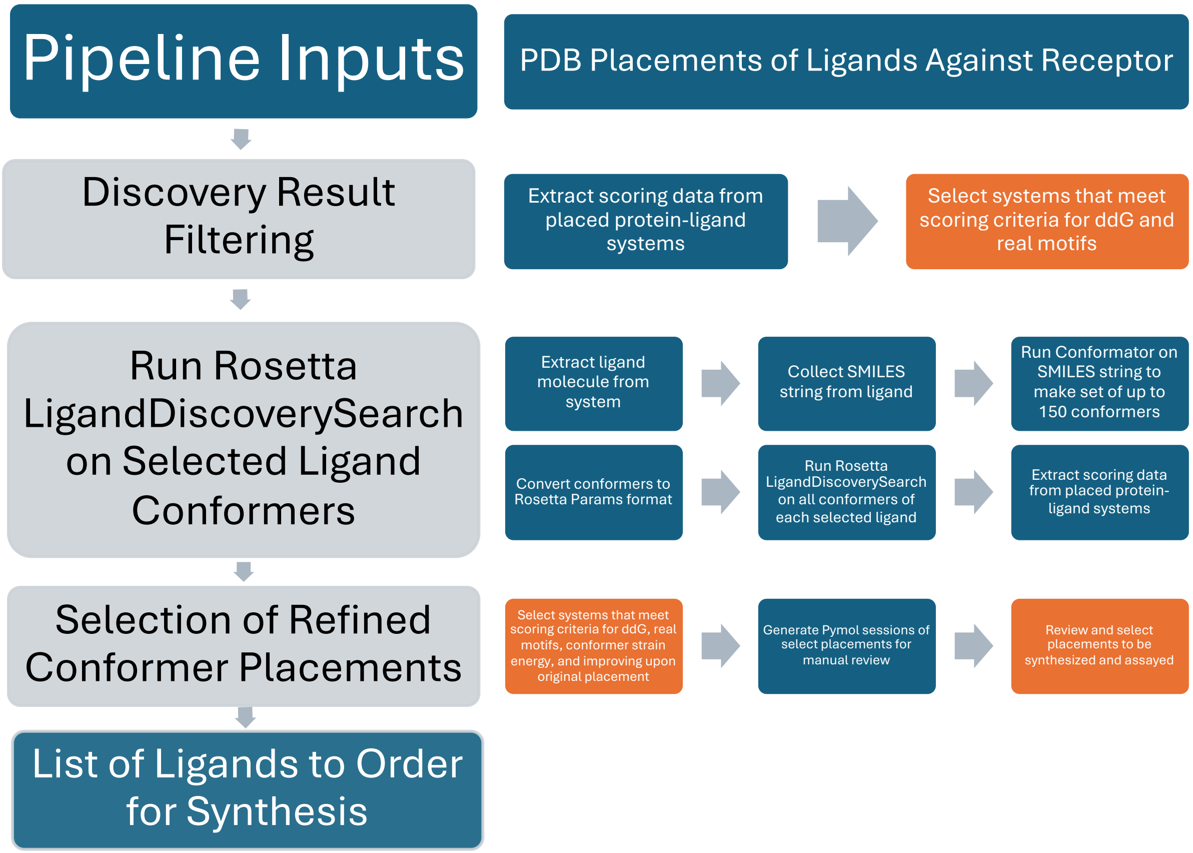


***Preliminary Ligand Scoring Evaluation***

After the discovery algorithm was complete, a post-discovery selection script was used to compare each placement for prioritization. The script score_placed_ligands_with_filtering.py was run on each output directory, quantifying and listing characteristics such as the following:

```
-ddg: The free energy score of the system with a placed ligand, determined by
Rosetta.
-total_motifs: Total number of motif-like interactions identified by Rosetta
between the ligand and protein.
-significant_motifs: Total number of motif-like interactions identified by Rosetta
between the ligand and protein. This only counts interactions against pre-
determined residues of interest using the significant_residues_for_motifs in
Rosetta discovery.
-real_motif_ratio: The ratio of motif-like interactions identified in total_motifs
that resemble motifs from the starting motif library.
-hbond_motif_count: The total number of motif-like interactions that are recognized
by Rosetta to be a hydrogen bond and have an energy value.
-hbond_motif_energy_sum: The sum of hydrogen bond energy values from motif-like
interactions that is calculated by Rosetta.
-closest_autodock_recovery_rmsd: The root mean square distance of the closest
placement of the ligand when trying to dock the ligand against the receptor using
AutoDock Vina.
-closest_autodock_recovery_ddg: The free energy score from AutoDock Vina that
corresponds to the placement that produced the value for
closest_autodock_recovery_rmsd.
-strain_energy: The strain energy of the conformer that is measured by the Tldr
Strain program.
```

Example call to score_placed_ligands_with_filtering.py:

```
#This program requires being operated in a conda environment with the following
packages: openbabel and rdkit
#All flags are optional, and most scoring/functionality will be skipped if
corresponding flags are not used.
#--working_location – flag to indicate path of directory of placement files; will
use current location if flag is not used
#--autodock_vina_path – flag to the AutoDock Vina executable, to try to recover the
Rosetta placement; AutoDock will not be attempted without this flag
#--autodock_script_path – flag to the path of the helper script,
run_autodock_on_placed_ligands.py; AutoDock will not be attempted without this flag
#--maximum_autodock_recovery_rmsd – Flag to indicate the maximum allowable root
mean square distance in angstroms between the closest AutoDock placement and the
Rosetta placement. Higher values will cause the ligand to be skipped.
#--torsion_strain_path – flag to the location of the Tldr Strain script,
Torsion_Strain.py; Strain will not be attempted without this flag
#--torsion_script_path – flag to the path of the helper script,
run_torsion_check_on_placed_ligands.py; Strain will not be attempted without this
flag
#--maximum_strain_energy - Flag to indicate the maximum allowable strain energy for
the conformer that is calculated by the Strain program. Higher values will cause
the ligand to be skipped.
#--kill – flag to choose whether to ignore ligands that AutoDock completely fails
to place or Strain does not measure energy
#--minimum_real_motif_ratio – minimum allowable value for real_motifs_ratio for
ligands to be kept
#--get_motif_hbond_energies – choose to collect and report hydrogen bond energies
from motif-like interactions
#--residue_correction_key_file – flag for the user-named file that functions as a
key to translate residue indices to correct for index discrepancies that can occur
```

```
when Rosetta interprets an input PDB. This file is used when looking at motif
interactions from a placement
#--mandatory_motif_residues – flag to indicate residues that
#--weights_path – flag to a user-named file that contains weights to apply to
placement values to derive a final composite score

python
ginsparg_thymelab_thesis/discovery_placement_filtering/placement_scoring/score_plac
ed_ligands_with_filtering.py \
  --autodock_script_path ginsparg_thymelab_thesis/autodock_recovery/ \
  --autodock_vina_path /autodock_vina_1_1_2_linux_x86/bin/vina \
  --weights_path weights_file.txt \
  --torsion_script_path ginsparg_thymelab_thesis/tldr_torsion/ \
  --torsion_strain_path /STRAIN/STRAIN_FILTER \
  --working_location working_location/ \
  --minimum_real_motif_ratio 0.5 \
  --kill True \
  --residue_correction_key_file 4s0v_7l1u_key.csv \
  --mandatory_motif_residues 86 \
  --maximum_autodock_recovery_rmsd 3.5
```

Running score_placed_ligands_with_filtering.py creates two score .csv files named raw_scores.csv and weighted_scores.csv. The raw_scores.csv file uses values directly from the analyzed placement PDB files. The weighted_scores.csv file uses the values found in the raw scores and multiplies them by any score weights that were inputted in the --weights_path flag. The score weights functionality was not used in this study. Multiple files can be appended together for later analyses, and file headers should be removed before appending. An example file for the –weights_path flag can be found at ginsparg_thymelab_thesis/discovery_placement_filtering/placement_scoring/example_weights_file/score_weights.csv.

An example file for the --residue_correction_key_file flag can be found at: ginsparg_thymelab_thesis/discovery_placement_filtering/placement_scoring/example_residue_conversion_key/4s0v_7l1u_key.csv.

By default, the scoring csv files list the placement PDBs and not the full paths. The script ginsparg_thymelab_thesis/discovery_placement_filtering/placement_scoring/append_path_to_placement_file_name_in_score_files.py can be run on a score file, and it appends the path that the script was called in to the front of file names.

Example call to append_path_to_placement_file_name_in_score_files.py:

```
#This script uses the location that it is called from and applies that location to
the front of all files that are listed in the raw_scores.csv and
weighted_scores.csv files that are in that location.
#The script does not use any command line arguments

python append_path_to_placement_file_name_in_score_files.py
```

The script ginsparg_thymelab_thesis/discovery_placement_filtering/placement_scoring/copy_best_scoring_ligands_to_target_directory.py can be used to copy placed ligands from score files to a new target location. When used, placement files are iteratively copied to folders that are created in the target directory. The folders are named by numbers starting at zero, and up to 100 placements are copied into each folder. For example, if there were 120 placements, the first 100 palcements would be copied into a folder titled “0” in the target directory, and the last 20 would be copied into a folder named “1”. This can be useful for later analysis. The dividing of files into folders in batches of 100 allows for ease of use when running parallelized programs to process the data.

Example call to a copy_best_scoring_ligands_to_target_directory.py:

```
#argument 1 – Score csv file (can be of any name) that contains placement files of
interest. There must be one file per line, and the file must be in the first
position.
#argument 2 – Target destination for all placement files in the score file to be
copied to.

python
ginsparg_thymelab_thesis/discovery_placement_filtering/placement_scoring/copy_best_
scoring_ligands_to_target_directory.py 12M_-17_real_hbond_ddg_sorted.csv 12M_-
17_real_hbond_ddg_sorted/
```

After scoring the initial placements, placed ligands were selected if they met designated criteria. For example, in the main run for antagonists of HCRTR2 used the following filters:

- The rate of motifs identified around the ligand were found to match motifs from the Protein Data Bank at a rate of at least 50%.
- The rate of motifs identified around the ligand were found to match motifs from the Protein Data Bank at a rate of at least 25% AND the system free energy was less than or equal to -16 Rosetta Energy Units (REU).

The selection process is detailed in Jupyter notebooks in ginsparg_thymelab_thesis/jupyter_notebooks/initial_placement_selection.

#### *Placement Refinement*

An expanded set of conformers were generated for all ligands passing the initial selection step. This expansion serves to further prioritize ligands by optimizing fits and minimizing minor clashes. First, the placed ligand conformers were extracted from their placement PDB and converted to .sdf format. Placements in each directory were extracted using the script ginsparg_thymelab_thesis/discovery_placement_filtering/placement_pymol_session_creation/convert_placement_pdb_ligands_to_sdf.py:

```
#This script looks at all .pdb files in the directory that it is called from, and
attempts to extract all molecules with HETATM tags (i.e. placed ligands) to convert
into .sdf format.
#The script creates an intermediate .pdb file that only contains HETATM data, which
is used for the conversion to .sdf
#This script uses OpenBabel software to convert from .pdb to .sdf, and requires an
OpenBabel executable called as “obabel” to work. The OpenBabel Conda package is
supported for this.
#This script does not use any command line arguments.
```

```
python ginsparg_thymelab_thesis/
discovery_placement_filtering/placement_pymol_session_creation/convert_placement_pd
b_ligands_to_sdf.py
```

Example call to Open Babel within convert_placement_pdb_ligands_to_sdf.py:

```
#-O – Indicates output file name

obabel ligand.pdb -O ligand.sdf
```

After the ligands are converted to .sdf format, the script ginsparg_thymelab_thesis/conformator/run_conformator_on_all_files_in_directory_using_smiles.py used Conformator to create expanded conformer sets of the placed ligands. Extracting a Simplified Molecular Input Line Entry System (SMILES) string from the placement ligand and using it as an input to Conformator creates the same set that would be created by the original ligand.

Example call to run_conformator_on_all_files_in_directory_using_smiles.py:

```
#This script looks at all .sdf files in the directory that it is called from
argument 1 to create conformer sets using Conformator.
#The script creates an intermediate .smi file that contains the SMILES string of
the initial ligand to be used as an input for Conformator.
#This script needs to be run in an environment that has the rdkit package.

#Argument 1 – Path to the directory that contains the placed ligand .sdf files.
#Argument 2 – The name Conformator executable to be called (with path to the
executable).

python
ginsparg_thymelab_thesis/conformator/run_conformator_on_all_files_in_directory_usin
g_smiles.py workinglocation/ conformator_1.2.1/conformator
```

Example call to Conformator within run_conformator_on_all_files_in_directory_using_smiles.py:

```
#This call to Conformator does not use the -n flag, and uses its default conformer
limit of 250.

conformator -i ligand.smi -o ligand_confs.sdf --keep3d --hydrogens -v 0
```

With the new expanded conformer files generated, the script ginsparg_thymelab_thesis/ conformator/split_sdf_files_in_directory_to_individual.py was used to split the output file with all conformers for a ligand to one conformer per file, putting each file into a unique directory.

Example call to split_sdf_files_in_directory_to_individual.py:

```
#This script looks at all conformer .sdf files in the directory that it is called
from argument 1 to split conformer files into individual .sdf files
#This script uses OpenBabel software to convert from .pdb to .sdf, and requires an
OpenBabel executable called as “obabel” to work. The OpenBabel Conda package is
supported for this.
```

```
#Argument 1 – Path to the directory that contains the expanded ligand conformer
sets in .sdf format.

python
ginsparg_thymelab_thesis/conformator/split_sdf_files_in_directory_to_individual.py
working_directory
```

Example call to Open Babel within split_sdf_files_in_directory_to_individual.py:

```
#-O – Indicates output file name (with path to folder named after ligand)
#-m – Indicates to put each conformer in ligand_confs.sdf into separate files with
unique numbers at the end (i.e. ligand_0.sdf, ligand_1.sdf, etc.)

obabel ligand_confs.sdf -O ligand/ligand_.sdf -m
```

Then, params files for each conformer were created using the script ginsparg_thymelab_thesis/params_file_compression/make_params_files_from_conformers.py:

```
#This script is intended to work with the folder of ligands with respective folders
of separated individual conformer .sdf files from a ligand that result from
split_sdf_files_in_directory_to_individual.py

#Argument 1 - Path to the directory that contains the expanded ligand conformer
sets in .sdf format with one conformer per file.
#Argument 2 – Path to the Rosetta python script molfile_to_params.py to be used for
converting conformers from sdf to params format.

python
ginsparg_thymelab_thesis/params_file_compression/make_params_files_from_conformers.
py ligands_location/ rosetta/source/scripts/python/public/molfile_to_params.py
```

Submissions of REAL-M discovery jobs for each new conformer were run in parallel for test_params directories, using the Slurm job distributor and script running_rosetta/ligand_discovery_search/make_and_submit_discovery_jobs_slurm.py. This script can be adapted for other job distributors or jobs can be run manually. The arguments file needs to have the argument for params_directory_path removed, as it is written by the script.

Example call to make_and_submit_discovery_jobs_slurm.py:

```
#This script is intended to work with the folder of ligands with respective folders
of separated individual test_params folders that are prepared for Rosetta
LigandDiscoverySearch.

#Argument 1 - Path to the directory that contains the ligand folders that contain
test_params directories
#Argument 2 – Path and file for the common arguments file to be used in all jobs.
This file should not contain the params_directory_path flag, as that flag will be
written by this python script
#Argument 3 – Path to the Rosetta binary executable to ligand_discovery_search

python
ginsparg_thymelab_thesis/running_rosetta/ligand_discovery_search/make_and_submit_di
scovery_jobs_slurm.py working_directory/ args_file
/rosetta/source/bin/identify_ligand_motifs.linuxgccrelease
```

The resulting placements were re-scored and the difference in scores from the original placement was also computed with the script ginsparg_thymelab_thesis/discovery_placement_filtering/placement_scoring/conformer_refinement_analysis_single_directory.py. For each folder, a new file called delta_scores.csv writes the comparisons for all placements from the original and expanded conformer sets. The Slurm job distributor was used to run an array job that ran this script for each respective ligand directory to improve throughput by running jobs in parallel.

Example call to conformer_refinement_analysis_single_directory.py:

```
#This script is intended to look at two placement data csv files that use the same
ligand for placement for both the original placement and placements from an
expanded set of conformers of the original ligand. The script compares the metrics
of the expanded conformers against the original and writes the data to a csv file.

#Argument 1 - Path and name of the initial ligand placement data csv file
#Argument 2 – Path and name of the expanded conformer placement data csv file
python ginsparg_thymelab_thesis/
discovery_placement_filtering/placement_scoring/conformer_refinement_analysis_singl
e_directory.py directory_folder/initial_ligand_placement_score_data.csv
directory_folder/expanded_conformer_placement_score_data.csv
```

All delta_scores.csv files were appended together for analysis in Jupyter notebooks. Conformers were selected for manual selection based on selected criteria, depending on the run. Placements selected for manual review also had the torsion strain of the ligand evaluated using the Torsion_Strain.py script from the TLDR Strain package. The script ginsparg_thymelab_thesis/discovery_placement_filtering/tldr_torsion/run_torsion_check_on_placed_ligands.py was used to automate running the strain script on all selected placements for manual review.

Example call to ginsparg_thymelab_thesis/discovery_placement_filtering/tldr_torsion/run_torsion_check_on_placed_ligands.py:

```
#This script is intended to look at a folder of placement pdb files and use the
tldr Strain script to determine the torsions strain energy of each ligand. The
script extracts the ligand from each placement into a new pdb file, and converts
the pdb file to mol2 format so that tldr can analyze it.

#Argument 1 - Path and name of the directory with the placement pdb files to
analyze for torsion strain energy
#Argument 2 – Path to the location of the location of the tldr Strain package where
the Torsion_Strain.py script is.

python
ginsparg_thymelab_thesis/discovery_placement_filtering/tldr_torsion/run_torsion_che
ck_on_placed_ligands.py placements_for_review/ path/to/STRAIN/STRAIN_FILTER/
```

Example call to Torsion_Strain.py:

```
#This script is intended to look at a folder of placement pdb files and use the
tldr Strain script to determine the torsions strain energy of each ligand. The
script extracts the ligand from each placement into a new pdb file, and converts
the pdb file to mol2 format so that tldr can analyze it.
```

```
#Argument 1 - Path and name of mol2 file of placed ligand

python Torsion_Strain.py path/to/placed_ligand.mol2
```

Ligand conformations with a torsion score based on the average of the upper and lower bounds of the predicted range that were greater than 8 energy units were eliminated from consideration. Torsion score has a minimum of 0 and unbound maximum.

Manual review was completed using Pymol, using the script ginsparg_thymelab_thesis/discovery_placement_filtering/placement_pymol_session_creation/make_pymol_sessions_of_placements_pymol2.py to make sessions. Residues of interest were displayed as spheres under certain conditions and colored as follows:

- Cyan – Default backbone/residue color
- Yellow – Residues within 5 Å of ligand
- Orange – User-specified residues of interest
- Brown – Not real motif-like interaction with residue, spheres shown
- Magenta – Real motif-like interaction with residue, spheres shown
- White – Ligand

Example call to ginsparg_thymelab_thesis/discovery_placement_filtering/placement_pymol_session_creation/make_pymol_sessions_of_placements_pymol2.py:

```
#This script is intended to create a readable Pymol session on a directory of placement pdb files. Arguments are used to help indicate residues of interest to highlight in the session.

#Argument 1 - Path and name of directory of placement pdb files to add to a Pymol session. All pdb files only in the directory will be added to the session.
#Argument 2 – Position index of ligand in pdb files (this program assumes all pdb files are of the same protein backbone with different ligand placements)
#Argument 3 – Pymol string of residue index/indices to highlight. If selecting multiple residues, each index must be separated by a ‘+’: i.e. 86+134+253
#Argument 4 – (Optional) Path and name of a key file in csv format to allow for post-hoc conversions of residue indices to a custom set. This is to help resolve discrepancies from Rosetta indexing conventions. A key file has no header line, and has 3 entries per line: line identified string (not used), original index value, value to convert index to

python
ginsparg_thymelab_thesis/discovery_placement_filtering/placement_pymol_session_creation/make_pymol_sessions_of_placements_pymol2.py placements_directory/ 282
86+134+253 7l1u_residue_key.csv
```

Benchmarking REAL-M Discovery Search

***Benchmark Library Acquisition***

REAL-M was benchmarked on an 84 protein-ligand system set from the Database of Useful Decoys-Enhanced (DUD-E). From the 102 protein-ligand systems in the DUD-E library, systems were eliminated if there were compatibility issues with running them in any of the software's being compared. Code and files for the benchmark can be found in the GitHub repository: https://github.com/toadlover/patel_ginsparg_rosetta_lds_dude_benchmark.

All software started with an initial single-chain empty protein receptor in .pdb format. The software was also given the corresponding ligand in .sdf and .mol2 format, which starts in its native conformation. Rosetta, Dock6, and AutoDock Vina all use the native conformation as a starting point for docking. Rosetta and Dock6 directly use the inputted rigid native ligand conformation to sample docking placements. AutoDock uses the native conformer as a starting point for sampling conformers to attempt to dock. Schrodinger does not take the native ligand conformation into account in conformer generation or docking.

To prepare DUD-E systems for Rosetta, the protein receptor for each system was separated from the ligand. The ligand was converted to params format, with a corresponding test_params directory created for Rosetta discovery. Files were created to hold all arguments for running Rosetta on each system. Input files, except for motifs files, can be found for respective systems in patel_ginsparg_rosetta_lds_dude_benchmark/files/all

***Non-Homologous Motif Library Preparation***

To ensure that the original protein PDB file or homologous PDB systems were not being used to place the ligand for a given system, unique motifs lists were created for each protein-ligand system, removing any motifs that were derived from homologous systems. To determine homologous PDB systems, BLAST version 2.10.0 was run from the command line on all protein receptors in FASTA format. The FASTA strings of all receptors are stored in patel_ginsparg_rosetta_lds_dude_benchmark/files/fasta/fasta_compiled.fasta.

blastp command used from patel_ginsparg_rosetta_lds_dude_benchmark/files/fasta/fasta.job:

```
#results were outputted to
patel_ginsparg_rosetta_lds_dude_benchmark/files/fasta/fasta_out.log
ncbi-blast-2.10.0+/bin/blastp -query fasta_compiled.fasta -evalue 0.05 -db pdbaa -
remote > fasta_out.log
```

The script patel_ginsparg_rosetta_lds_dude_benchmark/files/fasta/ remove_homologous_pdbs_from_motif_list.py parsed the BLAST output and created unique motifs lists for each system:

```
#Argument 1 – BLAST output file, which notes homologous PDB entries by 4 letter
code that correspond to DUD-E systems
#Argument 2 – Initial motifs list that will be copied into unique lists per system,
with motifs from homologous systems removed
python
patel_ginsparg_rosetta_lds_dude_benchmark/files/fasta/remove_homologous_pdbs_from_m
otif_list.py fasta_out.log FINAL_motifs_list_filtered_2_3_2023.motifs
```

The unique motif lists are stored at patel_ginsparg_rosetta_lds_dude_benchmark/files. Within the GitHub repository, the motifs files are compressed in .bz2 format. The shell script patel_ginsparg_rosetta_lds_dude_benchmark/files/decompress_motifs.sh needs to be run to decompress the motifs files before use. There is a corresponding compress_motifs.sh script that compresses all motifs files in the patel_ginsparg_rosetta_lds_dude_benchmark/files directory.

#### *Library Benchmarking*

To run the benchmark, patel_ginsparg_rosetta_lds_dude_benchmark/scripts/benchmark.py automates the pipeline of the REAL-M run and analysis of the results:

```
python patel_ginsparg_rosetta_lds_dude_benchmark/scripts/benchmark.py
```

This script requires an arguments file named bench_params:

```
-l /data/user/abgvg9/FINAL_motifs_list_filtered_2_3_2023.motifs #Motifs library
location
-b /scratch/abgvg9/rosetta_may_for_checkin/rosetta/ #Path to top of Rosetta
Repository
-r 100 #fa_rep_cutoff
-a -5 #fa_atr_cutoff
-s 0.8 #space fill sub area score cutoff
-m false #print the space fill matrix using the output_space_fill_matrix_pdbs flag;
used for debugging
-d 0.3 #space fill sub area differential score cutoff
```

The benchmark.py script calls upon the following scripts found in patel_ginsparg_rosetta_lds_dude_benchmark/scripts/, listed in the order that they are called:

- auto_place.py – Runs Rosetta LigandDiscoverySearch on all systems with all pre-defined anchor residues.
- auto_rmsd.py – Determines the RMSD of all Rosetta-predicted placements against the DUD-E placement, and determines the closest Rosetta-predicted placement to the DUD-E placement for a given anchor residue for a single system.
- auto_rmsd_total.py – Compiles RMSD data and determines the closest Rosetta-predicted placement to the DUD-E placement across all starting anchor residues for each system.
- auto_time.py – Determines the amount of time spent on discovery, which was primarily useful in developing the method, and optimizing operations for runtime.
- stats.py – Compiles all data into a log text file named stats.txt that simplifies the results of the benchmark.

Example stats.txt contents from run 116 results:

```
Initial Receptors
102
Number of Params Generated
91
Number of AA found
463
Number of Residues with placements found
283
Number of Receptors with placements found
85

Placements within 1A
73
Placements between 1A and 2A
6
Placements between 2A and 5A
```

```
4
Placements greater than 5A
2

Avg Time: 25.958747300215983
Max Time: 443.53333333333336
Time within 5: 164
Time between 5 and 10: 82
Time between 10 and 20: 65
Time greater than 20: 152
Timeouts: 0
```

- homology.py - Confirmation that placement motifs are not from homologous PDB files.

Benchmarking AutoDock Vina against Rosetta

The Python script args_creator.py iteratively generates a .txt file for each system containing arguments to be passed through AutoDock Vina. args_creator.py used center_finder.py to calculate the center coordinates of each ligand. autodock_runner.py generated the commands to run docking for each system, which were then manually run in the command line.

Example command with args_creator.py:

```
#generates .txt file with arguments for ADV docking

#calls on center_finder.py within script to calculate center coordinates of ligand (a necessary argument)

python args_creator.py
```

Example ADV benchmarking command:

```
#aa2ar is an example ligand from the DUD-E benchmarking library

#--config – configures docking output using given .txt argument file

“vina.exe” --config “/aa2ar/args.txt”
```

ADV_rmsd_evaluator.py iterated through the output data for each system and calculated RMSD data of placements in comparison to reference ligands. The best RMSD value for each system was written into a single .csv file for data analysis.

Example command using ADV_rmsd_evaluator.py:

```
#determines best RMSD value for each system and writes into a single .csv file

python ADV_rmsd_evaluator.py
```

Benchmarking DOCK6 against Rosetta

Scripts and input files related to the benchmark can be found in the GitHub repository: https://github.com/toadlover/thyme_lab_internship_2024/

***Preparation of PDB Files and Surface Generation***

The PDB file of each system was converted into mol2 file format using the Dock Prep tool (Tools→Structure→Editing→Dock Prep) within the Chimera software. A representation of the receptor protein's surface was then generated using the mol2 file (Actions→Surface→show), to be saved as a DMS file.

***Spheres Generation***

The Python script sphgen_runner.py was run on the target directory of protein-ligand systems to generate spherical representations of the receptors.

Example command using /scripts/dock6/sphgen_runner.py:

```
#generates spherical representation of entire receptor for each system

python sphgen_runner.py
```

***Remaining Input Generation and Docking***

The Python script /scripts/dock6/docking_runner.py executed the creation of remaining necessary input files for docking based on a user-specified sphere selection radius value, then generated the placement for each system (with the given radius). docking_runner.py iteratively used command line arguments for each step of input file generation as well as docking execution.

Example command using docking_runner.py:

```
#generates input files for docking with a selected spheres radius of 1.0, then
executes docking

python docking_runner.py 1.0
```

Below are example commands that docking_runner.py executed in the command line to generate input files and the final placement for a system.

```
#aa2ar is an example ligand from the DUD-E benchmarking library

#1.0 is an example selected spheres radius

#isolating receptor spheres within user-specified distance from ligand

/dock6/bin/sphere_selector aa2ar.sph crystal_ligand.mol2 1.0

#generating grid reference box

/dock6/bin/showbox < showbox.in

#generating energy grid and bump grid of the protein

/dock6/bin/grid -i grid.in -o gridinfo.out

#minimizing potential energy of ligand

/dock6/bin/dock6 -i min.in -o min.out

#at this point, a .in file containing parameters for docking is generated for the
specific system

#generating docking placement

/dock6/bin/dock6 -i rigid1.0.in -o /output_files/aa2ar/rigid1.0.out
```

Nine placements were generated for each system with selected spheres radii of 1.0 to 9.0. Radii values were incremented by 1.

/scripts/rmsd/dock_rmsd_eval.py iterated through the placement files in each system to calculate and choose the best RMSD value in comparison to reference ligands. Placements that failed due to insufficiently large radii were ignored in calculations. The best RMSD value for each system was written into a single .csv file for data analysis:

```
#determines best RMSD value for each system and writes into a single .csv file

python dock_rmsd_eval.py
```

Benchmarking Schrödinger-Maestro against Rosetta

Scripts and input files related to the benchmark can be found in the GitHub repository: https://github.com/toadlover/thyme_lab_internship_2024/

Protein and ligand structures were prepared using Protein Preparation Wizard and LigPrep, respectively. The former optimized hydrogen bond networks, minimized waters and generated energy-minimized protein conformations. LigPrep produced energy-minimized ligand structures with appropriate ionization states, stereoisomers, and tautomers, utilizing the OPLS4 force field. Receptor grids were defined using the centroid of the native ligand binding site within a 10 Å radius. Docking was performed with Glide in Standard Precision mode, generating up to 100 poses per ligand to return conformations closest to the native conformation. Post-docking minimization was conducted to refine docking results by improving the accuracy of the ligand's binding pose, reducing steric clashes, and ensuring a more realistic and stable protein-ligand complex. Pose Viewer was used to view the different poses, and the center coordinates of the conformations were collected for subsequent RMSD calculations. The schrodinger_rmsd.py script computed RMSD values for docked ligand poses relative to native ligand coordinates. The script reads the native ligand coordinates from mol2 files and calculates the RMSD for each docked pose based on the Euclidean distance between corresponding atoms. The distances are accumulated for each pose, the average RMSD is computed, and the pose with the lowest RMSD is identified as the best. Results for each system, including the best RMSD, pose index, and total number of poses tested were recorded for further analysis and visualization in Jupyter Notebook.

Example call to /scripts/rmsd/schrodinger_rmsd.py:

```
# Make sure to run this script from within the scripts/rmsd directory for all
location-based operations to work

# Running script: python schrodinger_rmsd.py

cd /path/to/thyme_lab_internship_2024/scripts/rmsd

python schrodinger_rmsd.py
```

Benchmarking AlphaFold 3 (AF3) against Rosetta

***Preparation and Running AF3***

17 protein-ligand systems from 2025 were selected, so that the exact protein-ligand combination would not be present in AF3 database: 9HZ0, 9I0T, 9I1H, 9IH4, 9LFU, 9LGN, 9LO7, 9LSL, 9LYF, 9M3N, 9M3O, 9M7M, 9ODR, 9OG3, 9QAC, 9QEK, 9QEL

The benchmark was started by creating a set of folders in a single location with the name of each folder being a unique name for each system in the benchmark. Within each unique folder name is at least a PDB file of the protein chain of interest with all ligands removed, with the same name as the folder and ending with ".pdb". Also in this folder is a file of the ligand in mol2 format named "crystal_ligand.mol2". For example, for the system "aa2ar", there is a folder named "aa2ar" with a file of the receptor protein named "aa2ar.pdb" and its ligand named "crystal_ligand.mol2".

To dock with AF3, there are two calls to AF3. The first is to call a multiple sequence alignment (MSA) on the residue sequence of the receptor only, which creates a new base json script that can be used for docking. The second script also needs to add the ligand in SMILES format.

The wrapper script thyme_lab_internship_2024/scripts/alphafold3/run_alphafold_on_dude_systems_new.py automates the preparation of the AF3 inputs, writes the initial MSA job script, and runs it using the AF3 container. The wrapper script utilizes two scripts from the ginsparg_thymelab_thesis repository, ginsparg_thymelab_thesis/misc/alphafold_prep/get_smiles_of_ligand_file.py (creates a SMILES .smi file of the crystal_ligand.mol2 ligand) and ginsparg_thymelab_thesis/misc/alphafold_prep/get_residue_sequences_from_pdb_file.py (creates a .csv file of the residue sequence of each chain in the empty receptor file).

Example call to get_smiles_of_ligand_file.py:

```
# This script requires a python environment that contains the rdkit package for molecule file conversion

#Argument 1 – single ligand file in sdf, mol2, or pdb format

#Produces a .smi SMILES string file of the ligand

python get_smiles_of_ligand_file.py crystal_ligand.mol2
```

Example call to get_residue_sequences_from_pdb_file.py:

```
#Argument 1 – empty protein receptor in PDB format

#Produces a .csv file where each line is a chain with the first index being the chain ID and the second index being the residue sequence of the chain

python get_residue_sequences_from_pdb_file.py aa2ar.pdb
```

Example call to run_alphafold_on_dude_systems_new.py:

```
#Argument 1 – full path to (including) the ginsparg_thymelab_thesis repository

#produces a json file and shell script to run Alphafold3 on each system in the thyme_lab_internship_2024/dude_library_simple location. The scripts are generated in the thyme_lab_internship_2024/alphafold3 location

python run_alphafold_on_dude_systems_new.py /path/to/ ginsparg_thymelab_thesis
```

Example created json script (thyme_lab_internship_2024/
alphafold3/aa2ar/af_input/aa2ar_protein_only.json):


```
{
	"name": "aa2ar",
	"dialect": "alphafold3",
	"version": 3,
	"modelSeeds": [1, 2, 3, 4, 5, 6, 7, 8, 9, 10],
	"sequences": [
		{
			"protein": {
				"id": ["A"],
				"sequence": "IMGSSVYITVELAIAVLAILGNVLVCWAVWLNSNLQNVTNYFVVSLAAADIAVGVLAIPFAITISTGFCAACHGCLFIACFVLVLTQSSIFSLLAIAIDRYIAIRIPLRYNGLVTGTRAKGIIAICWVLSFAIGLTPMLGWNNCGQSQGCGEGQVACLFEDVVPMNYMVYFNFFACVLVPLLLMLGVYLRIFLAARRQLRSTLQKEVHAAKSLAIIVGLFALCWLPLHIINCFTFFCPDCSHAPLWLMYLAIVLSHTNSVVNPFIYAYRIREFRQTFRKIIRSHVLRQ"
			}
		}
	]
}
```


Example created shell script for MSA (thyme_lab_internship_2024/
alphafold3/aa2ar/aa2ar_no_inference.sh):

```
module load apptainer
export AF3_RESOURCES_DIR=/pi/summer.thyme-umw/alphafold3
export AF3_IMAGE=${AF3_RESOURCES_DIR}/alphafold3_cuda7.sif
export AF3_CODE_DIR=${AF3_RESOURCES_DIR}/code
export AF3_INPUT_DIR=/pi/summer.thyme-umw/2024_intern_lab_space/thyme_lab_internship_2024/alphafold3/aa2ar/af_input
export AF3_OUTPUT_DIR=/pi/summer.thyme-umw/2024_intern_lab_space/thyme_lab_internship_2024/alphafold3/aa2ar/af_output
export AF3_MODEL_PARAMETERS_DIR=${AF3_RESOURCES_DIR}/params
export AF3_DATABASES_DIR=${AF3_RESOURCES_DIR}/db


apptainer exec \
     --nv \
```

```
     --env
XLA_PYTHON_CLIENT_PREALLOCATE=false,TF_FORCE_UNIFIED_MEMORY=true,XLA_CLIENT_MEM_FRA
CTION=3.2 \

     --bind $AF3_INPUT_DIR:/root/af_input \

     --bind $AF3_OUTPUT_DIR:/root/af_output \

     --bind $AF3_MODEL_PARAMETERS_DIR:/root/models \

     --bind $AF3_DATABASES_DIR:/root/public_databases \

     $AF3_IMAGE \

     python ${AF3_CODE_DIR}/alphafold3/run_alphafold.py --run_inference=FALSE --
flash_attention_implementation=xla \

     --json_path=/root/af_input/aa2ar_protein_only.json \

     --model_dir=/root/models \

     --db_dir=/root/public_databases \

     --output_dir=/root/af_output\
```

The shell script was given executable permissions using chmod and then run on a job scheduler using bsub:

```
#give executable permissions

chmod 777 aa2ar_no_inference.sh

#call shell script with bsub

bsub -n 8 -R "rusage[mem=2048]" -W 300 -gpu "num=1:gmodel=TeslaV100_SXM2_32GB-
30G:mode=shared:j_exclusive=no" -q gpu -o aa2ar_protein_preparation_log.txt bash
aa2ar_no_inference.sh
```

After running run_alphafold_on_dude_systems_new.py, the script thyme_lab_internship_2024/scripts/alphafold3/run_alphafold_on_dude_systems_new_round_2_ligand_placement.py is called to run the docking of the ligand. The script is very similar to run_alphafold_on_dude_systems_new.py, and utilizes the generated MSA json files for docking along with the same scripts from the ginsparg_thymelab_thesis repository.

Example call to run_alphafold_on_dude_systems_new_round_2_ligand_placement.py:

```
#Argument 1 – full path to (including) the ginsparg_thymelab_thesis repository

#Also requires a pre-generated MSA json file

#produces a directory of predicted placements of the input ligand

python run_alphafold_on_dude_systems_new_round_2_ligand_placement.py /path/to/
ginsparg_thymelab_thesis
```

The MSA json file is modified to insert the ligand SMILES into the sequences block of the file. The inserted block looks like this (snippet from thyme_lab_internship_2024/alphafold3/aa2ar/af_input/aa2ar_data.json):

```
          {
             "ligand": {
                   "id": "B",
                   "smiles": "NC1=NC(NCCc2ccccc2)=NC2=N[C](c3ccco3)NN12"
                   }
             },
```

Example created shell script for MSA (thyme_lab_internship_2024/ alphafold3/aa2ar/aa2ar_docking.sh):

```
module load apptainer

export AF3_RESOURCES_DIR=/pi/summer.thyme-umw/alphafold3

export AF3_IMAGE=${AF3_RESOURCES_DIR}/alphafold3_cuda7.sif

export AF3_CODE_DIR=${AF3_RESOURCES_DIR}/code

export AF3_INPUT_DIR=/pi/summer.thyme-
umw/2024_intern_lab_space/thyme_lab_internship_2024/alphafold3/aa2ar/af_input

export AF3_OUTPUT_DIR=/pi/summer.thyme-
umw/2024_intern_lab_space/thyme_lab_internship_2024/alphafold3/aa2ar/af_output

export AF3_MODEL_PARAMETERS_DIR=${AF3_RESOURCES_DIR}/params

export AF3_DATABASES_DIR=${AF3_RESOURCES_DIR}/db

apptainer exec \

     --nv \

     --env
XLA_PYTHON_CLIENT_PREALLOCATE=false,TF_FORCE_UNIFIED_MEMORY=true,XLA_CLIENT_MEM_FRA
CTION=3.2 \

     --bind $AF3_INPUT_DIR:/root/af_input \

     --bind $AF3_OUTPUT_DIR:/root/af_output \

     --bind $AF3_MODEL_PARAMETERS_DIR:/root/models \

     --bind $AF3_DATABASES_DIR:/root/public_databases \

     $AF3_IMAGE \

     python ${AF3_CODE_DIR}/alphafold3/run_alphafold.py --run_data_pipeline=FALSE -
-flash_attention_implementation=xla \

     --json_path=/root/af_input/aa2ar_data.json \

     --model_dir=/root/models \

     --db_dir=/root/public_databases \

     --output_dir=/root/af_output\
```

The shell script was given executable permissions using chmod and then run on using bsub:

```
#give executable permissions

chmod 777 aa2ar_docking.sh

#call shell script with bsub

bsub -n 8 -R "rusage[mem=2048]" -W 300 -gpu "num=1:gmodel=TeslaV100_SXM2_32GB-30G:mode=shared:j_exclusive=no" -q gpu -o aa2ar_docking_log.txt bash aa2ar_docking.sh
```

***AlphaFold 3 (AF3) Predicted Placement Recovery Analysis***

The script thyme_lab_internship_2024/scripts/alphafold3/get_placement_rmsd.py was used to determine the closest predicted AF3 placement to the native system placement. The script utilizes pymol2 to align AF3 placements to the original systems, Open Babel for file type conversions, and RDKit for the GetBestRMS() function to determine the closest placement. The use of GetBestRMS (or similar algorithm) is necessary as AF3's use of SMILES strings erodes the conservation of atom identifiers for simple pairwise distance derivation. AF3 does not provide system energies but does provide a confidence score for each placement. Confidence scores were paired with each placement distance and treated like system energies provided by other algorithms to determine the best RMSD for the most confident placement, best 10 most confident, and of all placements.

Example call to get_placement_rmsd.py:

```
#This script requires the following packages:

#pymol2

#rdkit

#numpy

#openbabel

#re

python get_placement_rmsd.py
```

Reproducing Published Discovery Outcomes with REAL-M

***Search for Enamine Drug Library Publications***

A literature review was performed to identify publications that utilized the Enamine drug library for drug discovery. The goal was to obtain the protein and ligand molecules that were experimentally success and attempt to replicate their discovery with REAL-M. The search yielded 13 publications of interest, from which protein PDB files were obtained. For ligand molecules in the mol2 format, the Open Babel software was used for conversion to sdf files.

***Automating Ligand Conformer Generation***

Ligand conformers were generated with the /scripts/rosetta_enamine_recovery/process_sdf_directory3.py script, which takes the sdf files within a given publication directory path as input, generates 3D conformers, separates them, converts them into Rosetta-compatible params files, and modifies these files for REAL-M docking. The script first defines paths for the input sdf directory, output sdf directory (‘Processed-SDFs’), and params directory (‘Params-Files’). Note that the output directories are first created for the Processed-SDFs and Params-Files directories before defining their paths.

For each sdf file in the input directory, the Conformator tool generated the ligand conformers.

Example call to Conformator:

```
conformator -i input_file.sdf -o confs.sdf --keep3d --hydrogens -n 15
```

The command reads the input SDF file, generates up to 15 conformers, and outputs them to a single confs.sdf file, while retaining the 3D-coordinates and adding hydrogen atoms.

The confs.sdf files were then split using Open Babel software with the following command:

```
obabel confs.sdf -O output_conf.sdf -m
```

The -m flag separates the multi-conformer sdf files into separate sdf files. Each new sdf file is named sequentially based on the original filename, with additional numbering to differentiate between unique conformers of the same ligand. After the conformers are separated into individual sdf files, each of the files is processed with the Rosetta suite molfile_to_params.py script for conversion of the sdf files into params files, which is the required format for Rosetta’s computational protocols. By default, the params files contain placeholders for the compound name, typically named as ‘NAME LG’ and ‘I0_String LG1’.

***Decoy Ligand Set Generation***

To emulate the REAL-M discovery pipeline, we selected sets of 10,000 ligands to place alongside the ligands from published screens. The ligands were derived from the chunks of the conformer library with the closest average molecular weight to the average molecular weight of the ligands from the respective publication. The script ginsparg_thymelab_thesis/benchmarking_other_papers/get_mw_of_sdf_ligands_in_directory.py was used to derive the average molecular weight of heavy atoms of all sdf files in a directory:

```
#This script is intended to print the average molecular weight of all ligand files
in sdf format in the current directory. Data for the molecular weight of each
detected ligand and the directory average is written to a file called
“ligand_mw.csv”.

python
ginsparg_thymelab_thesis/benchmarking_other_papers/get_mw_of_sdf_ligands_in_directo
ry.py
```

The chunks with the closest average molecular weight to the input were identified with the script ginsparg_thymelab_thesis/conformer_library_analytics/find_chunk_with_closest_average_mw_t o_request.py and file ginsparg_thymelab_thesis/conformer_library_analytics/average_mw_per_chunk.csv:

```
#This script takes in a floating point variable that represents a molecular weight average to determine which chunk in the conformer library most closely matches that value. The closest chunk index, average molecular weight, and different from input are printed to the standard output.

python
ginsparg_thymelab_thesis/conformer_library_analytics/find_chunk_with_closest_averag
e_mw_to_request.py 223.3865
```

For each sub-chunk, all conformers were extracted, converted to Rosetta params format, and organized into unique numbered test_params directories with up to 100 conformers per directory. The test_params directories were given a new specified address to be used on the OSG computational cluster. ginsparg_thymelab_thesis/prepare_ligands_for_discovery/prepare_whole_subchunk_for_discovery.py was used to automate these processes:

```
#This script works with a select sub-chunk, prepares Rosetta test_params directories for discovery, and then pushes them to a given bucket location.

#Argument 1 – The bucket location of the sub-chunk to prepare for discovery.

#Argument 2 – The top level of the bucket location to push all created test_params directories of conformers.

python
ginsparg_thymelab_thesis/prepare_ligands_for_discovery/prepare_whole_subchunk_for_d
iscovery.py
s3://ariosg/ligand_library/00297/for_s3/condensed_params_and_db_0.tar.gz
s3://ariosg/benchmarking_other_papers/ai_powered/ligand_inputs/
```

***REAL-M Discovery***

Compressed test_params directories were used as inputs for discovery. Anchor residues were selected based on residues highlighted in the respective publication. The submission scripts utilized the shell script ginsparg_thymelab_thesis/benchmarking_other_papers/discovery_on_prepared_test_params_directory.sh for identifying the best placements and ginsparg_thymelab_thesis/benchmarking_other_papers/discovery_on_prepared_test_params_directory_keep_placements.sh to run discovery and retain placements for comparison against any models that were available in the publication.

***Ranking of Placements***

After discovery was performed on the publication and decoy ligands, the simplified placement data was processed using the script ginsparg_thymelab_thesis/ benchmarking_other_papers/pull_data_files_and_rank_ligands.py. This script pulls down all placement data to determine where publication ligands rank against the decoy set on the metrics of free energy, total interaction count, and ratio of real motif-like interactions. The script creates a single csv file named paper_ligands_rank.csv:

```
#This script works with a select sub-chunk, prepares Rosetta test_params directories for discovery, and then pushes them to a given bucket location.

#Argument 1 – The bucket location of the publication output data

python ginsparg_thymelab_thesis/
benchmarking_other_papers/pull_data_files_and_rank_ligands.py
s3://ariosg/benchmarking_other_papers/ai_powered/
```

Ligand rankings were visualized in Jupyter notebooks in ginsparg_thymelab_thesis/jupyter_notebooks/figure_making/replicating_other_publications/good

***Native Placement Recovery***

Comparison of the REAL-M placements to published ligand orientations was only possible if the publication included an experimentally derived or modeled structure of the complex. For Gryniukova et al., we compared placements for Enamine drug Z26395438 from their placement file jm3c00128_si_002.pdb. For Sing, Li et al, we compared placements for Enamine drugs Z1907784975, Z2732986066, Z3343635604, Z4324535763, Z5185631889, Z5185631911, Z5348530222, Z5348530626, Z5348530683, Z5348530741. Atom names were manually mapped to match the atom names used by Rosetta so that RMSD could be calculated. The scripts ginsparg_thymelab_thesis/benchmarking_other_papers/4zzi_placement_analysis/rmsd_analysis.py and /5c8s_placement_analysis/rmsd_analysis.py were used to calculate the RMSD of the published placement against the REAL-M predictions.

Comparison of Human and Zebrafish GPCR Binding Pocket Conservation

This study was conducted using scripts in the github repository: thyme_lab_internship_2025.

***System Collection***

Human GPCR gene annotations with HUGO Gene Nomenclature Committee (HGNC)-approved nomenclature were obtained from the HGNC gene database. Receptors associated for olfaction or taste and Class B2 and C were filtered out. Human-zebrafish orthology data were retrieved from the Zebrafish Information Network (ZFIN). These datasets were cross-referenced to identify overlapping human genes and their corresponding zebrafish orthologs. Structural models for the identified human and zebrafish GPCRs were then downloaded from AlphaFold Protein Structure Database. 200 Class A, 12 Class B2, and 13 Class F GPCRs had Human and Zebrafish analog predicted structures downloaded from the AlphaFold Protein Structure Database. The file tm-align_alignments/outputs/ all_receptors_initial.csv lists all 225 human receptors initially considered for investigation.

***Structure Alignment***

The following reference GPCR structures were used to align human and zebrafish analogues of each receptor, per-class: Class A – 2RH1, Class B1 – 4K5Y, Class F – 4JKV. The script tm-align_alignments/scripts/make_alignments.py used TM-align software to align the human and zebrafish receptors against the reference for a constant alignment space.

***Binding Pocket Comparison***

For each system, binding pocket residues were selected using the script tm-align_alignments/scripts/get_pocket.py, which used pymol to select residues within 5 angstroms of the ligand in the reference for an automated conservative representation of residues around where a ligand would bind. The script tm-align_alignments/scripts/alignment_analysis_identity_and_similarity.py was used to parse each system set and compare binding pocket residues for identity and functional similarity. For each residue in the human receptor, the closest zebrafish residue was considered for comparison. The zebrafish residue was selected for comparison if it was within 2 angstroms of the human residue and was not being considered for comparison with another residue. For each system, counts were tallied for the number of residues in a position that were identical, residues that were functionally similar, were neither identical nor similar, or if there was not a zebrafish residue that was close enough to the human receptor residue. Percentages of each metric of the total residues was collected for reporting in results. Binding pocket residue percent identity, similarity, and mismatch did not include residues where there was not a receptor that was close enough to compare. 14 systems where >34% of all binding pocket residues did not have a human residue compare against a zebrafish residue due to being too far were removed from this study for having too many residues that could not be compared. 189 Class A, 11 Class B1, and 11 Class F systems were represented in the final findings of this study.

### Supplementary Figures

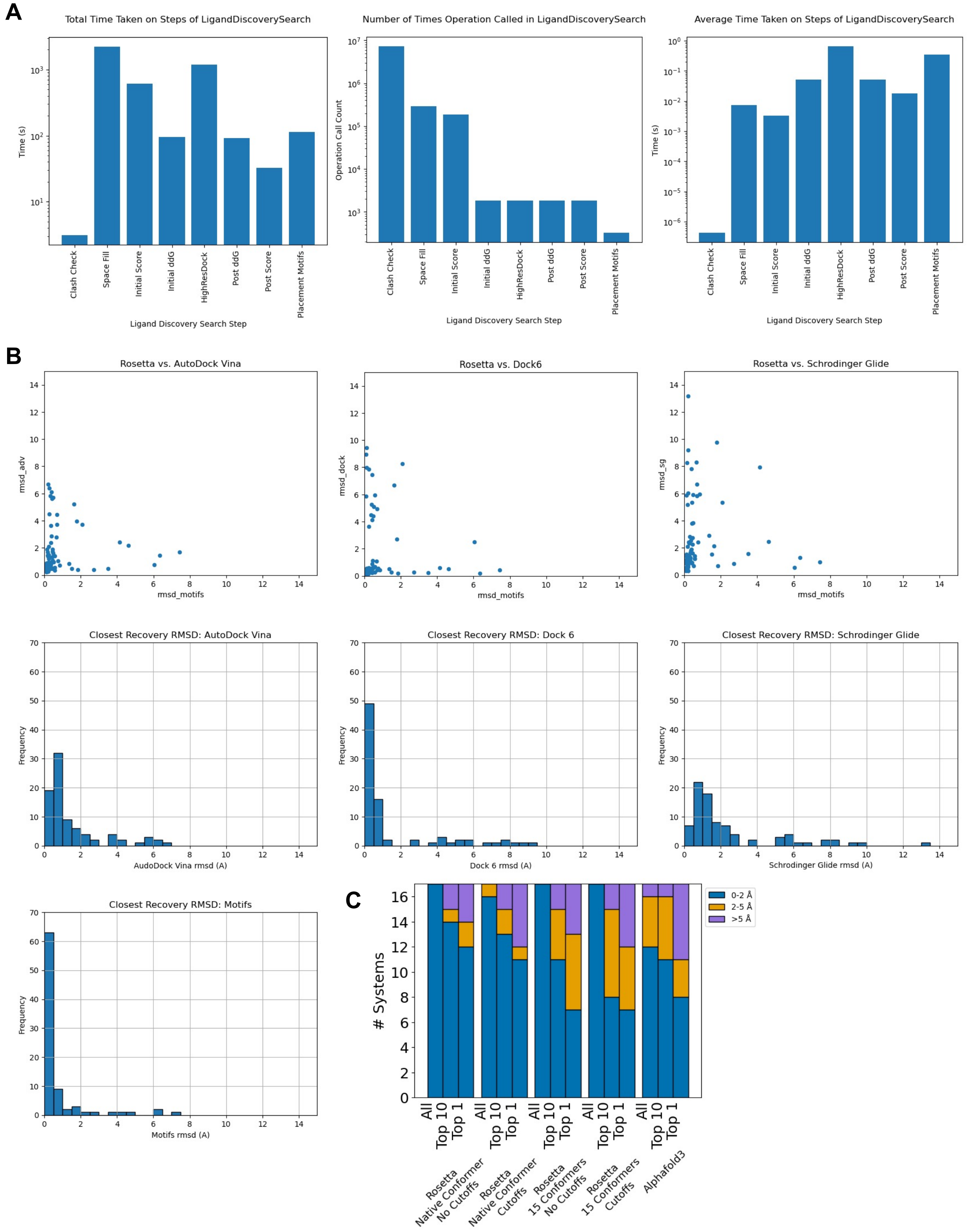


**Supplementary Figure 1. REAL-M benchmarking.** (**A**) Time cost analysis of REAL-M discovery. This analysis revealed the importance of fast-running selection steps in reducing the total runtime. An iteration of the protocol, modified to include time collection points, was performed on a set of 100 ligands from the set of ligands from agonist discovery against the active HCRTR2 (PDB 7L1U). The total time spent on processing and filtering steps, the number of times each step was called, and the average time taken for each step was derived as the number of step calls divided by total step time. Arguments and inputs for this study can be found in ginsparg_thymelab_thesis/jupyter_notebooks/figure_making/100_lig_timing_analysis_data. (**B**) Recovery benchmark of systems from DUD-E library across docking algorithms. Individual comparisons depicting the closest recovery from all placement attempts for each system for REAL-M compared to AutoDock Vina, DOCK 6, and Schrödinger-Maestro Glide. Schrödinger-Maestro Glide only returns a single placement. (**C**) AlphaFold 3 recovery of ligand placements in PDB structures from 2025. The selected protein-ligand pairings are too new to exist in the curated REAL-M motifs library or AlphaFold 3 database. Stacked bar plots representing the systems where the software's closest placement to the native placement by RMSD of heavy atoms was within 2 Å (blue), between 2-5 Å (orange), and beyond 5 Å (purple). The three comparisons are the closest recovery placement was taken out of all recovered placements, the top 10 placements by free energy, and the single top free energy placement. We compared REAL-M run with the native conformer included and our standard filtering cutoffs such as for clashes and space-fill, with native conformer and no filtering cutoffs, with 15 conformers generated with Conformator with and without filtering.

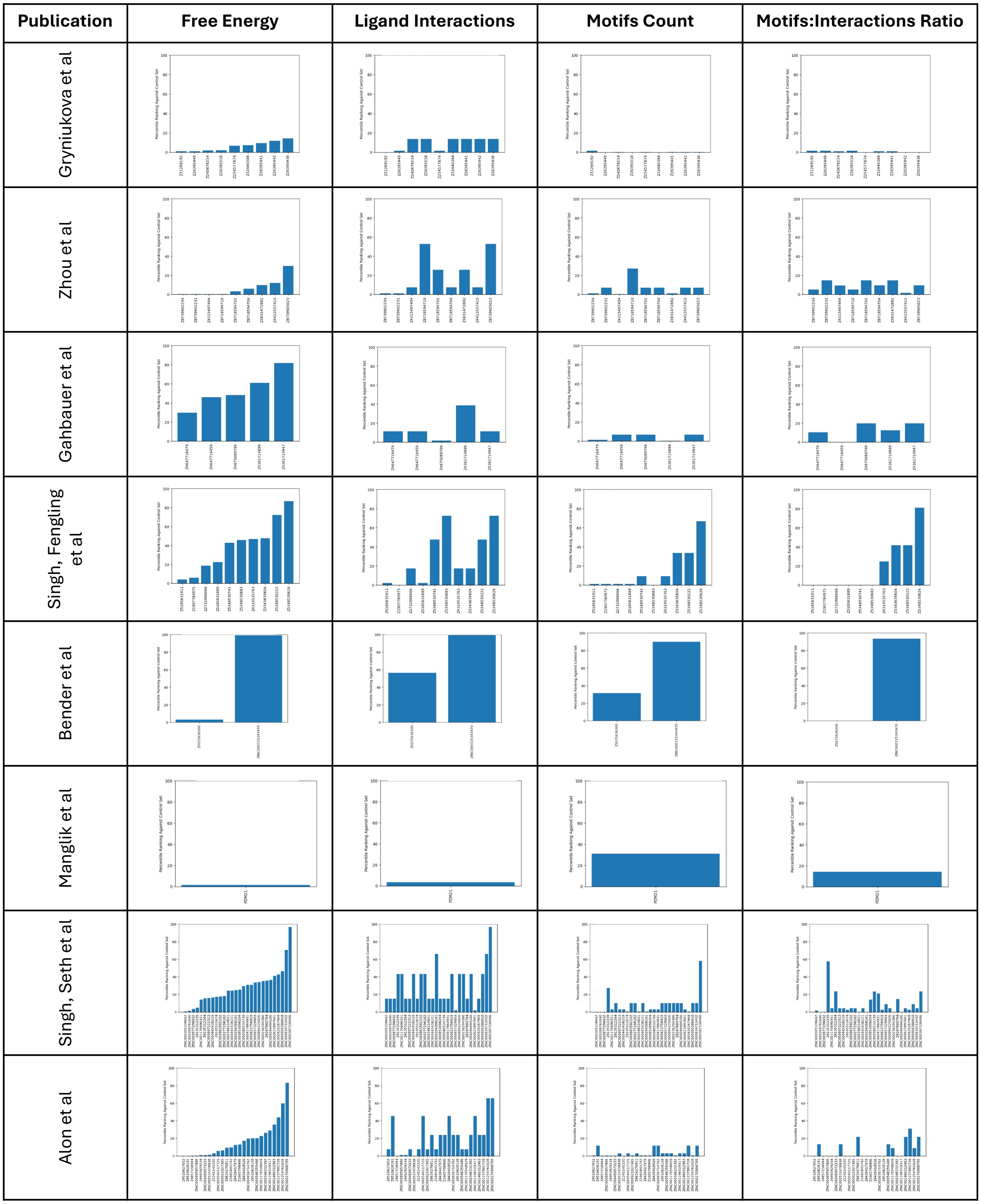


**Supplementary Figure 2. REAL-M discovery of selected ligands from other publications.** Each of these eight publications experimentally validated ligands that were selected from computational docking campaigns using Enamine virtual libraries. For each ligand, up to 15 conformers were generated and combined with 10,000 random ligands of a similar molecular weight. The following metrics were collected from each placement: system free energy, total protein-ligand interactions, real motif-like interactions, and ratio of real motif-like interactions to total interactions. Graphs show the percentile of the best placement from each publication in relation to the other 10,000 ligands.

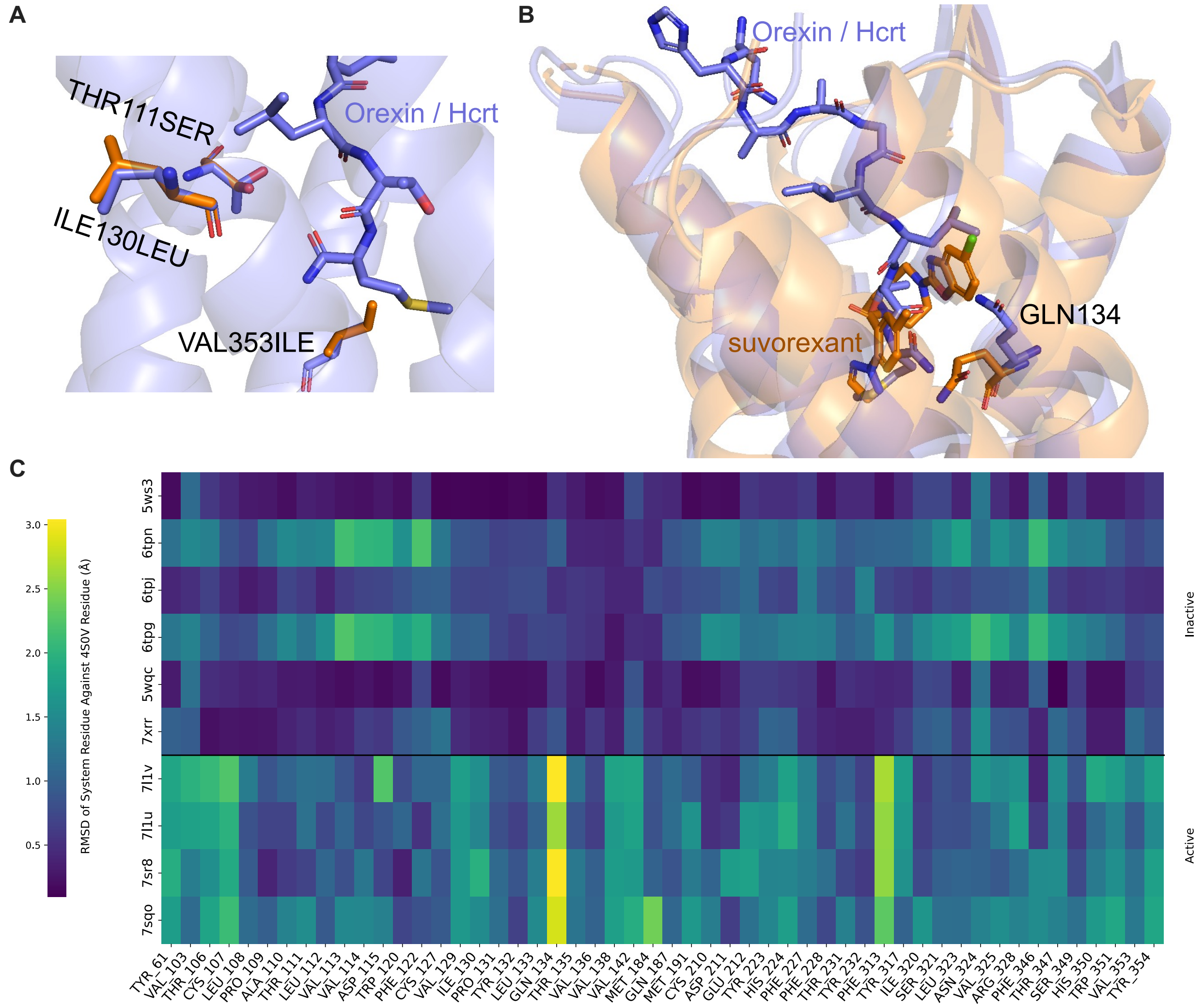


**Supplementary Figure 3. Shifts in the HCRTR2 binding pocket residues between the active and inactive states.** (**A**) Conservation of human HCRTR2 with the orexin peptide (7L1U, blue) compared to the zebrafish Hcrtr2 receptor. All binding pocket residues are identical except those shown here in orange and labeled. (**B**) Comparison of active (7L1U, blue) and inactive (4S0V, orange) structures, depicting the upward flip of GLN134 in the active state. (**C**) Four published systems of HCRTR2 in an agonist-bound state and six published systems of HCRTR2 in an antagonist-bound state were aligned against PDB 4S0V (HCRTR2-suvorexant) using the Pymol align function. Residues within 7 Å of suvorexant in 4S0V were selected. The RMSD of the residue heavy atoms for each residue in each system was calculated against the corresponding residue in 4S0V.

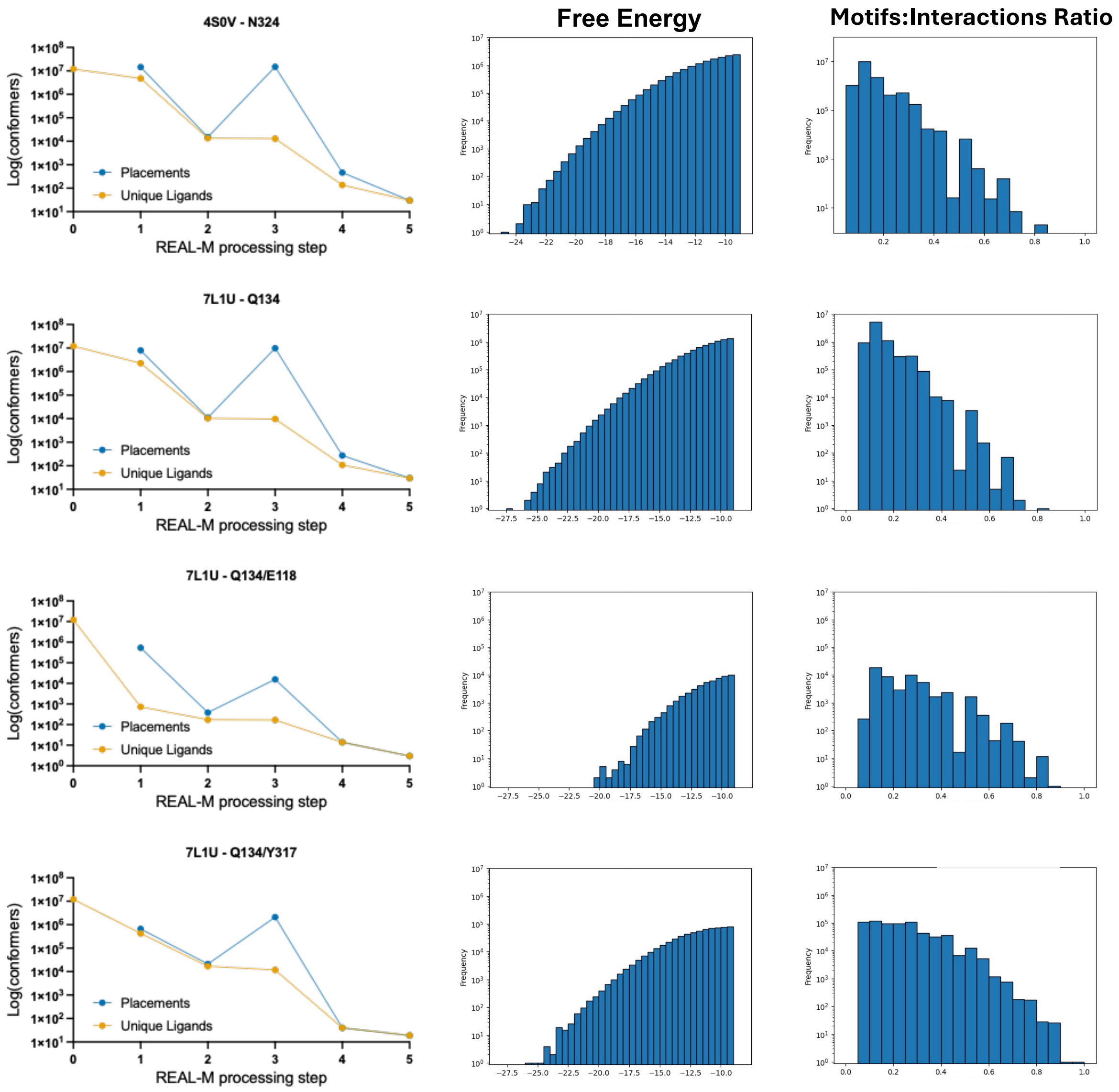


**Supplementary Figure 4. Progression and filtering at major steps in the REAL-M pipeline.** The line plots (left) plot the number of placement and corresponding unique ligands at the following key steps: Step 0: initial count of VAMS ShapeDB-selected conformers. Step 1: initial placements of ligand conformers. Step 2: filtering of initial placements by free energy or real interactions. Step 3: placements of ligands with expanded sets of up to 250 conformers. Step 4: filtering of expanded conformer set placements by free energy or motif-like interactions and conformer strain energy. Ligand placements at this stage were considered for manual review. Step 5: selected ligands successfully synthesized by Enamine. Histogram plots show the distribution of system free energy and ratio of interactions that can be observed in the motif library from the initial REAL-M placements (step 2). These metrics, in addition to several others detailed in the Supplementary Methods, were key to selecting ligands for refinement with the expanded set of up to 250 conformers. Two groups were selected for refinement: (1) stringent free energy threshold (≤ -16 REU) and lenient allowance for ligand-receptor interactions that are found in the PDB (≥ 0.25). (2) A lenient free energy threshold (≤ -9 REU) and stringent allowance for ligand-receptor interactions that are found in the PDB (≥ 0.5).

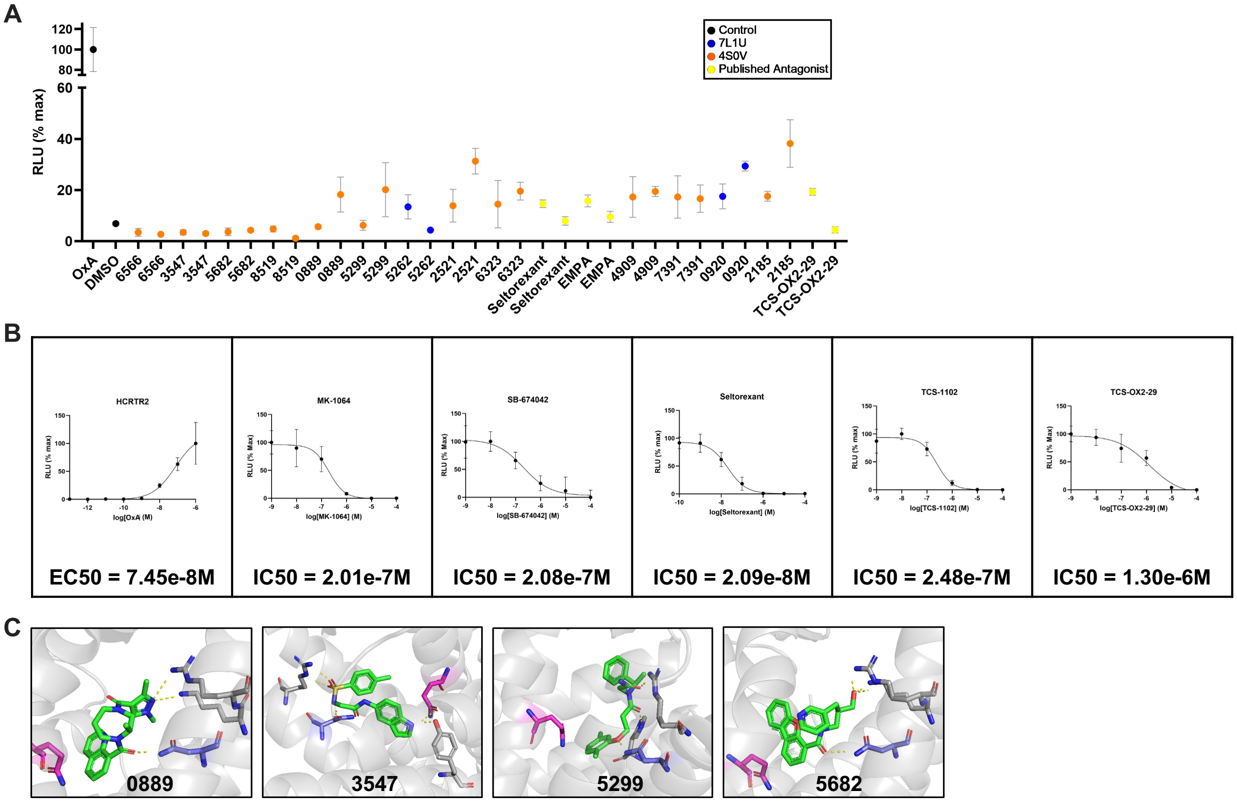


**Supplementary Figure 5. Additional PRESTO-Tango results for antagonists.** (**A**) Replication of antagonism for the top 13 newly identified antagonists (Fig. 2A), comparing original data (left) to the independent replication experiment (right). (**B**) Dose-response curves for OxA and published antagonists. The OxA curve is normalized to 100% RLU at 1e-6 M OxA. All antagonist curves are normalized to 100% RLU at 1e-7 M OxA. The EC50 or IC50 for each curve was calculated with GraphPad Prism using three parameter nonlinear regression (**C**) Predicted binding modes of experimental antagonists 0889, 3547, 5299, and 5682. Ligands are represented in green, GLN134 in magenta, and ASN324 blue. Pymol sessions are available on the linked Zenodo repository.

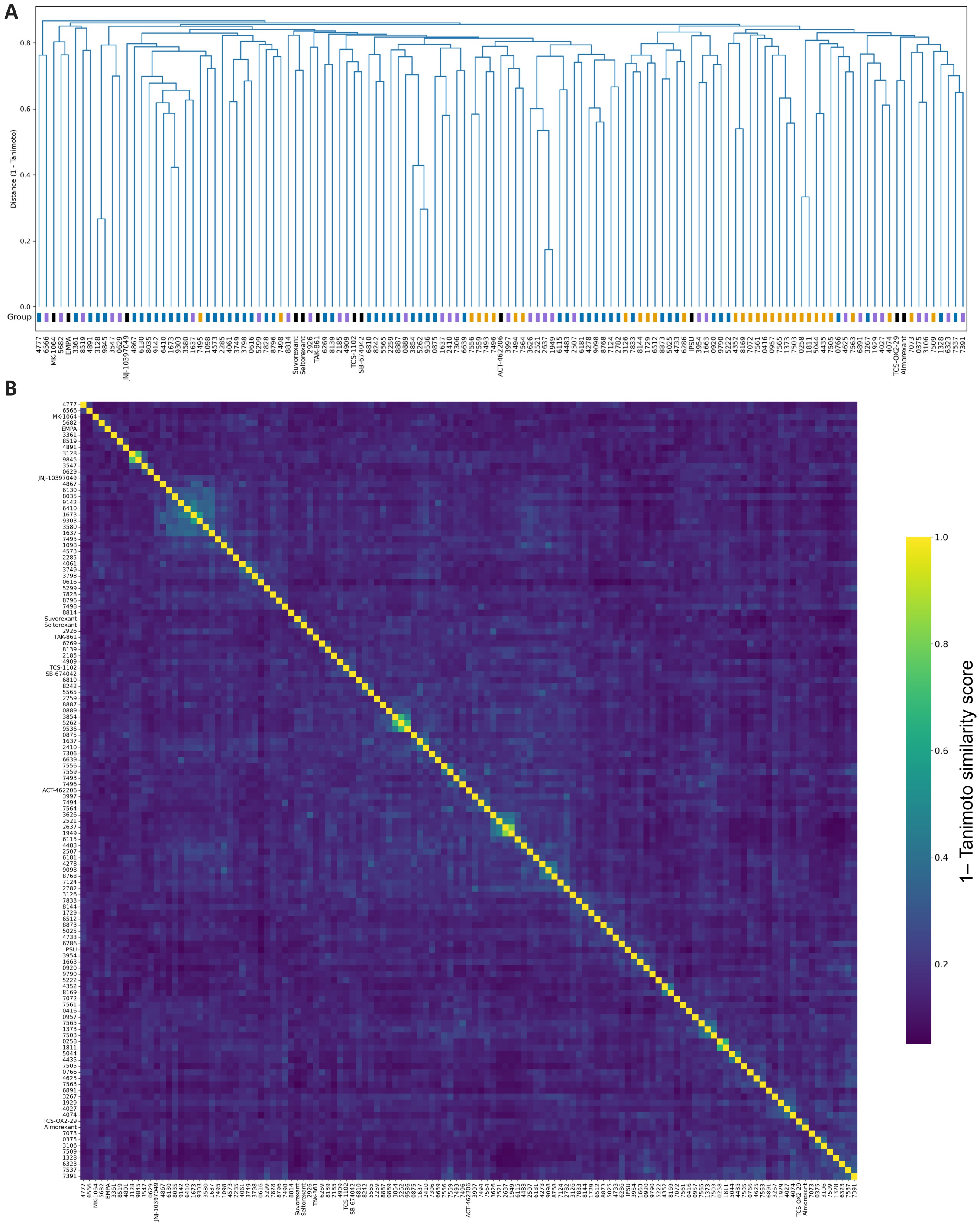


**Supplementary Figure 6. Chemical Similarity of New and Existing HCRTR2 Ligands.** (**A**) Molecules were clustered by inverted Tanimoto score similarity, derived by pairwise SMILES string comparisons using a radius a Morgan fingerprint radius of 2 and bit size of 2048. Colored group bars indicate whether the drug is published or which HCRTR2 PDB the experimental drug was discovered from. Coloring: 4S0V – purple, 7L1U – blue, 7SR8 – orange, published – black. (**B**) Heatmap representation for pairwise molecular structure similarity comparison that is displayed in Supplementary Figure 6A dendrogram. Similarity was determined by Tanimoto score, comparing drug SMILES strings using same fingerprint radius and bit size.

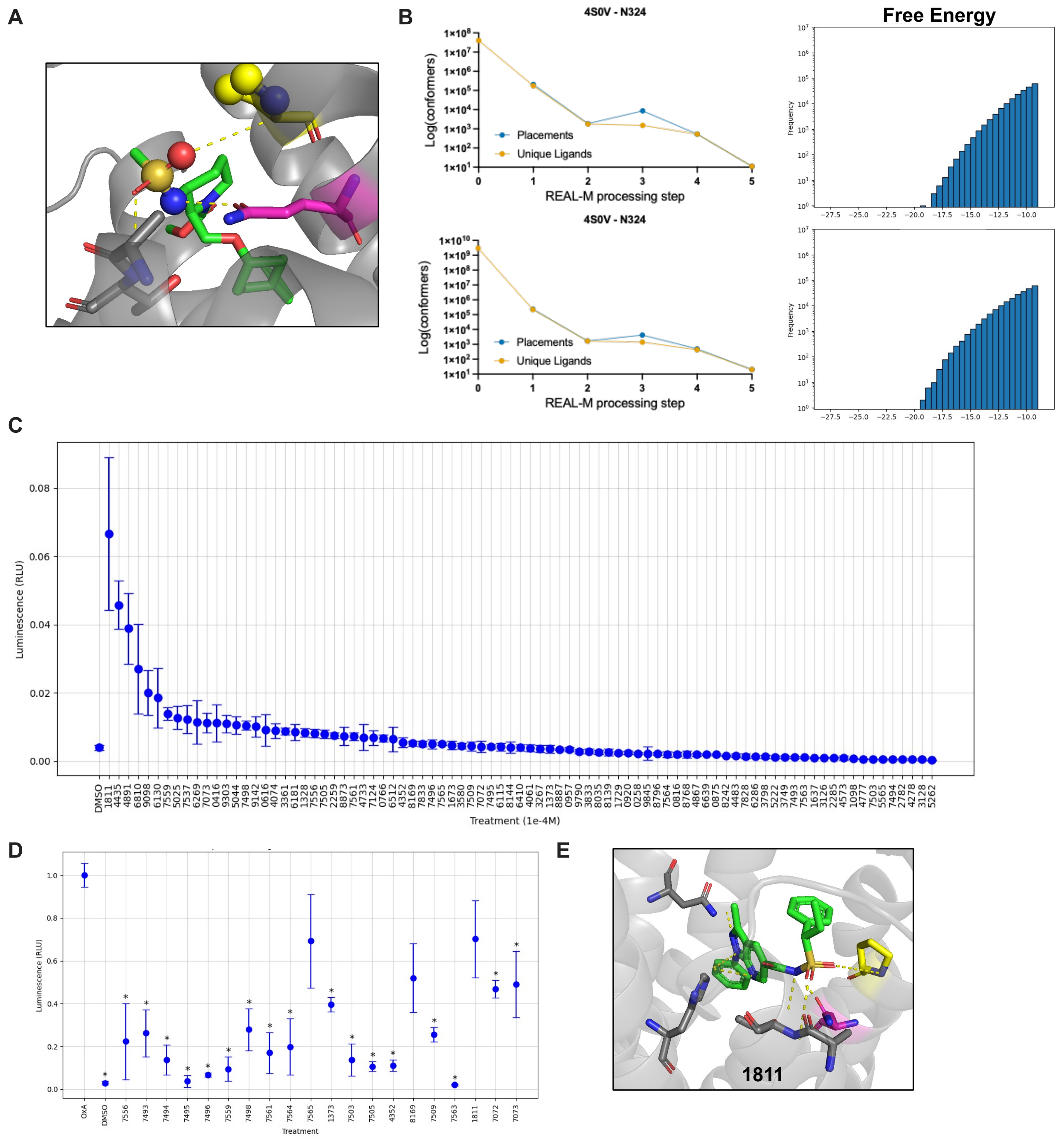


**Supplementary Figure 7. Chemical Similarity of New and Existing HCRTR2 Ligands.** (**A**) Highlighted interactions of sulfonamide group in TAK-925 (green) in PDB 7SR8. Motif atoms between TAK-925 sulfonamide and PRO131 that was used by Rosetta for discovery are represented by spheres. ALA110 and THR111 are represented in silver, PRO131 is represented in yellow, and GLN134 is represented in magenta. (**B**) Progression and filtering at major steps in the REAL-M pipeline, plotted similarly to Supplementary Figure 4. The results for the screen from 2.6B ligands are on top and from 64.9B ligands are on the bottom. The line plots (left) plot the number of placement and corresponding unique ligands at key steps. Histogram plots (right) show the distribution of system free energies from initial placements before refinement. (**C**) PRESTO-Tango screen for agonists at 1e-4 M. Relative luminescence values are normalized against the TAK-861 positive control, which has similar agonist activity to OxA, set equal to 1. The TAK-861 positive control is omitted from plots to better visualize the luminescence magnitude. (**D**) PRESTO-Tango screen for sulfonamide-containing ligands as antagonists. Relative luminescence values are normalized against the OxA positive control, set equal to 1. (**E**) Predicted binding mode of a newly identified agonist.

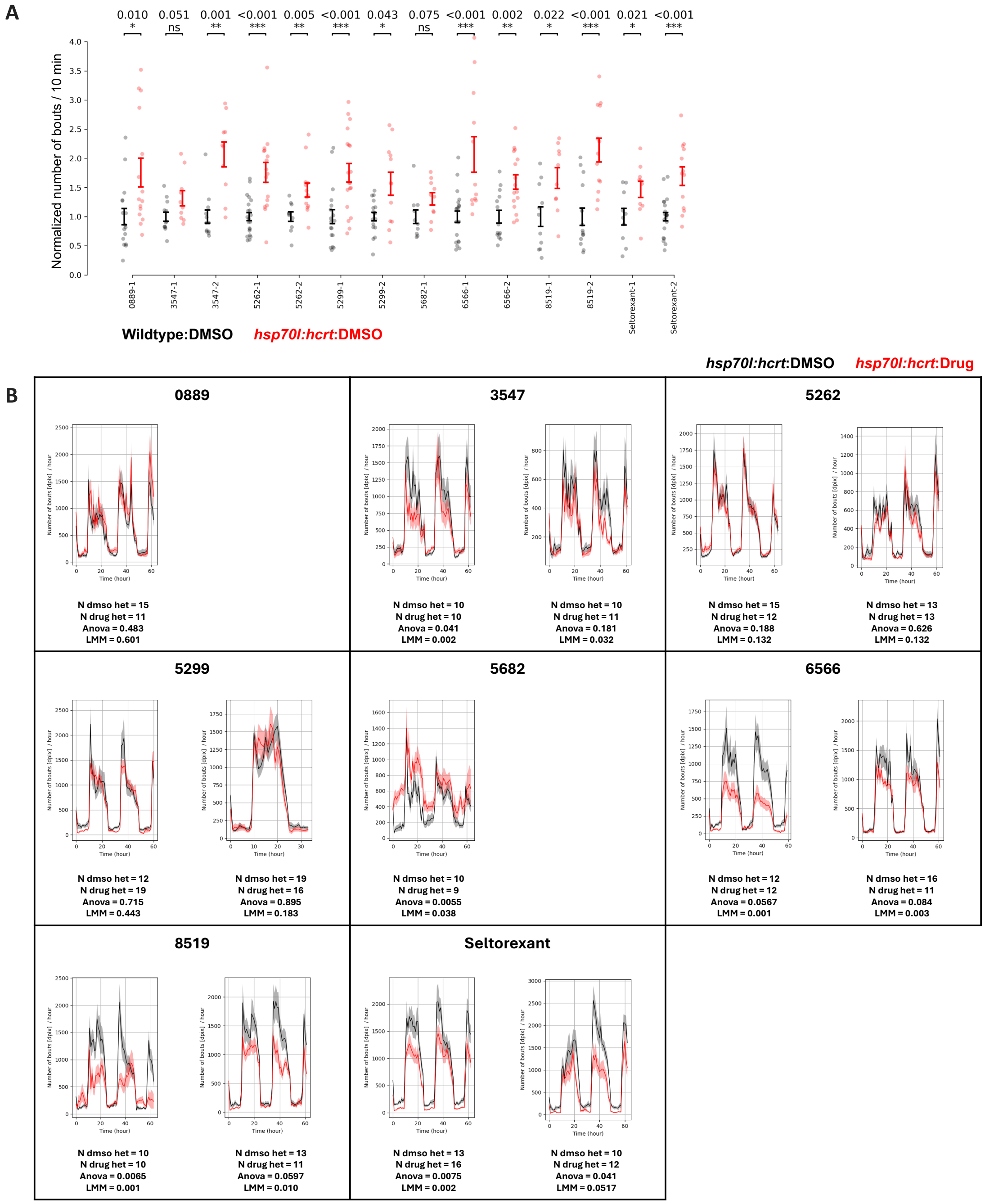


**Supplementary Figure 8. Modulation of activity of hypocretin-overexpressing zebrafish with receptor antagonists.** (**A**) Validation of heat shock response inducing increased hypocretin production in *hsp70l:hcrt* heterozygous fish (red) compared to wild-type siblings (black) on the first experiment day (9:00-23:00, 5 dpf) following heat shock All depicted fish were treated with DMSO. The number of movement bouts per fish was binned per 600 seconds, then averaged for the timespan measured for the plot. Movement bout count was normalized against the average bout count from the control group per experiment to compare between experiments. (**B**) Example multi-day activity profiles of larval zebrafish following heat shock and corresponding replicates. The zebrafish experience heat shock on the afternoon of 4 dpf and the activity tracking begins that night. The significance values are for the first experiment day.

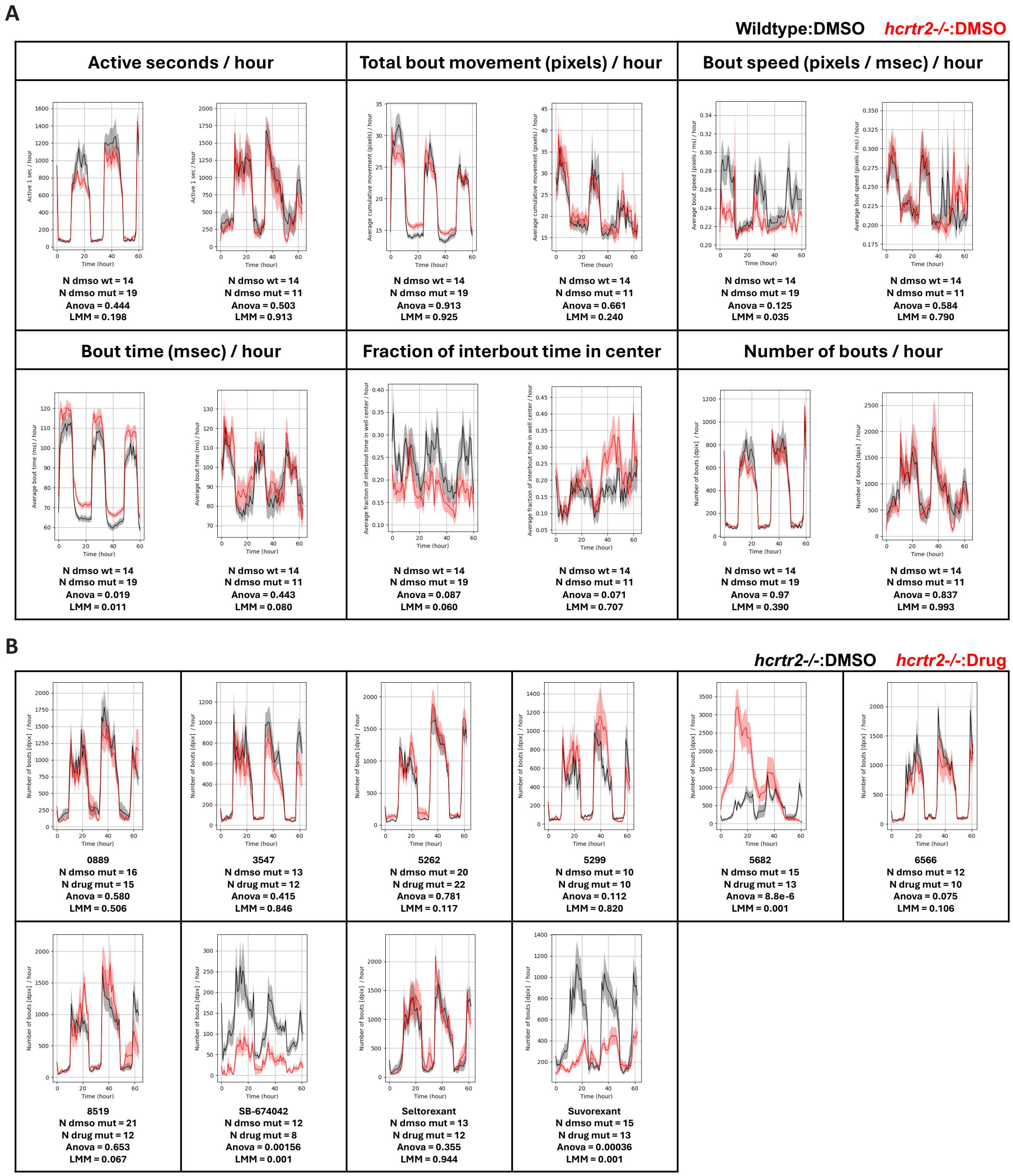


**Supplementary Figure 9. Determining of off-target behavioral effects of experimental and commercially available drugs.** (**A**) Lack of a consistent, strong phenotype across multiple behavioral measures comparing homozygous *hcrtr2* mutants to wild-type siblings. (**B**) Example multi-day activity profiles of *hcrtr2* mutant larval zebrafish, comparing drug-treated to DMSO-treated. All drugs were assessed at 20 µM with the exception of suvorexant, which was tested at 5 µM because it was lethal at higher doses. The following molecules that were found to be completely lethal at 20 µM and at 4 dpf (data not shown): IPSU (MedChemExpress - cat # HY-13796), JNJ-10397049 (MedChemExpress - cat # HY-10896), and Almorexant (MedChemExpress - cat # HY-10805).

### Supplementary Tables

**Supplementary Table 1. Motif library composition by amino acid type.** Counts of each motif were derived after removing motifs that were considered duplicate, based on having the same residue, atom types, and being within the cutoff RMSD of 0.4 Å and 0.3 radians. Glycine is not included, as it lacks side chain atoms to form a motif.

| Amino Acid | Motif Count |
|---|---|
| **Polar Uncharged** | |
| ASN | 74,601 |
| CYS | 25,371 |
| GLN | 68,200 |
| SER | 60,693 |
| THR | 74,430 |
| TYR | 126,514 |
| **Positively Charged** | |
| ARG | 168,379 |
| HIS | 88,900 |
| LYS | 135,648 |
| **Negatively Charged** | |
| ASP | 68,826 |
| GLU | 76,226 |
| **Nonpolar Aliphatic** | |
| ALA | 34,383 |
| ILE | 90,035 |
| LEU | 158,623 |
| MET | 65,611 |
| PRO | 46,636 |
| VAL | 83,630 |
| **Nonpolar Aromatic** | |
| PHE | 130,108 |
| TRP | 73,401 |
| **Total** | **1,650,215** |

**Supplementary Table 2. Antagonist molecular shapes used as inputs for ShapeDB.** Three molecules, all from published structures bound to HCRTR2, that were used for antagonist discovery with REAL-M.

| **Antagonists** | | | |
|---|---|---|---|
| Ligand Name | Chemical Formula | 2D Structure | 3D Structure |
| EMPA | $C_{23}H_{26}N_4O_4S$ | 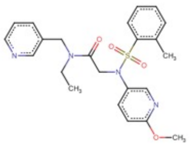 | 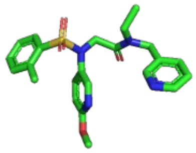 |
| HTL6641 | $C_{20}H_{15}F_3N_4O_5S$ | 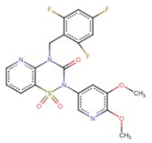 | 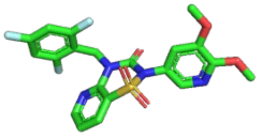 |
| Lemborexant | $C_{22}H_{20}F_2N_4O_2$ | 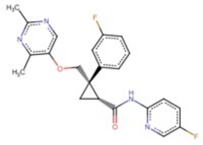 | 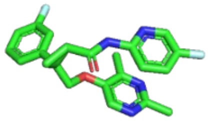 |

**Supplementary Table 3. Agonist molecular shapes used as inputs for ShapeDB.** Four truncations of OxA and four truncations of OxB, based on the 4-7 terminal residues that are in the binding pocket of 7L1U.

| Agonists | | | |
|---|---|---|---|
| Ligand Name | Residue Sequence | 2D Structure | 3D Structure |
| OxA_4 | ILTL | 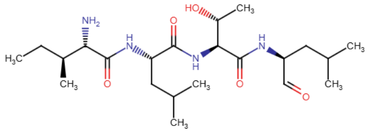 | 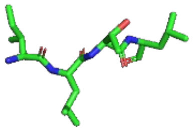 |
| OxA_5 | GILTL | 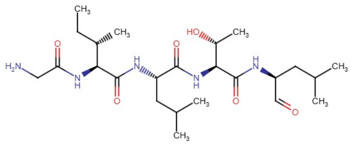 | 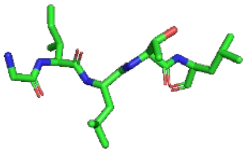 |
| OxA_6 | AGILTL | 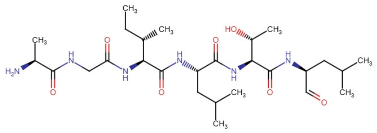 | 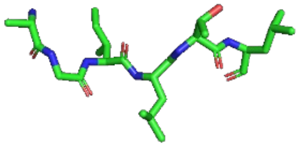 |
| OxA_7 | AAGILTL | 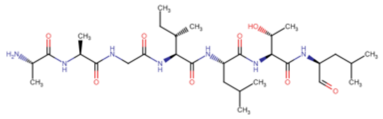 | 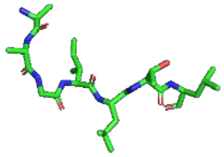 |
| OxB_4 | ILTM | 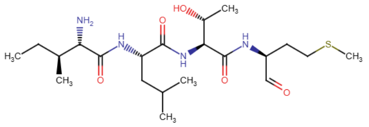 | 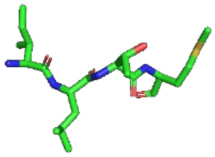 |
| OxB_5 | GILTM | 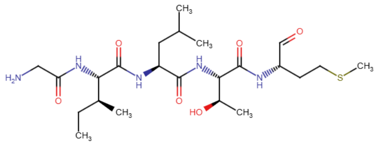 | 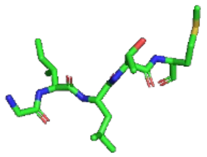 |
| OxB_6 | AGILTM | 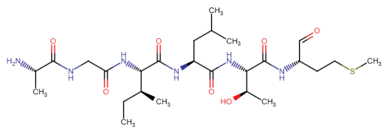 | 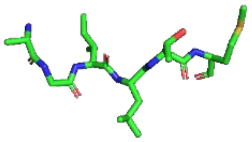 |
| OxB_7 | AAGILTM | 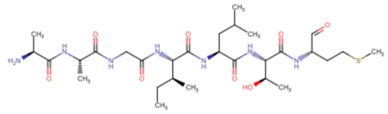 | 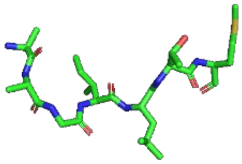 |

**Supplementary Table 4. Ligands or conformer count at primary steps of the REAL-M discovery pipeline.** Each column summarizes counts at a step in the pipeline, with the first column providing the starting number of conformers for each discovery run. In the case of 7SR8 discovery, conformers were generated in runtime, and up to 10 conformers were generated. The second column lists the number of placements generated by the initial screen and corresponding number of unique ligands. The third column lists the number of placements corresponding unique ligands that pass filters for free energy or realistic interactions from the expanded conformer screen. Values in parenthesis depict the breakdown of systems that pass a filter. The fourth column lists the number of placements generated by the expanded conformer screen and corresponding number of unique ligands. The fifth column lists the number of placements corresponding unique ligands that pass filters for free energy or realistic interactions from the expanded conformer screen. The sixth column represents the ligands passed manual selection review and could successfully be synthesized by Enamine.

| | | **Starting Library**<br>Conformers | **Initial Screen Placements**<br>Placements / Ligands | **Initial Screen Selected Placements**<br>Placements (low ddG + real interactions) / Ligands (low ddG + real interactions) | **Refined Screen Placements**<br>Placements (low ddG + real interactions) / Ligands (low ddG + real interactions) | **Ligands For Manual Review**<br>Placements (low ddG + real interactions) / Ligands (low ddG + real interactions) | **Ligands Selected and Synthesized**<br>Ligands (low ddG + real interactions) |
|---|---|---|---|---|---|---|---|
| 4S0V | ASN324 | 11,998,267 | 14,532,857 / 4,754,346 | 14,926 (7,669 + 7,257) / 13,461 (6,598 + 6,863) | 14,853,928 (13,548,099 + 14,853,928) / 12,999 (6,207 + 6,792) | 461 (364 + 97) / 137 (104 + 33) | 30 |
| 7L1U | GLN134 | 12,035,273 | 7,962,840 / 2,286,028 | 11,108 (7,382 + 3,726) / 10,408 (6,837 + 3,572) | 9,923,178 (6,949,767 + 2,973,411) / 9,716 (6733 + 2,983) | 273 (138 +135) / 108 (52 + 56) | 30 |
| | GLN134 + GLU118 | 12,035,273 | 538,345 / 734 | 386 (0 + 386) / 172 (0 + 172) | 15,625 (0 + 15,625) / 168 (0 +168) | 14 (0 + 14) / 14 (0 + 14) | 3 |
| | GLN134 + TYR317 | 12,035,273 | 662,332 / 426,881 | 20,896 (471 + 20,425) / 16,983 (437 + 16,546) | 2,114,139 (40,069 + 2,074,070) / 11,743 (388 + 11,355) | 40 (6 + 34) / 40 (6 + 34) | 19 |
| 7SR8 | GLN134 2.6B Library | 40,222,330 x 10 | 206,855 / 175,521 | 1,845 / 1,762 | 8,601 / 1,505 | 530 / 530 | 11 |
| | GLN134 64.9B Library | 2,986,380,814 x 10 | 237,958 / 222,138 | 1,692 / 1,672 | 4,279 / 1,411 | 498 / 436 | 20 |

**Supplementary Table 5. Successfully synthesized ligands for 4S0V.** Ligands selected from the Enamine catalog identified by REAL-M. Each ligand is depicted along with its corresponding Enamine ID, SMILES string, chemical formula, and mass in atomic mass units. Open Babel was used to derive ligand SMILES strings from their original .sdf format. The Protein Data Bank Chemical Sketch Tool was used to draw each ligand from its SMILES string.

| | | | | | |
|---|---|---|---|---|---|
| 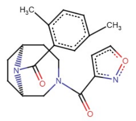<br>PV-000555062637<br>Cc1ccc(C)c(C(=O)N2[C@H]3CC[C@@H]2CN(C(=O)c2ccon2)CC3)c1<br>C20H23N3O3<br>353.41 a.m.u. | 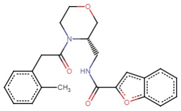<br>PV-000574352410<br>Cc1ccccc1CC(=O)N1CCOC[C@@H]1CNC(=O)c1cc2ccccc2o1<br>C23H24N2O4<br>392.45 a.m.u. | 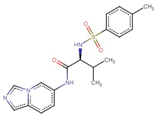<br>PV-002221910629<br>Cc1ccc(S(=O)(=O)N[C@H](C(=O)Nc2ccc3cncn3c2)C(C)C)cc1<br>C19H22N4O3S<br>386.47 a.m.u. | 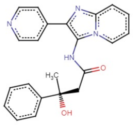<br>PV-002353454027<br>C[C@@](O)(CC(=O)Nc1c(-c2ccncc2)nc2ccccn12)c1ccccc1<br>C22H20N4O2<br>372.42 a.m.u. | 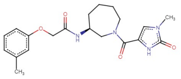<br>PV-002657363997<br>Cc1cccc(OCC(=O)N[C@H]2CCCCN(C(=O)c3cn(C)c(=O)[nH]3)C2)c1<br>C20H26N4O4<br>386.44 a.m.u. | 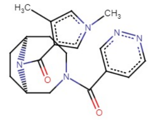<br>PV-002738102521<br>Cc1cn(C)cc1C(=O)N1[C@H]2CC[C@@H]1CN(C(=O)c1ccnnc1)CC2<br>C19H23N5O2<br>353.42 a.m.u. |
| 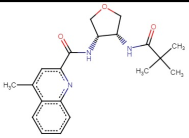<br>PV-003064601637<br>Cc1cc(C(=O)N[C@@H]2COC[C@@H]2NC(=O)C(C)(C)C)nc2ccccc12<br>C20H25N3O3<br>355.43 a.m.u. | 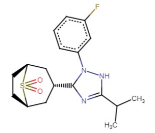<br>PV-003960018814<br>CC(C)C1=NC([C@H]2C[C@@H]3CC[C@H](C2)S3(=O)=O)N(c2cccc(F)c2)N1<br>C18H24FN3O2S<br>365.46 a.m.u. | 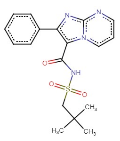<br>PV-004156311929<br>CC(C)(C)CS(=O)(=O)NC(=O)c1c(-c2ccccc2)nc2ncccn12<br>C18H20N4O3S<br>372.44 a.m.u. | 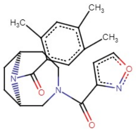<br>PV-004632151949<br>Cc1cc(C)c(C(=O)N2[C@H]3CC[C@@H]2CN(C(=O)c2ccon2)CC3)cc1C<br>C21H25N3O3<br>367.44 a.m.u. | 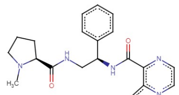<br>PV-005091654909<br>CN1CCC[C@H]1C(=O)NC[C@H](NC(=O)c1nccn2nccc12)c1ccccc1<br>C21H24N6O2<br>392.45 a.m.u. | 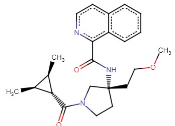<br>PV-005143787306<br>COCC[C@@]1(NC(=O)c2ncccc3ccccc23)CCN(C(=O)[C@@H]2[C@@H](C)[C@H]2C)C1<br>C23H29N3O3<br>395.49 a.m.u. |
| 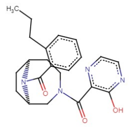<br>PV-005191963626<br>CCCc1ccccc1C(=O)N1[C@H]2CC[C@@H]1CN(C(=O)c1nccnc1O)CC2<br>C22H26N4O3<br>394.47 a.m.u. | 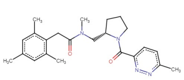<br>PV-005396532185<br>Cc1cc(C)c(CC(=O)N(C)C[C@@H]2CCCN2C(=O)c2ccc(C)nn2)c(C)c1<br>C23H30N4O2<br>394.51 a.m.u. | 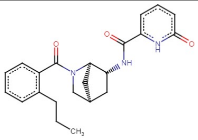<br>PV-005932782926<br>CCCc1ccccc1C(=O)N1C[C@@H]2C[C@@H](NC(=O)c3cccc(=O)[nH]3)[C@H]1C2<br>C22H25N3O3<br>379.45 a.m.u. | 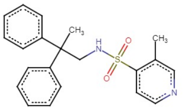<br>PV-006204130375<br>Cc1cnccc1S(=O)(=O)NCC(C)(c1ccccc1)c1ccccc1<br>C21H22N2O2S<br>366.48 a.m.u. | 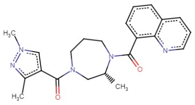<br>PV-006254270889<br>Cc1nn(C)cc1C(=O)N1CCCN(C(=O)c2cccc3cccnc23)[C@H](C)C1<br>C22H25N5O2<br>391.47 a.m.u. | 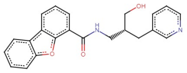<br>PV-006439055682<br>O=C(NC[C@H](CO)Cc1cccnc1)c1cccc2c1oc1ccccc12<br>C22H20N2O3<br>360.41 a.m.u. |
| 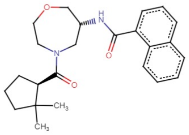<br>PV-006512772507<br>CC1(C)CCC[C@H]1C(=O)N1CCOC[C@H](NC(=O)c2cccc3ccccc23)C1<br>C24H30N2O3<br>394.51 a.m.u. | 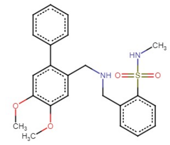<br>Z1141296323<br>CNS(=O)(=O)c1ccccc1CNCc1cc(OC)c(OC)cc1-c1ccccc1<br>C23H26N2O4S<br>426.53 a.m.u. | 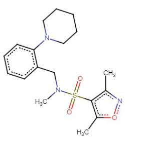<br>Z133963954<br>Cc1noc(C)c1S(=O)(=O)N(C)Cc1ccccc1N1CCCCC1<br>C18H25N3O3S<br>363.47 a.m.u. | 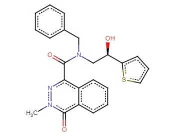<br>Z1393238519<br>Cn1nc(C(=O)N(Cc2ccccc2)C[C@@H](O)c2cccs2)c2ccccc2c1=O<br>C23H21N3O3S<br>419.50 a.m.u. | 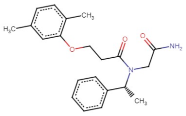<br>Z1410815299<br>Cc1ccc(C)c(OCCC(=O)N(CC(N)=O)[C@H](C)c2ccccc2)c1<br>C21H26N2O3<br>354.44 a.m.u. | 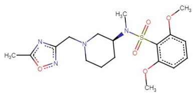<br>Z1725521663<br>COc1cccc(OC)c1S(=O)(=O)N(C)[C@H]1CCCN(Cc2noc(C)n2)C1<br>C18H26N4O5S<br>410.49 a.m.u. |
| 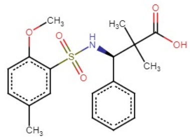<br>Z1738787391<br>COc1ccc(C)cc1S(=O)(=O)N[C@H](c1ccccc1)C(C)(C)C(=O)O<br>C19H23NO5S<br>377.45 a.m.u. | 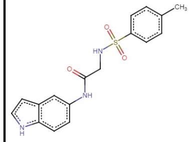<br>Z2169713547<br>Cc1ccc(S(=O)(=O)NCC(=O)Nc2ccc3[nH]ccc3c2)cc1<br>C17H17N3O3S<br>343.40 a.m.u. | 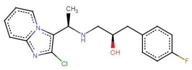<br>Z3360656891<br>C[C@@H](NC[C@H](O)Cc1ccc(F)cc1)c1c(Cl)nc2ccccn12<br>C18H19ClFN3O<br>347.81 a.m.u. | 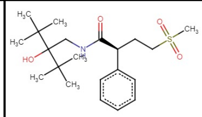<br>Z3914924625<br>CC(C)(C)C(O)(CNC(=O)[C@@H](CCS(C)(=O)=O)c1ccccc1)C(C)(C)C<br>C21H35NO4S<br>397.57 a.m.u. | 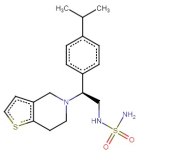<br>Z741186566<br>CC(C)c1ccc([C@@H](CNS(N)(=O)=O)N2CCc3sccc3C2)cc1<br>C18H25N3O2S2<br>379.54 a.m.u. | 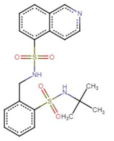<br>[illegible]<br>CC(C)(C)NS(=O)(=O)c1ccccc1CNS(=O)(=O)c1cccc2cnccc12<br>C20H23N3O4S2<br>433.54 a.m.u. |

**Supplementary Table 6. Successfully synthesized ligands for 7L1U.** Ligands selected from the Enamine catalog identified by REAL-M. Each ligand is depicted along with its corresponding Enamine ID, SMILES string, chemical formula, and mass in atomic mass units. The two molecules are the end are the orthologs of 5262, and the red circles represent the parts of the molecules that were modified to eliminate key interactions.

| | | | | | |
|---|---|---|---|---|---|
| 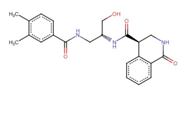<br>PV-001123528796<br>Cc1ccc(C(=O)NC[C@H](CO)NC(=O)[C@H]2CNC(=O)c3ccccc32)cc1C<br>$C_{22}H_{25}N_3O_4$<br>395.45 a.m.u. | 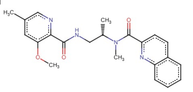<br>PV-001569394061<br>COc1cc(C)cnc1C(=O)NC[C@H](C)N(C)C(=O)c1ccc2ccccc2n1<br>$C_{22}H_{24}N_4O_3$<br>392.45 a.m.u. | 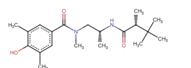<br>PV-003145840616<br>Cc1cc(C(=O)N(C)C[C@@H](C)NC(=O)[C@H](C)C(C)(C)C)cc(C)c1O<br>$C_{20}H_{32}N_2O_3$<br>348.48 a.m.u. | 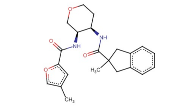<br>PV-004504869303<br>Cc1coc(C(=O)N[C@H]2COCC[C@H]2NC(=O)C2(C)Cc3ccccc3C2)c1<br>$C_{22}H_{26}N_2O_4$<br>382.45 a.m.u. | 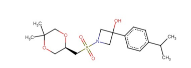<br>PV-005718134777<br>CC(C)c1ccc(C2(O)CN(S(=O)(=O)C[C@H]3COC(C)(C)CO3)C2)cc1<br>$C_{19}H_{29}NO_5S$<br>383.50 a.m.u. | 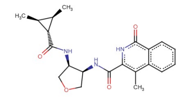<br>PV-006130153580<br>Cc1c(C(=O)N[C@H]2COC[C@H]2NC(=O)[C@@H]2[C@@H](C)[C@H]2C)[nH]c(=O)c2ccccc12<br>$C_{21}H_{25}N_3O_4$<br>383.44 a.m.u. |
| 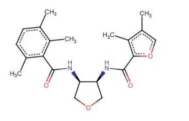<br>PV-006544139142<br>Cc1coc(C(=O)N[C@H]2COC[C@H]2NC(=O)c2c(C)ccc(C)c2C)c1C<br>$C_{21}H_{26}N_2O_4$<br>370.44 a.m.u. | 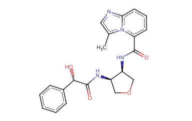<br>PV-006612138035<br>Cc1cnc2cccc(C(=O)N[C@H]3COC[C@H]3NC(=O)[C@@H](O)c3ccccc3)n12<br>$C_{21}H_{22}N_4O_4$<br>394.42 a.m.u. | 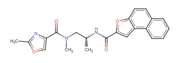<br>PV-006614933798<br>Cc1nc(C(=O)N(C)C[C@@H](C)NC(=O)c2cc3c(ccc4ccccc43)o2)co1<br>$C_{22}H_{21}N_3O_4$<br>391.42 a.m.u. | 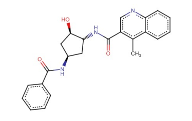<br>PV-006621251098<br>Cc1c(C(=O)N[C@@H]2C[C@@H](NC(=O)c3ccccc3)C[C@H]2O)cnc2ccccc12<br>$C_{23}H_{23}N_3O_3$<br>389.45 a.m.u. | 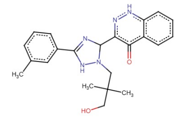<br>PV-006634533361<br>Cc1cccc(C2=NC(c3n[nH]c4ccccc4c3=O)N(CC(C)(C)CO)N2)c1<br>$C_{22}H_{25}N_5O_2$<br>391.47 a.m.u. | 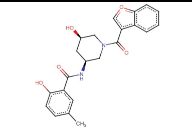<br>PV-006645844483<br>Cc1ccc(O)c(C(=O)N[C@H]2C[C@@H](O)CN(C(=O)c3coc4ccccc34)C2)c1<br>$C_{22}H_{22}N_2O_5$<br>394.42 a.m.u. |
| 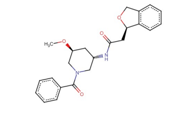<br>PV-006655328768<br>CO[C@H]1C[C@H](NC(=O)C[C@H]2OCc3ccccc32)CN(C(=O)c2ccccc2)C1<br>$C_{23}H_{26}N_2O_4$<br>394.46 a.m.u. | 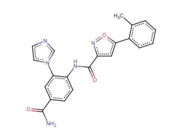<br>PV-006666022285<br>Cc1ccccc1-c1cc(C(=O)Nc2ccc(C(N)=O)cc2-n2ccnc2)no1<br>$C_{21}H_{17}N_5O_3$<br>387.39 a.m.u. | 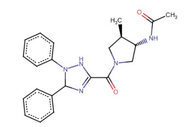<br>PV-006667477124<br>CC(=O)N[C@@H]1CN(C(=O)C2=NC(c3ccccc3)N(c3ccccc3)N2)C[C@H]1C<br>$C_{22}H_{25}N_5O_2$<br>391.47 a.m.u. | 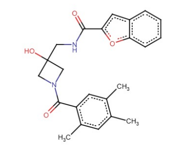<br>PV-006668031637<br>Cc1cc(C)c(C(=O)N2CC(O)(CNC(=O)c3cc4ccccc4o3)C2)cc1C<br>$C_{23}H_{24}N_2O_4$<br>392.45 a.m.u. | 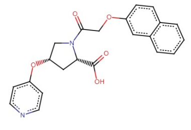<br>PV-006669822259<br>O=C(O)[C@@H]1C[C@H](Oc2ccncc2)CN1C(=O)COc1ccc2ccccc2c1<br>$C_{22}H_{20}N_2O_5$<br>392.40 a.m.u. | 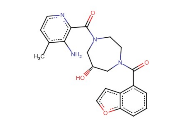<br>PV-006680046115<br>Cc1ccnc(C(=O)N2CCN(C(=O)c3cccc4occc34)C[C@H](O)C2)c1N<br>$C_{21}H_{22}N_4O_4$<br>394.42 a.m.u. |
| 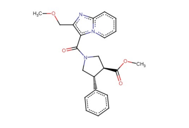<br>PV-006687433267<br>COCc1nc2ccccn2c1C(=O)N1C[C@@H](C(=O)OC)[C@H](c2ccccc2)C1<br>$C_{22}H_{23}N_3O_4$<br>393.43 a.m.u. | 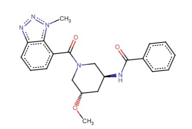<br>PV-006688879098<br>CO[C@H]1C[C@H](NC(=O)c2ccccc2)CN(C(=O)c2cccc3nnn(C)c23)C1<br>$C_{21}H_{23}N_5O_3$<br>393.44 a.m.u. | 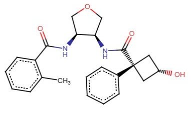<br>PV-006689746410<br>Cc1ccccc1C(=O)N[C@H]1COC[C@H]1NC(=O)[C@]1(c2ccccc2)C[C@H](O)C1<br>$C_{23}H_{26}N_2O_4$<br>394.46 a.m.u. | 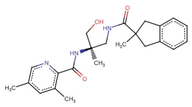<br>PV-006695134867<br>Cc1cnc(C(=O)N[C@@](C)(CO)CNC(=O)C2(C)Cc3ccccc3C2)c(C)c1<br>$C_{23}H_{29}N_3O_3$<br>395.49 a.m.u. | 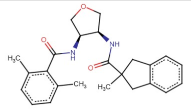<br>PV-006700551673<br>Cc1cccc(C)c1C(=O)N[C@H]1COC[C@H]1NC(=O)C1(C)Cc2ccccc2C1<br>$C_{24}H_{28}N_2O_3$<br>392.42 a.m.u. | 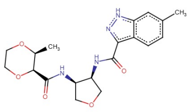<br>PV-006706116130<br>Cc1ccc2c(C(=O)N[C@H]3COC[C@H]3NC(=O)[C@H]3OCCO[C@H]3C)n[nH]c2c1<br>$C_{19}H_{24}N_4O_5$<br>388.42 a.m.u. |
| 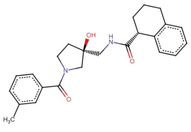<br>PV-006707508242<br>Cc1cccc(C(=O)N2CC[C@](O)(CNC(=O)[C@@H]3CCCc4ccccc43)C2)c1<br>$C_{24}H_{28}N_2O_3$<br>392.49 a.m.u. | 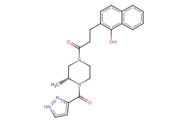<br>PV-006710095262<br>C[C@@H]1CN(C(=O)CCc2ccc3ccccc3c2O)CCN1C(=O)c1cc[nH]n1<br>$C_{22}H_{24}N_4O_3$<br>392.45 a.m.u. | 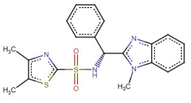<br>Z1316825222<br>Cc1nc(S(=O)(=O)N[C@H](c2ccccc2)c2nc3ccccc3n2C)sc1C<br>$C_{20}H_{20}N_4O_2S_2$<br>412.53 a.m.u. | 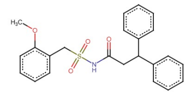<br>Z1679037537<br>COc1ccccc1CS(=O)(=O)NC(=O)CC(c1ccccc1)c1ccccc1<br>$C_{23}H_{23}NO_4S$<br>409.50 a.m.u. | 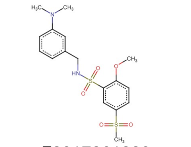<br>Z2017261328<br>COc1ccc(S(C)(=O)=O)cc1S(=O)(=O)NCc1cccc(N(C)C)c1<br>$C_{17}H_{22}N_2O_5S_2$<br>398.50 a.m.u. | 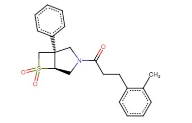<br>Z5052318887<br>Cc1ccccc1CCC(=O)N1C[C@@H]2[C@](c3ccccc3)(C1)CS2(=O)=O<br>$C_{21}H_{23}NO_3S$<br>369.48 a.m.u. |
| 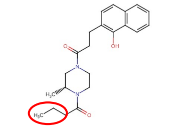<br>Z9530264738<br>CCCC(=O)N1CCN(C[C@@H]1C)C(=O)CCC=2C=CC3=CC=CC=C3C2O<br>$C_{22}H_{28}N_2O_3$<br>368.47 a.m.u. | 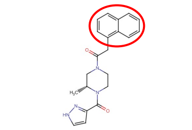<br>Z9530264737<br>C[C@@H]1CN(CCN1C(=O)C=2C=CNN2)C(=O)CC=3C=CC=C4C=CC=CC34<br>$C_{21}H_{22}N_4O_2$<br>362.42 a.m.u. | | | | |

**Supplementary Table 7. Additional ligands for 7L1U.** Ligands selected from the Enamine catalog identified by REAL-M and grouped based on whether they are from the GLU70 (first three molecules) or TYR220 motif-biased selection (20 molecules after double line). Each ligand is depicted along with its corresponding Enamine ID, SMILES string, chemical formula, and mass in atomic mass units.

| | | |
|---|---|---|
| 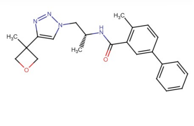<br>PV-006644207828<br>CC=1C=CC(=CC1C(=O)N[C@H](C)CN2C=C(N=N2)C3(C)COC3)C=4C=CC=CC4<br>$C_{23}H_{26}N_4O_2$<br>390.20 a.m.u. | 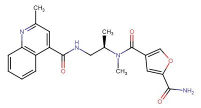<br>PV-006653503749<br>CC=1C=C(C(=O)NC[C@@H](C)N(C)C(=O)C2=COC(=C2)C(N)=O)C=3C=CC=CC3N1<br>$C_{21}H_{22}N_4O_4$<br>394.16 a.m.u. | 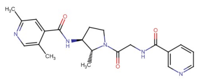<br>PV-006697174573<br>CC1=CN=C(C)C=C1C(=O)N[C@H]2CCN(C(=O)CNC(=O)C=3C=CC=NC3)[C@@H]2C<br>$C_{21}H_{25}N_5O_3$<br>395.20 a.m.u. |

| | | | | | |
|---|---|---|---|---|---|
| 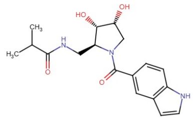<br>PV-003233863128<br>CC(C)C(=O)NC[C@H]1[C@H](O)[C@H](O)CN1C(=O)C=2C=CC=3NC=CC3C2<br>$C_{18}H_{23}N_3O_4$<br>345.17 a.m.u. | 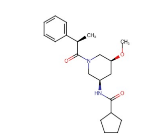<br>PV-003389854278<br>CO[C@H]1C[C@H](CN(C1)C(=O)[C@H](C)C=2C=CC=CC2)NC(=O)C3CCCC3<br>$C_{21}H_{30}N_2O_3$<br>358.23 a.m.u. | 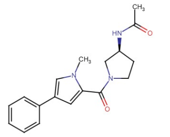<br>PV-003669632782<br>CC(=O)N[C@H]1CCN(C1)C(=O)C2=CC(=CN2C)C=3C=CC=CC3<br>$C_{18}H_{21}N_3O_2$<br>311.16 a.m.u. | 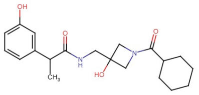<br>PV-003930965565<br>CC(C(=O)NCC1(O)CN(C1)C(=O)C2CCCCC2)C=3C=CC=C(O)C3<br>$C_{20}H_{28}N_2O_4$<br>360.20 a.m.u. | 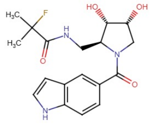<br>PV-003968789845<br>CC(C)(F)C(=O)NC[C@H]1[C@H](O)[C@H](O)CN1C(=O)C=2C=CC=3NC=CC3C2<br>$C_{18}H_{22}FN_3O_4$<br>363.16 a.m.u. | 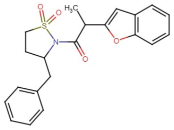<br>PV-004157240875<br>CC(C(=O)N1C(CC=2C=CC=CC2)CCS1(=O)=O)C3=CC=4C=CC=CC4O3<br>$C_{21}H_{21}NO_4S$<br>383.12 a.m.u. |
| 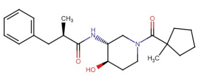<br>PV-004277076181<br>C[C@H](CC=1C=CC=CC1)C(=O)N[C@@H]2CN(CC[C@H]2O)C(=O)C3(C)CCCC3<br>$C_{22}H_{32}N_2O_3$<br>372.24 a.m.u. | 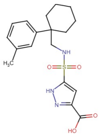<br>PV-005499336810<br>CC=1C=CC=C(C1)C2(CNS(=O)(=O)C3=CC(=NN3)C(=O)O)CCCCC2<br>$C_{18}H_{23}N_3O_4S$<br>377.14 a.m.u. | 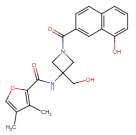<br>PV-006134416639<br>CC1=COC(C(=O)NC2(CO)CN(C2)C(=O)C=3C=CC=4C=CC=C(O)C4C3)=C1C<br>$C_{22}H_{22}N_2O_5$<br>394.15 a.m.u. | 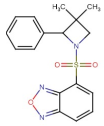<br>Z1438098873<br>CC1(C)CN(C1C=2C=CC=CC2)S(=O)(=O)C3=CC=CC4=NON=C34<br>$C_{17}H_{17}N_3O_3S$<br>343.10 a.m.u. | 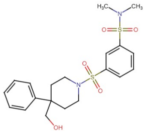<br>Z2212817833<br>CN(C)S(=O)(=O)C=1C=CC=C(C1)S(=O)(=O)N2CCC(CO)(CC2)C=3C=CC=CC3<br>$C_{20}H_{26}N_2O_5S_2$<br>438.13 a.m.u. | 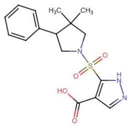<br>Z2437615025<br>CC1(C)CN(CC1C=2C=CC=CC2)S(=O)(=O)C=3NN=CC3C(=O)O<br>$C_{16}H_{19}N_3O_4S$<br>349.11 a.m.u. |
| 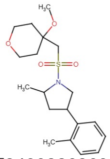<br>Z2480226269<br>COC1(CS(=O)(=O)N2CC(CC2C)C=3C=CC=CC3C)CCOCC1<br>$C_{19}H_{29}NO_4S$<br>367.18 a.m.u. | 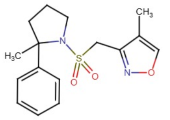<br>Z3474884733<br>CC1=CON=C1CS(=O)(=O)N2CCCC2(C)C=3C=CC=CC3<br>$C_{16}H_{20}N_2O_3S$<br>320.12 a.m.u. | 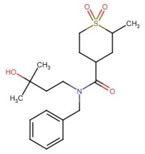<br>Z3583874891<br>CC1CC(CCS1(=O)=O)C(=O)N(CCC(C)(C)O)CC=2C=CC=CC2<br>$C_{19}H_{29}NO_4S$<br>394.16 a.m.u. | 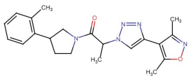<br>PV-005484108139<br>CC1=NOC(C)=C1C2=CN(N=N2)C(C)C(=O)N3CCC(C3)C=4C=CC=CC4C<br>$C_{21}H_{25}N_5O_2$<br>379.20 a.m.u. | 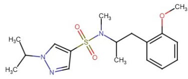<br>Z1271390920<br>COC=1C=CC=CC1CC(C)N(C)S(=O)(=O)C=2C=NN(C2)C(C)C<br>$C_{17}H_{25}N_3O_3S$<br>351.16 a.m.u. | 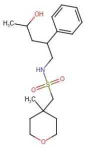<br>Z3408050766<br>CC(O)CC(CNS(=O)(=O)CC1(C)CCOCC1)C=2C=CC=CC2<br>$C_{18}H_{29}NO_4S$<br>355.18 a.m.u. |
| 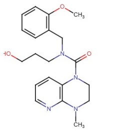<br>Z4126269790<br>COC=1C=CC=CC1CN(CCCO)C(=O)N2CCN(C)C=3N=CC=CC32<br>$C_{20}H_{26}N_4O_3$<br>370.20 a.m.u. | | | | | |

**Supplementary Table 8. Sulfonamide-containing ligands for 7SR8.** Ligands selected from the 2.6B (top set) or 64.9B (bottom set) Enamine library. Each ligand is depicted along with its corresponding Enamine ID, SMILES string, chemical formula, and mass in atomic mass units.

| | | | | | |
|---|---|---|---|---|---|
| 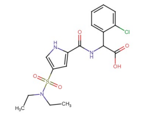<br>Z1603665044<br>CCN(CC)S(=O)(=O)C1=CNC(=C1)C(=O)NC(C(=O)O)C2=CC=CC=C2Cl<br>C17H20ClN3O5S<br>413.08 a.m.u. | 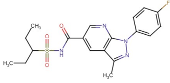<br>Z1616530258<br>CCC(CC)S(=O)(=O)NC(=O)C=1C=NC2=C(C1)C(C)=NN2C=3C=CC(F)=CC3<br>C19H21FN4O3S<br>404.13 a.m.u. | 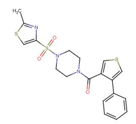<br>Z2172433126<br>CC1=NC(=CS1)S(=O)(=O)N2CCN(CC2)C(=O)C3=CSC=C3C=4C=CC=CC4<br>C19H19N3O3S3<br>433.06 a.m.u. | 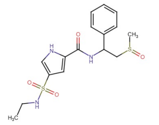<br>Z1319614435<br>CCNS(=O)(=O)C1=CNC(=C1)C(=O)NC(CS(C)=O)C=2C=CC=CC2<br>C16H21N3O4S2<br>383.10 a.m.u. | 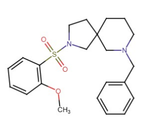<br>Z1324236286<br>COC=1C=CC=CC1S(=O)(=O)N2CCC3(C2)CCCN(CC=4C=CC=CC4)C3<br>C22H28N2O3S<br>400.18 a.m.u. | 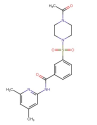<br>Z1481848144<br>CC(=O)N1CCN(CC1)S(=O)(=O)C2=CC=CC(=C2)C(=O)NC=3C=C(C)C=C(C)N3<br>C20H24N4O4S<br>416.15 a.m.u. |
| 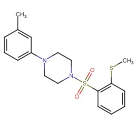<br>Z1765621729<br>CSC=1C=CC=CC1S(=O)(=O)N2CCN(CC2)C3=CC=CC(C)=C3<br>C18H22N2O2S2<br>362.11 a.m.u. | 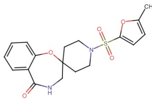<br>Z2919346512<br>CC1=CC=C(O1)S(=O)(=O)N2CCC3(CC2)CNC(=O)C4=CC=CC=C4O3<br>C18H20N2O5S<br>376.11 a.m.u. | 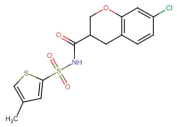<br>Z9704925833<br>CC1=CSC(=C1)S(=O)(=O)NC(=O)C2COC=3C=C(Cl)C=CC3C2<br>C15H14ClNO4S2<br>371.00 a.m.u. | 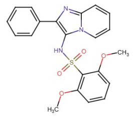<br>Z1729674074<br>COC1=CC=CC(OC)=C1S(=O)(=O)NC2=C(N=C3C=CC=CN23)C=4C=CC=CC4<br>C21H19N3O4S<br>409.11 a.m.u. | 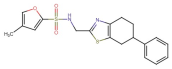<br>Z9704925832<br>CC1=COC(=C1)S(=O)(=O)NCC2=NC=3CCC(CC3S2)C=4C=CC=CC4<br>C19H20N2O3S2<br>388.09 a.m.u. | |

| | | | | | |
|---|---|---|---|---|---|
| 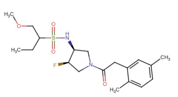<br>Z9862967556<br>CCC(COC)S(=O)(=O)N[C@H]1CN(C[C@H]1F)C(=O)CC2=CC(C)=CC=C2C<br>C19H29FN2O4S<br>400.18 a.m.u. | 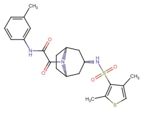<br>Z9862967493<br>CC1=CSC(C)=C1S(=O)(=O)N[C@H]2C[C@@H]3CC[C@H](C2)N3C(=O)C(=O)NC=4C=CC=C(C)C4<br>C22H27N3O4S2<br>461.14 a.m.u. | 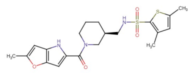<br>Z9862967494<br>CC1=CC=2NC(=CC2O1)C(=O)N3CCC[C@@H](CNS(=O)(=O)C=4SC(C)=CC4C)C3<br>C20H25N3O4S2<br>435.13 a.m.u. | 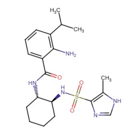<br>Z9862967495<br>CC=1NC=NC1S(=O)(=O)N[C@H]2CCCC[C@@H]2NC(=O)C3=CC=CC(=C3N)C(C)C<br>C20H29N5O3S<br>419.20 a.m.u. | 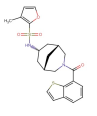<br>Z9862967496<br>CC=1C=COC1S(=O)(=O)N[C@H]2C[C@@H]3C[C@H](C2)CN(C3)C(=O)C=4C=CC=C5C=CSC45<br>C22H24N2O4S2<br>444.12 a.m.u. | 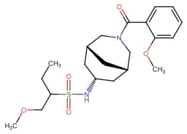<br>Z9862967559<br>CCC(COC)S(=O)(=O)N[C@H]1C[C@@H]2C[C@H](C1)CN(C2)C(=O)C3=CC=CC=C3OC<br>C21H32N2O5S<br>424.20 a.m.u. |
| 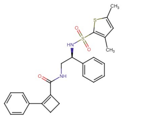<br>Z9862967498<br>CC1=CC(C)=C(S1)S(=O)(=O)N[C@H](CNC(=O)C=2CCC2C=3C=CC=CC3)C=4C=CC=CC4<br>C25H26N2O3S2<br>466.14 a.m.u. | 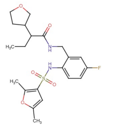<br>Z9862967561<br>CCC(C(=O)NCC1=CC(F)=CC=C1NS(=O)(=O)C=2C=C(C)OC2C)C3CCOC3<br>C21H27FN2O5S<br>438.16 a.m.u. | 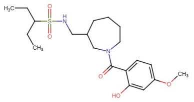<br>Z9862967564<br>CCC(CC)S(=O)(=O)NCC1CCCCN(C1)C(=O)C2=CC=C(C=C2O)OC<br>C20H32N2O5S<br>412.20 a.m.u. | 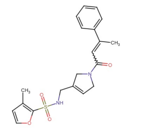<br>Z9862967565<br>CC(=CC(=O)N1CC=C(CNS(=O)(=O)C=2OC=CC2C)C1)C=3C=CC=CC3<br>C20H22N2O4S<br>386.13 a.m.u. | 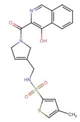<br>Z9902831373<br>CC1=CSC(=C1)S(=O)(=O)NCC2=CCN(C2)C(=O)C=3N=CC=4C=CC=CC4C3O<br>C20H19N3O4S2<br>429.08 a.m.u. | 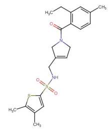<br>Z9862967503<br>CCC=1C=C(C)C=CC1C(=O)N2CC=C(CNS(=O)(=O)C3=CC(C)=C(C)S3)C2<br>C21H26N2O3S2<br>418.14 a.m.u. |
| 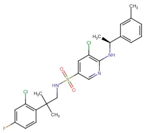<br>Z9862967505<br>CC=1C=CC=C(C1)[C@H](C)NC=2N=CC(=CC2Cl)S(=O)(=O)NCC(C)(C)C=3C=CC(F)=CC3Cl<br>C24H26Cl2FN3O2S<br>509.11 a.m.u. | 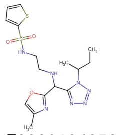<br>Z9908134352<br>CCC(C)N1N=NN=C1C(NCCNS(=O)(=O)C2=CC=CS2)C3=NC(C)=CO3<br>C16H23N7O3S2<br>425.13 a.m.u. | 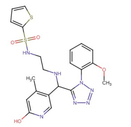<br>Z9862978169<br>COC=1C=CC=CC1N2N=NN=C2C(NCCNS(=O)(=O)C3=CC=CS3)C=4C=NC(O)=CC4C<br>C21H23N7O4S2<br>501.13 a.m.u. | 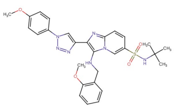<br>Z9862967509<br>COC=1C=CC(=CC1)N2C=C(N=N2)C=3N=C4C=CC(=CN4C3NCC=5C=CC=CC5OC)S(=O)(=O)NC(C)(C)C<br>C28H31N7O4S<br>561.22 a.m.u. | 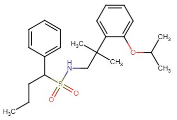<br>Z9862967563<br>CCCC(C=1C=CC=CC1)S(=O)(=O)NCC(C)(C)C2=CC=CC=C2OC(C)C<br>C23H33NO3S<br>403.22 a.m.u. | 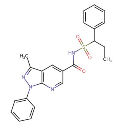<br>Z2209201811<br>CCC(C=1C=CC=CC1)S(=O)(=O)NC(=O)C=2C=NC3=C(C2)C(C)=NN3C=4C=CC=CC4<br>C23H22N4O3S<br>434.14 a.m.u. |
| 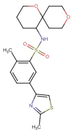<br>Z9882627072<br>CC1=NC(=CS1)C=2C=CC(C)=C(C2)S(=O)(=O)NC3CCCOC43CCOCC4<br>C20H26N2O4S2<br>422.13 a.m.u. | 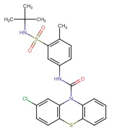<br>Z9882627073<br>CC=1C=CC(=CC1S(=O)(=O)NC(C)(C)C)NC(=O)N2C=3C=CC=CC3SC=4C=CC(Cl)=CC42<br>C24H24ClN3O3S2<br>501.09 a.m.u. | | | | |

**Supplementary Table 9. Genotyping information for zebrafish lines.**

| **Zebrafish Line** | *hcrtr2*[uab480] | *Tg(hsp70l:hcrt), zf12Tg* |
|---|---|---|
| **Ensembl Gene** | ENSDARG00000023722 | ENSDARG00000070932 |
| **Transcript; Exon #** | ENSDART00000038434.8;e1 | NA |
| **ZFIN ID** | ZDB-GENE-070228-3 | ZDB-GENE-040324-1 |
| **Mutation Area - WT DNA Sequence** | GGCGCAAACATGTCCGGGATCTCC GTCCAGCGCGCCTGCAACTCCTGC TTCACCTCAGCCCAGCACCTCAAC TCCTCCACAATCTCGCATTCCCAC GCGGAAAACGAAGACGACGAGCTC CTCAAGTACATCTGTCTCTGGTCG GAAACACACTGGGTAAGTTTTGAC GCGCGCGCGTAACTCACA | NA |
| **Mutation Area - Mutant DNA Sequence** | GGCGCAAACATGTCCGGGATCTCC GTCCAGCGCGCCTGCAACTCCTGC TTCACCTCAGCCCAGCACCTCAAC TCCTCCACAATCTCGCATTCCCAC GCGGAAAACGAAGACGACGAGCTC CTCAAGTACATCTGTCTCTGGTCG GAAACACACTGGGTAAGTTTTGAC GCGCGCGCGTAACTCACA | Transgenic insertion with I-SceI |
| **Guide RNA Target Sites** | ATGCGAGATTGTGTCCGCGGAGG GCTCCTCAAGTACATCTGGAGGG TACGAATGGGTGCTCATAGCTGG TGTTTCCGACCAGAGACACGAGG | NA |
| **Genotyping Primers** | F:GGCGCAAACATGTCCGGG R:tgtgagttacgcgcgcgcgtc | F:CGGGACCACCATGGACT R:GGTTTGTCCAAACTCATCAATGT |
| **WT Genotyping Size** | 262 | No band |
| **MUT Genotyping Size** | 186 | 470 |
